# Genome biology and evolution of mating type loci in four cereal rust fungi

**DOI:** 10.1101/2023.03.02.530769

**Authors:** Zhenyan Luo, Alistair McTaggart, Benjamin Schwessinger

**Affiliations:** Research Biology School, Australian National University, Canberra, ACT, Australia; Centre for Horticultural Science, Queensland Alliance for Agriculture and Food Innovation, The University of Queensland, Ecosciences Precinct, Dutton Park, Queensland, Australia

## Abstract

Obligate heterozygous loci such as sex- or mating-compatibility regions often display suppression of recombination and signals of genomic degeneration. In Basidiomycota, two distinct gene loci confer mating compatibility. These encode for homeodomain (*HD*) transcription factors and pheromone receptor (*Pra*)-ligand pairs. To date genome level mating type (*MAT*) loci analysis is lacking for obligate biotrophic basidiomycetes in the order *Pucciniales*, which contains many economically important plant pathogens.

Here, we focus on four *Puccinia* cereal rust species, including *P. coronata* f. sp. *avenae*, *Puccinia graminis* f. sp. *tritici*, *P. triticina* and *P. striiformis* f. sp. *tritici*, which infect oat and wheat. *MAT* loci are located on two separate chromosomes supporting previous hypotheses of tetrapolar mating types in the *Pucciniales*. The *HD* locus is multiallelic in all four species while the *PR* locus appears to be biallelic except for *P. graminis* f. sp. *tritici* which displays genetic features of more than two alleles. *HD* loci were largely conserved in their macrosynteny within and between species without strong signals of recombination suppression. *PR* loci proximate regions, however, displayed extensive signs of recombination suppression and genomic degeneration in the three species with a clear biallelic *PR* locus. These observations suggest a correlation between recombination suppression, genomic degeneration and allele status of *MAT* loci which is consistent with recent mathematical modelling and simulations. Finally, we confirm the evolutionary conservation of *MAT* gene expression during the asexual infection cycle of the cereal host which we propose is related to correct nuclear pairing during spore formation. Together, our study provides insights into the evolution of *MAT* loci of key pathogenic *Puccinia* species. This detailed understanding is important to predict possible combinations of nuclear pairs that can arise via sexual reproduction or somatic recombination to enable the evolution of newly virulent isolates of these important plant pathogens.

**Author summary:** Sex of animals and some plants is determined by sex chromosomes. In fungi, mate compatibility is determined by mating type (*MAT*) loci, which share some features with sex chromosomes including recombination suppression around heterozygous loci. Here, we study the *MAT* loci in fungal pathogens from the order *Pucciniales* that can cause rust diseases on many economically important plants including wheat and oat. We show that one of the *MAT* loci is multiallelic, while the other is biallelic. The biallelic locus shows strong signs of recombination suppression and genetic deterioration with an increase in transposable elements and gene deserts surrounding the locus. Our findings on the genome biology of *MAT* loci in four economically important pathogens will lead to a better understanding and prediction of evolution of novel virulent isolates that can lead to large scale pandemics in agriculture.

## Introduction

The evolution of ‘sex and mating-type’ determining loci and chromosomes is a fundamental question in biology. It is widely accepted that X and Y chromosomes were originally homologous, but recombination suppression gradually resulted in genetic degeneration of the Y chromosome (1). Genetic degeneration footprints are alleviated rates of (non-)synonymous substitutions (d_N_, d_S_), accumulation of transposable elements (TEs), accumulation of inversions, reduced gene expression and/or reduced gene numbers which are all a consequence of recombination cessation (2-5). In contrast to many animals and plants, antagonistic selection is unlikely to be the evolutionary mechanism that causes recombination cessation in fungal mating-type (*MAT*) loci (6-8). Instead, mathematical modelling and stochastic simulations suggest that recombination suppressors, e.g. inversions or TEs, which are loaded with less deleterious mutations than average initially, are more likely to be fixed at permanently heterozygous *MAT* loci when compared to other autosomal loci. This initial fixation of recombination suppressors at *MAT* loci can lead to a progressive extension of non-recombining regions via the accumulation of additional inversion or TEs over longer evolutionary timeframes (2). Importantly, this model predicts non-recombining regions around *MAT* loci are larger at biallelic loci and in species with smaller effective population size, short haploid phase, outcrossing mating systems, high mutation rates, and extended dikaryotic life stages. Indeed, several fungal species carry extensive non-recombining regions around *MAT* loci. Recombination suppression can range from hundreds of kbp (kilo base pairs) to several mbp (mega base pairs) and lead to genetic degeneration (5, 9), such as in *Neurospora tetrasperma* (10), *Podospora anserina* (11), *Schizothecium tetrasporum* (12) and several *Microbotryum* spp. (8, 13).

Detailed analyses of *MAT* loci and their genome biology has been lacking behind for rust fungi of the order *Pucciniales* in the division Basidiomycota (9). This is despite the fact that *Pucciniales* is the fungal order with the most known plant pathogens with close to 8,000 described species that cause diseases with significant environmental and economic impact for various trees and crops such as poplar, paperbarks, wheat, oat, soybean, and coffee (14-16). Overall, Basidiomycota have evolved unique mating systems to govern nuclear compatibility, mate selection, and completion of the full life cycle (9). From a genetic perspective, two *MAT* loci determine mating-type identity and non-self-recognition in most Basidiomycota. The pheromone receptor (*PR*) locus contains a pheromone receptor (*Pra*) gene and at least one mating pheromone peptide precursor gene (*mfa*) (17). The *PR* locus defines pre-mating compatibility and gamete fusion. The PRA protein is a transmembrane localized G protein-coupled receptor, which recognizes the processed and posttranslationally modified mature pheromone peptide MFA encoded by a matching allelic *PR* locus (9). Downstream of initial gamete fusion, the homeodomain (*HD*) locus determines success in postmating development (9). The *HD* locus contains two tightly linked homeodomain transcription factors genes, (*bW-HD1* and *bE-HD2* in *Pucciniales* (18, 19), also known as *bE-HD1* and *bW-HD2* in other fungi (9)) which share a promoter and are outwardly transcribed in opposite directions. Their protein products need to be of different allelic specificity to form heterocomplexes and to initiate a transcriptional cascade downstream of PRA-MFA activation (9). The *HD* transcription factors regulate cellular development during mating, maintenance of the dikaryotic state, and control pathogenicity in some plant pathogens such as smuts (13). The *HD* and *PR* loci can be either physically linked or unlinked which means they segregate together or independently, respectively. The mating system in Basidiomycota influences the genomic organization and resulting segregation patterns of *MAT* loci. Most basidiomycete species are heterothallic, where sexual gametes must carry different alleles at both *MAT* loci and intra-haploid mating is not possible. Hence, heterothallic species can either have bipolar or tetrapolar mating systems. In bipolar species the *HD* and *PR* loci are physically linked or one locus has lost its function in mating. In any case, in bipolar species only one *MAT* locus determines mate compatibility. In tetrapolar species, the *HD* and *PR* loci are unlinked and both determine mate compatibility. Bipolar mating systems are favorable in inbreeding populations that undergo selfing. Under selfing a single locus is advantageous as the likelihood of compatibility amongst gametes derived from the same diploid individual is 50% (13). In comparison, a tetrapolar mating system has at most a 25% compatibility under the same selfing conditions. In contrast, outcrossing mating systems benefit from multiallelism at *MAT* loci because it increases compatibility at syngamy of haploid gametes from different diploid individuals (6, 9, 13). In many tetrapolar basidiomycete species, the *HD* locus is highly polymorphic in the population with ten to hundred known or estimated alleles that are under negative, frequency dependent selection (13, 20, 21). The tetrapolar mating system is thought to be the ancestral state in Basidiomycota, yet several species have evolved bipolarity independently. For example, *Microbotryum* spp. are highly selfing, and multiple independent evolutionary events have linked the *HD* locus and the *PR* locus into evolutionary strata of different sizes and different ages in this genus (3, 7, 8, 22, 23). These evolutionary strata show clear signatures of recombination suppression and genetic degeneration. Similarly, in the Ustilaginomycotina, several species evolved bipolarity linking both *MAT* loci such as *Ustilago hordei, U. bromivora* or *Sporisorium scitamineum* (9). In contrast, a recent study showed that tetrapolarity has evolved multiple times in the human skin fungus genus *Malassezia,* where the ancestral state is pseudobipolar (24).

Studying mating systems in fungal species of the order *Pucciniales* has been difficult because of their obligate biotrophy and often complex life cycles that prevent culturing outside of their hosts and often requires multiple hosts to complete their sexual life cycle (9). For example, macrocyclic and heteroecious rust fungi have five spore stages and require two hosts to complete their asexual and sexual life cycles. During the asexual infection cycle on the “primary” host, e.g. wheat and oat, rust fungi are dikaryotic and produce re-infective urediniospores that harbor two distinct nuclei carrying compatible mating types. The sexual cycle is initiated with the production of teliospores, which represent the short diploid phase of rust fungi. Resulting haploid basidiospores infect the “alternate” host, e. g. *Berberis sp*., and form haploid infection structures with distinct mating types. Fusion of cell with compatible mating types generates dikaryotic intercellular mycelia within the plant tissue that gives rise to dikaryotic aeciospores which are only infective on the “primary” host (25). Infection of the primary host completes the full life cycle (9, 26). The role of *MAT* genes during the life cycle of rust fungi is rudimentary. Though it is hypothesized that *MAT* genes regulate appropriate nuclear paring during dikaryotic spore production and mediate the compatibility of haploid cell fusions on the “alternate” host (9, 26). Original research suggested that most rust fungal species are bipolar (26), yet more recent studies on the rust fungi *Melampsora lini* and *P. coronata* var. *coronata* revealed tetrapolar mating systems (27, 28). In the absence of direct experimental evidence, genomic insights into *MAT* loci from genome assemblies of the dikaryotic state can provide hypotheses on the breeding systems. Independent initial genome analyses of several rust fungi revealed two unlinked *MAT* loci with *PR* and *HD* loci being located on different contigs, scaffolds or chromosomes including in *Austropuccinia psidii* (29)*, P. graminis* f. sp. *tritici* (30) and *P. tritici* (18, 19). This genome organization supports the hypothesis of a tetrapolar mating system in these rust fungi. These rust fungi encode *Pra* receptor genes belonging to the STE3 family. The likely *bona fide* receptors are encoded by *STE3.2-2* and *STE3.2-3* that display clear signatures of ancient trans-specific polymorphisms which are likely ancestral to Basidiomycota (31). Here the sequences of a given mating type is more similar to pheromone receptor alleles of the same mating type in distantly related fungi than the alternate allele in the same species. In most cases, at least one *mfa* allele has been found in proximity to the *STE3.2-2/3* locus. The additional STE3 family member *STE3.2-1* is nearly homozygous in studied species such as *P. tritici* and is hypothesized not to be involved in mating (18). To date, the *PR* locus has been found to be biallelic in available population level datasets for *P. tritici* and *P. striiformis* f. sp. *tritici* (18, 21, 32), yet *Melampsora larici-populina* might be carrying additional alleles and population level analyses are missing for other rust fungi. In contrast, the *HD* locus is multiallelic in *P. triticina* and *P. striiformis* f. sp. *tritici* (18, 21, 32) encoding variants of *bW-HD1* and *bE-HD2* that are highly dissimilar only in the variable N-terminal domain, shown to be essential for functional heterodimerization in other Basidiomycota (33, 34). It is noteworthy that direct biochemical or genetic evidence is lacking for any of these *MAT* genes’ involvement in mating in rust fungi. Yet, complementation assays in *U. maydis* and host induced gene silencing on wheat demonstrated the importance of *Pra* and *HD* genes of *P. triticina* for mating in a heterologous system and for spore production during asexual infection (18). In addition, *MAT* genes were expressed during the asexual and/or sexual infection cycle in several rust fungal species (18, 35). These initial studies open up several outstanding knowledge gaps for *MAT* loci in rust fungi: 1) What is the chromosomal organization of *PR* and *HD* loci and how does it compare between closely related species? 2) What is the allelic diversity of *MAT* genes within rust fungal species? 3) What is the composition and organization of *MAT* loci proximal regions, how does it compare between *PR* and *HD* loci, and how does it vary between closely related species? 4) What is the compositional and organizational variation of *MAT* loci within rust fungal species? 5) Is the expression of *MAT* genes during the asexual infection cycle common in rust fungi?

Addressing these knowledge gaps has long been a challenge because initial genome assemblies of rust fungi were fragmented and the structure of the *MAT* loci could not be assessed consistently. For example, clustering of repetitive elements around *MAT* loci (17, 36, 37) is a challenge for studying *MAT* loci in rust fungi with repeat-rich genomes because repetitive sequences cause gaps, fragmentation, and phase-errors in the assembled genome. Here, we overcome this challenge and address important knowledge gaps around *MAT* loci in four cereal rust fungi by providing a detailed comparative analysis of publicly available chromosome scale phased genome assemblies and complementary Illumina short-read datasets. The study species include the oat crown rust fungus *P. coronata* f. sp. *avenae* (38-40), and the three wheat rust fungi, *P. graminis* f. sp. *tritici* (30, 41, 42), *P. triticina* (43-46) and *P. striiformis* f. sp. *tritici* (47-53), which combined are the biggest threat to wheat production globally causing losses of several billion dollars every year (16). Our detailed comparative analyses set out to address the following specific objectives. 1) To determine the chromosomal organization of *MAT* loci. 2) To assess the allelic diversity of these loci in the four cereal rust fungi. 3) To evaluate synteny, recombination suppression, and genomic deterioration around *PR* and *HD* loci with the prediction that multiallelic loci are more syntenic than biallelic loci and that the latter show stronger footprints of recombination suppression and genomic degeneration. 4) To gage the conservation and/or plasticity of *MAT* loci within a cereal rust fungus species using *P. triticina* as model system. And 5) To determine the expression of *Pra* and *HD* genes during the asexual infection cycle of the cereal hosts.

We show that *HD* genes are multiallelic in all four rust fungal species and *Pra* genes are unequivocal biallelic in three out of four. Only biallelic *MAT* loci show strong signs of recombination suppression and genomic degeneration which confirms with recent mathematical modelling. Our results provide novel insights into the genome biology of *MAT* loci in cereal rust fungi. These are important to consider when making predictions about mating and hybridization events that are possibly linked to the evolution of novel pathogenicity traits.

## Results

### Genomic organisation and inheritance of mating type genes suggest cereal rust fungi have tetrapolar mating systems

We set out to test previous preliminary observations that suggest cereal rust fungi have a tetrapolar mating system with *PR* and *HD* loci being unlinked (18, 19). We made use of the seven available chromosome-level genome assemblies of four cereal rust fungi *P. coronata* f. sp. *avenae* (*Pca*) (38, 40, 54), *P. graminis* f. sp. *tritici* (*Pgt*) (30, 41, 42, 53), *P. triticina* (*Pt*) (32, 43-46, 55) and *P. striiformis* f. sp. *tritici* (*Pst*) (47, 49-52, 56), in addition to five partially phased assemblies (Table 1). Our initial analysis focused on one reference genome per species with *Pca 203*, *Pgt 21-0*, *Pt 76*, and *Pst 134E*.

**Table 1.** Cereal Rust Fungi Genomes used in the present study. The table provides information on all genome assemblies used in this study, their total dikaryotic genome assembly size (“Genome size”), and completeness as assessed with BUSCO (“BUSCO (%)”). For the “BUSCO (%)” column the first number presents the percentage of complete single BUSCO genes identified. The numbers in the brackets provide the percentage of single copy and of duplicated BUSCO genes in the dikaryotic genome assembly. Additional presented metadata includes species name (“Species”), strain name (“Strain”), source (“Source”), download date in DD/MM/YYYY (“Download date”) and the initial reference (“Citation”). The * genome denotes dikaryotic genome assemblies which are chromosome scale phased genome. NCBI – National Center for Biotechnology information, JGI – Joint Genome Institute, CSIRO – Commonwealth Scientific and Industrial Research Organization.

| Species | Strain | Genome size | BUSCO (%) | Source | Download date | Citation |
| --- | --- | --- | --- | --- | --- | --- |
| *Puccinia graminis* f. sp. *tritici* | *Ug99* | 176.2 Mb | 93.4 [S:7.7, D:85.7] | NCBI | 26/05/2021 | Li et al. 2019 |
|  | *21-0\** | 176.9 Mb | 94.2 [S:8.3, D:85.9] | NCBI | 26/05/2021 |  |
| *Puccinia striiformis* f. sp. *tritici* | *DK0911* | 157.8 Mb | 91.1 [S:38.4, D:52.7] | JGI | 1/06/2021 | Schwessinger et al. 2020 |
|  | *104E* | 126.5 Mb | 92.1 [S:7.9, D:84.2] | JGI | 26/05/2021 | Schwessinger et al. 2018 |
|  | *134E\** | 167.7 Mb | 92.6 [S:6.8, D:85.8] | Provided by the authors | 21/06/2021 | Schwessinger et al. 2022 |
| *Puccinia coronata* f. sp. *avenae* | *12NC29* | 105.2 Mb | 89.9 [S:18.1, D:71.8] | NCBI | 26/05/2021 | Miller et al. 2018 |
|  | *12SD80* | 99.2 Mb | 90.2 [S:33.1, D:57.1] | NCBI | 26/05/2021 | Miller et al. 2018 |
|  | *203\** | 208.1 Mb | 92.1 [S:6.9, D:85.2] | CSIRO data access portal | 20/06/2022 | Henningesen et al. 2022 |
| *Puccinia triticina* | *76\** | 253.5 Mb | 91.0 [S:1.7, D:89.3] | Provided by the authors | 28/05/2021 | Duan et al. 2022 |
|  | *15\** | 243.9 Mb | 91.1 [S:1.6, D:89.5] | NCBI | 02/11/2023 | Li et al. 2022 |
|  | *19NSW04\** | 253 Mb | 91.2[S:0.9, D:90.3] | NCBI | 10/10/2023 | Sperschneider et al. 2023 |
|  | *20QLD87\** | 248.1 Mb | 91.3[S:1.3, D:90.0] | NCBI | 10/10/2023 | Sperschneider et al. 2023 |
| *Puccinia polysora* f. sp. *zea* | *GD1913\** | 1.7 Gb | 91.1 [S:1.4, D:89.7] | NCBI | 07/14/2023 | Liang et al. 2023 |

We used previously characterized *MAT* genes from the *Pt* isolate *BBBD* (Table 2) (18) as queries to identify orthologs in available genomes and proteomes (Table 1). We identified *HD* (*bW-HD1* and *bE-HD2*) and *Pra* (*STE3.2-2* and *STE3.2-3*) alleles on chromosome 4 and chromosome 9 respectively, whereas *STE3.2-1* alleles were located on chromosome 1 (S1 Fig). This is consistent with previous preliminary analysis for *Pgt* and *Pt,* which suggested *HD* and *PR* loci are located on two distinct chromosomes (30, 32). We confirmed these results using five genome resources for cereal rust fungi that are only partially phased and non-chromosome scale (Table 1). We identified two alleles of *bW-HD1, bE-HD2* and *PRA* (*STE3.2-2* or *STE3.2-3)* in all five genome assemblies with the exception for *STE3.2-2* in *Pst 104E* and *Pca 12SD80* (38, 48). In the case of *Pst 104E, STE3.2-2* was absent from the genome assembly but we confirmed its presence in the raw sequencing data by mapping sequencing reads against the *Pst 134E* phased chromosome scale genome assembly (47). *STE3.2-2* displayed average genome-wide read coverage without any nucleotide variation confirming that *Pst 104E* was also heterozygous for *Pra*. In the case of *Pca 12SD80,* we identified two copies of *STE3.2-2* on contig 000183F and 000183F_004. This miss-characterization of *STE3.2-2* in *Pst 104E* and *Pca 12SD80* is likely caused by early long-read genome assembly errors using low accuracy PacBio long reads and suboptimal genome assembly algorithms that struggle with highly repetitive regions. Similar observations have been made for missing *Pra* genes reported for earlier genome versions of *Melampsora larici-populina* (57).

**Table 2.**
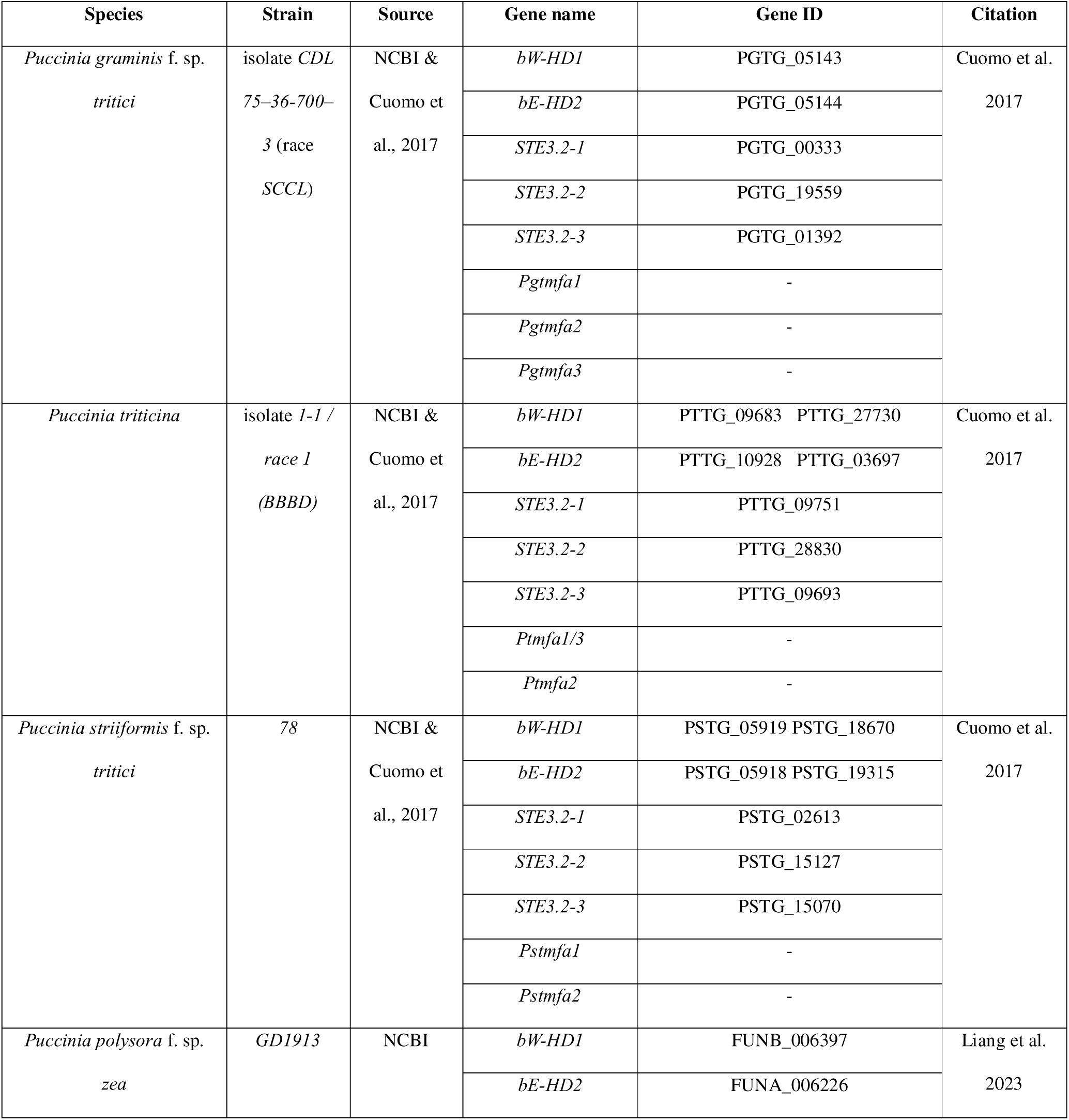
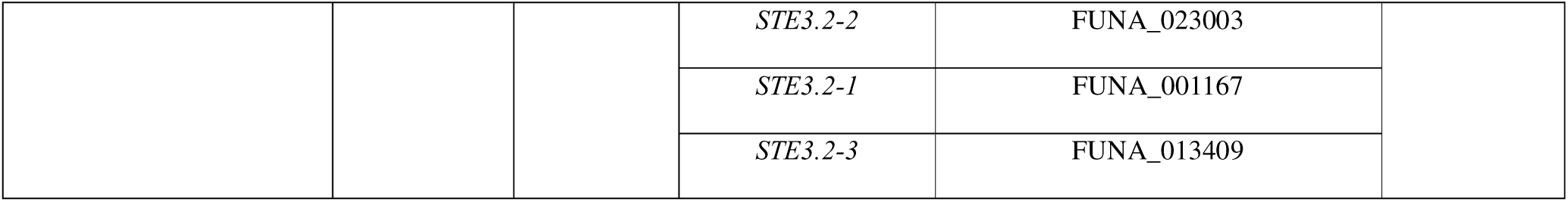
MAT reference genes used as query and outgroups. The table provides gene names (“Gene name”) and gene identifier (“Gene ID”) of *MAT* genes from *Puccinia triticina* initially used as query sequences and from *Puccinia polysora* f. sp. *zeae* used as outgroups. Additional presented metadata includes species name (“Species”), strain name (“Strain”), source (“Source”), and the initial reference (“Citation”). NCBI – National Center for Biotechnology information.

We also identified an additional *Pra*-like gene, *STE3.2-1,* on chromosome 1. However, its two alleles are nearly identical in all dikaryotic genome assemblies and are not associated with any pheromone peptide precursor genes. Hence, *STE3.2-1* is likely not involved mating compatibility in rust fungi.

Taken together, the two *MAT* loci, *HD* and *PR*, are unlinked and heterozygous in all twelve dikaryotic genome assemblies, which is consistent with a genome informed hypothesis of a tetrapolar mating system for cereal rust fungi.

### *Mfa* pheromone peptide precursor genes are closely linked to *STE3.2-2* but not *STE3.2-3*

We searched the available dikaryotic genomes for putative pheromone peptide precursor genes, which encode mating factor a (MFA) peptides. *Mfas* are often linked to *Pra* alleles and predicted to bind to the compatible pheromone receptors encoded at the allelic *PR* locus. We used the previously identified *mfa*1, *mfa2,* and *mfa3* genes of *Pst*, *Pgt,* and *Pt* as queries (18). We identified a single *mfa2* pheromone peptide precursor gene in all species. In all cases, *mfa2* and *STE3.2-2* were closely linked, located within 500–1100 bp from each other, and encoded on the same DNA strand (S2 Fig). The *mfa2* derived peptides were all 34 amino acids long with a characteristic CAAX motif at the C-terminus, where C is cysteine, A is an aliphatic amino acid, and X is any amino acid (58) (S3B Fig). The *MFA2* amino acid sequences were 100% identical at the species level (S3B Fig).

In contrast, the sequence, length, number, and location of *mfa* genes associated with *STE3.2-3* varied between species (S2, S3A and S3C Figs). *Pca* and *Pst* encoded a single *mfa1* allele. *Pcamfa1* of *Pca 203* encoded a 66 amino acid peptide and was located 0.54 mbp upstream of *STE3.2-3*. In *Pst*, *Pstmfa1* encoded a 64 amino acid peptide and was located 10 kbp and 13 kbp away from *STE3.2-3* in *Pst 134E* and *Pst DK0911*, respectively. In contrast, *Pt* carried two identical copies of *Ptmfa1/3* encoding for 61 amino acid peptides. In the *Pt 76* reference, *Ptmfa1/3* were associated with *STE3.2-3* at a distance of 0.24 mbp and 0.27 mbp, respectively (S2 Fig). In addition, *Pgt* encoded for two *mfa* genes in close proximity to *STE3.2-3* that were located upstream and downstream at a distance of 30 kbp and 96 kbp, respectively (S2 Fig). Yet in contrast to *Pt*, *Pgt mfa1* and *mfa3* encoded for highly distinct pheromone peptide precursor that were only 26.47% identical at the amino acid level and varied in length (59 vs 57 amino acids, respectively). Based on their amino acid differences, MFA1 and MFA3 were predicted to be processed into distinct mature MFA peptides that might have different receptor specificities.

In all four cereal rust fungi, *Pra* alleles were associated with species specific *mfa* genes. Yet, the number, distance, organization, and sequence of *mfa* genes varied between species.

### Genealogies elucidate distinct evolutionary histories for the different *MAT* genes

Having identified the *MAT* genes, we investigated the genealogical relationships of each individual gene and compared them between each other and with the species tree. This allowed us to test for shared or distinct evolutionary histories, which can provide indications for recombination within and between loci. For example, so called trans-specific polymorphism indicate that recombination cessation is older than speciation (31, 59). In the absence of a robust species tree (60), we first generated a multilocus species tree based on a multiple sequence alignment of 2284 single orthologs proteins (61) including *P. polysora* f. sp. *zeae* as an outgroup (Fig 1A). The species tree identified the three wheat rust fungi being most closely related to each other when compared to the oat rust fungus *Pca* (Fig 1A). In the wheat rust clade, *Pgt* was closer related to *Pt, w*hile *Pst* was more distantly related to the two. We aligned *HD* genes *bW-HD1* and *bE-HD2* separately to construct independent gene trees using *P. polysora* f.sp. *zeae* alleles as outgroups (Table 2, Figs 1B and 1C). The gene trees of *bW-HD1* and *bE-HD2* group alleles from each species into a species-specific clade (Figs 1B and 1C) and we therefore did not find any evidence for trans-specific polymorphisms for either *HD* gene. We identified multiple *bW-HD1* and *bE-HD2* alleles for the four cereal rust fungi suggesting the *HD* locus is multiallelic. We observed several shared *HD* alleles between isolates in the case of *Pt* while *HD* alleles in *Pca* and *Pgt* appeared to be more diverse. In the case of *Pgt*, we were limited to only four alleles, making any meaningful conclusion difficult. We next tested our null hypothesis that *bW-HD1* and *bE-HD2* have similar evolutionary histories because they are closely linked and share a promoter. We performed approximately unbiased (AU) tests (62) within each species to investigate if the tree topologies of *bW-HD1* and *bE-HD2* are concurrent with each other. In *Pca, Pst,* and *Pt*, we found that the *p*_AU-_value of the AU test was smaller than 0.05 which led us to reject the null hypothesis of concurrent tree topologies. This suggests distinct evolutionary histories for the *bW-HD1* and *bE-HD2* genes in these species, which might be caused by recombination within the *HD* locus. In contrast, the AU-test for *Pgt* suggested similar tree topologies for *bW-HD1* and *bE-HD2*, which likely is influenced by the low sample number of four alleles.

**Fig 1.**
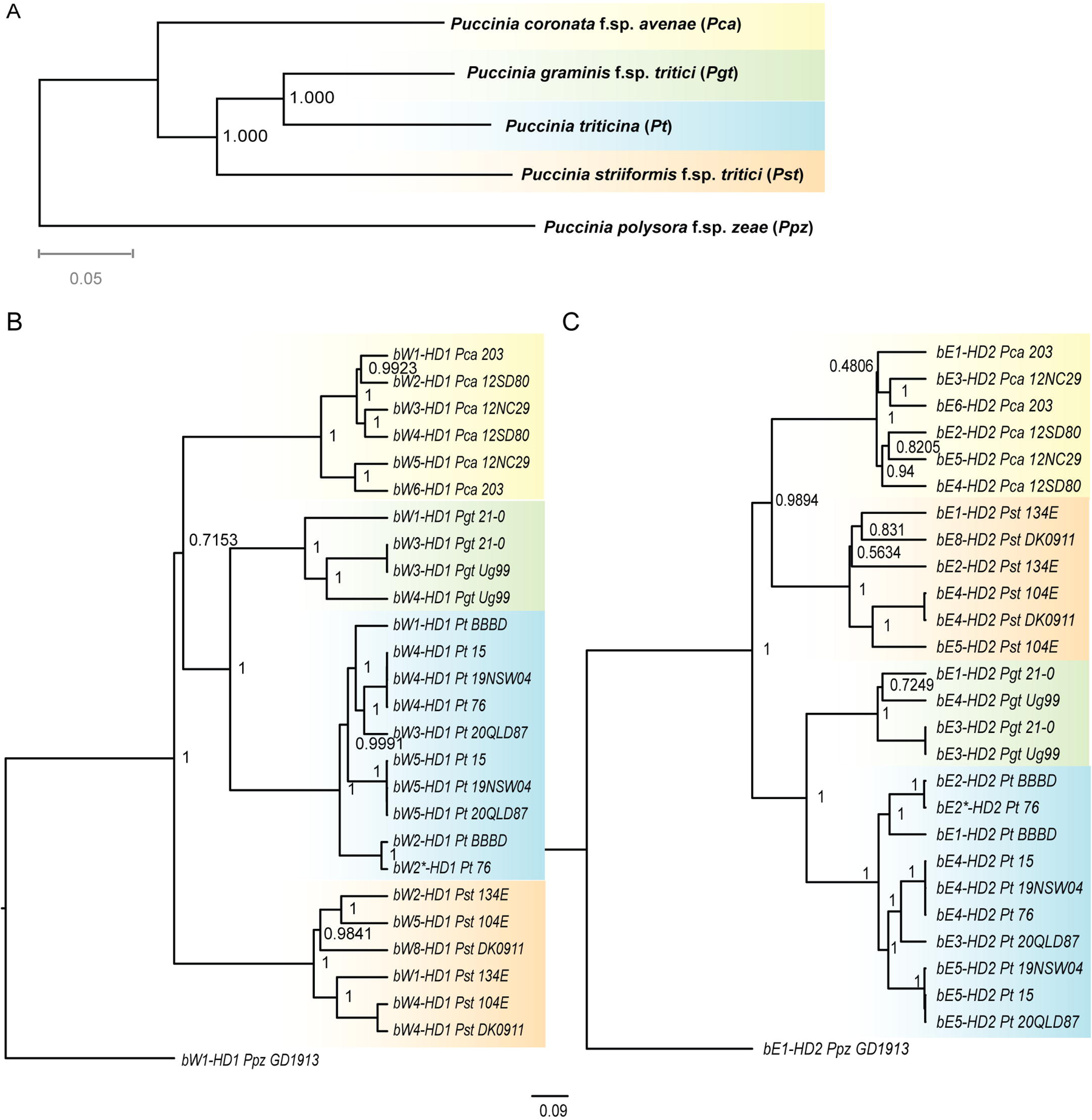
*HD* genealogies suggest distinct evolutionary histories for *bW-HD1* and *bE-HD2* in four cereal rust fungi. (A) Species tree of *Puccinia polysora* f. sp. *zeae* (*Ppz*), *P. coronata* f. sp. *avenae* (*Pca*), *P. graminis* f. sp. *tritici* (*Pgt*), *P. triticina* (*Pt*) and *P. striiformis* f. sp. *tritici* (*Pst*) inferred from 2284 single-copy orthogroups. Numbers on nodes represent local support value computed by Fasttree. *Puccinia polysora* f. sp. *zeae* (*Ppz*) has been used for rooting the species tree. (B) and (C) Bayesian rooted gene tree built from *bW-HD1* or *bE-HD2* coding-based sequence alignment, respectively. Trees are based on a HKY + G model of molecular evolution. In either case, alleles from the same species are grouped into the same clade. Each node is labelled with its values of posterior probability (PP). PP values above 0.95 are considered to have strong evidence for monophyly of a clade and PP values of identical alleles are not displayed. The scale bar represents the number of nucleotide substitutions per site. * marks an allele with minor variations only outside the variable domain, which means this allele is predicted to be functionally equivalent to its closest neighbor. Alleles of the same species are colored with identical background: *Pca* (yellow), *Pgt* (green), *Pt* (blue), *Pst* (orange).

Next, we explored the evolutionary relationship between *Pra* alleles by building a single gene tree using *P. polysora* f.sp. *zeae STE3.2-2* and *STE3.2-3* as outgroups. The *Pra* gene tree formed two obvious clades based on allele identity grouping the S*TE3.2-2* alleles of all species into one clade and all *STE3.2-3* alleles into another clade including respective alleles of *P. polysora* f.sp. *zeae* (Fig 2). In each clade, there was little to no intra-species variation for *STE3.2-2* and *STE3.2-3* based on the minimal branch lengths that separated allele copies in each species sub-clade. This indicates that *Pra* might be biallelic in the cereal rust fungi (see also below). The clear grouping by allele identity indicates strong trans-polymorphisms and a long-term suppression of recombination at the *PR* locus that predates ancestral species split.

**Fig 2.**
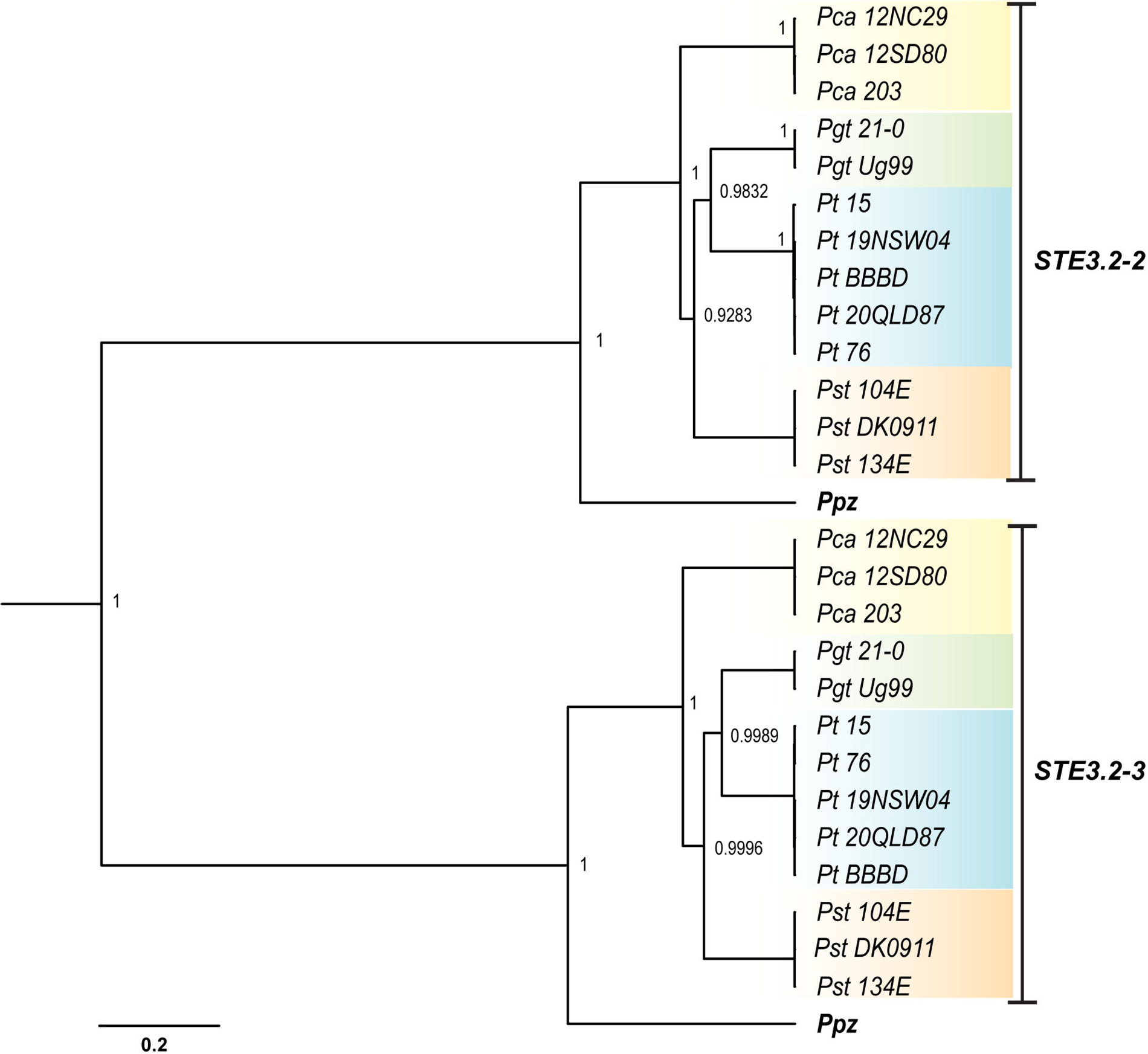
*Pra* genealogy displays ancient trans-specific polymorphism in cereal rust fungi. Bayesian gene tree built from *Pra (STE3.3-2 and STE3.3-3)* coding-based sequence alignment. The tree is based on a TN93 + I model of molecular evolution. The *Pra* alleles, *STE3.2-2* and *STE3.2-3,* were grouped into two clades by allele identity and not species identity. Each node is labelled with its values of posterior probability (PP). PP values above 0.95 are considered as strong evidence for monophyly of a clade and PP values of identical alleles are not displayed. The scale bar represents the number of nucleotide substitutions per site. Species-specific background coloring is the same as for Fig 1.

We next investigated if the putative pheromone peptide precursor *mfa* genes followed similar evolutionary histories compared to their collocated *Pra* genes. Like the *Pra* genealogy, the gene tree of *mfa* alleles revealed strong trans-specific polymorphisms because *mfa* copies clustered by allelic and not species identity (S4 Fig). For example, all *Mfa2* alleles grouped into one large clade according to their physical association with the specific *Pra* allele *STE3.2-2*. Similarly, all *Mfa1* alleles grouped together with similar topology as their linked *STE3.3-2* copies. The species specific *Pgtmfa3s*, which are also physically linked to *STE3.2-3,* were placed as a sister group to the *mfa1* clade, being more closely related to *mfa1* than to *mfa2*.

We also investigated the genealogy of the *Pra-*like *STE3.2-1*. We used *P*. *polysora* f. sp. *zeae STE3.2-1* as outgroup for rooting the gene tree (63). The two *STE3.2-1* alleles identified in each isolate were almost completely identical and grouped by species identity. Overall, *STE3.2-1* showed very little variation within each species (S5 Fig). This further supports our conclusion that *STE3.2-1* is likely not involved in defining mate compatibility.

### *HD* loci display limited signs of genomic deterioration and their synteny is conserved within species

Different effects on recombination have been reported around *MAT* loci in basidiomycetes (8, 64). Hence, we investigated if we could identify signals of altered recombination, synteny, and genomic deterioration around the *MAT* loci in rust fungi. First, we analyzed whether there was a reduction in synteny on chromosome 4 surrounding the *HD* locus as a signature for long-term recombination suppression. We used mummer (65) to align the haplotypes of chromosome 4 from *Pca, Pgt*, *Pt,* and *Pst,* against each other (S6 Fig). Overall macro-synteny of chromosome 4 haplotypes was conserved in all cereal rust fungi with *Pt 76* being the most syntenic (S6C Fig). Macro-synteny in a 40 kb-sized window around the *HD* locus was mostly conserved in *Pt 76*, *Pca 203*, and *Pst 134E* while this initial analysis suggested that this locus was less syntenic in *Pgt 21-0* (S6 Fig).

We investigated the individual allelic divergence along chromosome 4 using synonymous divergence (d_S_) value as an additional measure of footprint of ancient recombination suppression events (8). This analysis tested if the *HD* and their proximal genes showed an increase in d_S_ values, which could be caused by long-term suppression of recombination and accumulation of independent mutations, as seen in other Basidiomycota (13). We calculated the pairwise d_S_ values for all genes across all sister chromosomes in all four species of *Puccinia*. This whole genome analysis of all allele pairs on sister chromosomes suggested that chromosome 4 did not have increased d_S_ values when compared to all other chromosomes (S7 Fig). The d_S_ value of alleles for both *HD* genes fell within the upper 95% quantile of d_S_ values on chromosome 4. Yet, d_S_ values of directly adjacent genes were not elevated beyond the background level when we plotted d_S_ values for allele pairs along chromosome 4 (Fig 3). Consistently, regions around *HD* loci did not show any obviously aberrant patterns of gene or transposon density, which are common features of genetic deterioration (Fig 3). We tested whether *HD* loci were potentially linked to centromeres as reported in other Basidiomycota (13, 66). We used previously identified centromere locations for *Pgt* and *Pst* (47, 67) and identified centromere locations for *Pca* and *Pt* using genome-wide Hi-C heatmaps based on the bowtie-like Hi-C interaction features known for fungal centromeres (S8-11 Figs) (68). This analysis revealed that *HD* loci are likely not directly linked to centromeres in any of the four cereal rust fungi.

**Fig 3.**
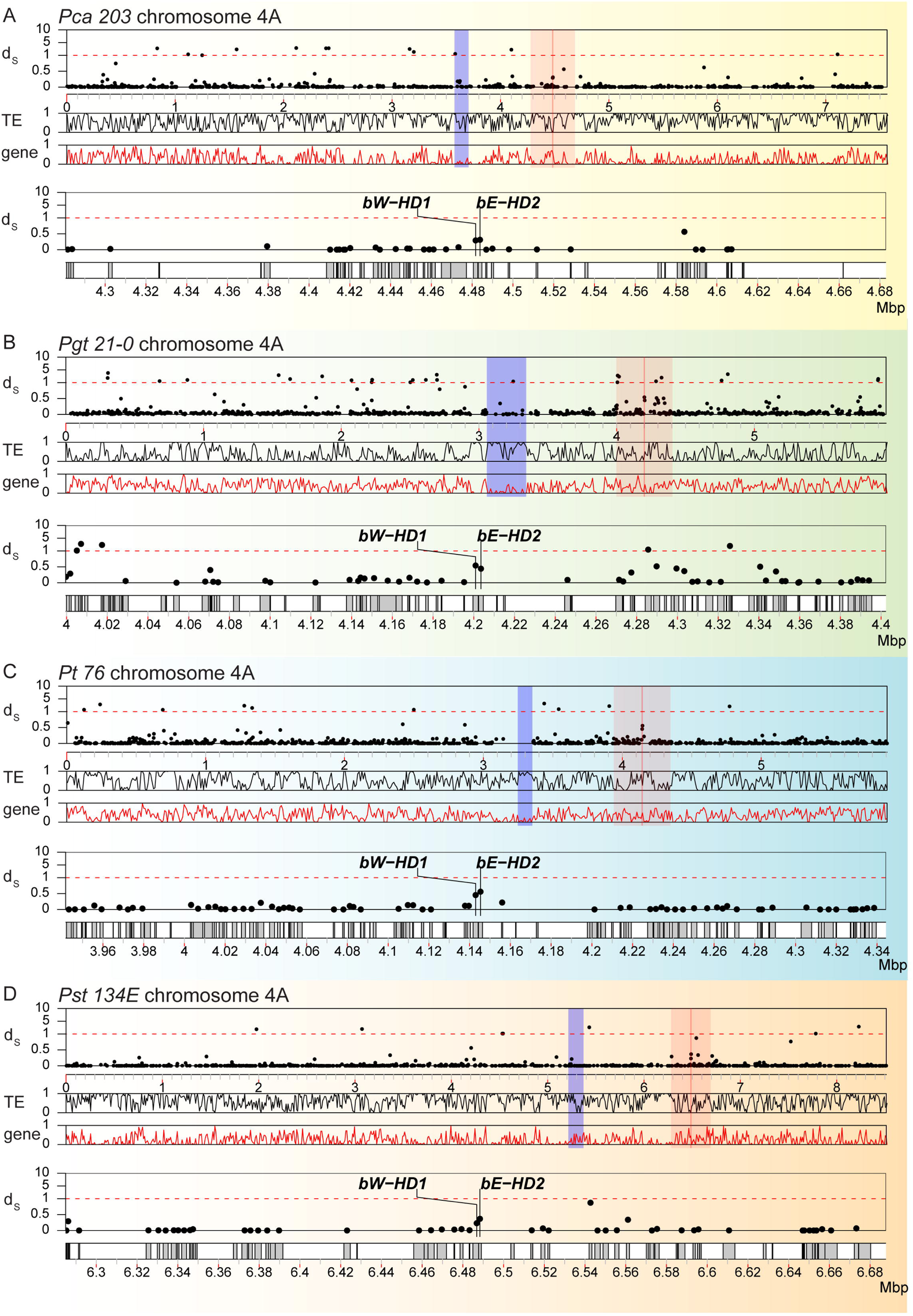
Synonymous divergence (d_S_) values of *HD* genes but not their immediate neighbors are slightly elevated on chromosome 4. Synonymous divergence values (d_S_) for all allele pairs are plotted along chromosome 4A for (A) *P. coronara* f. sp. *avenae* (“*Pca* 203”), (B) *P. graminis* f. sp. *tritici* (“*Pgt* 210”), (C) *P. triticina* (“*Pt* 76”), and (D) *P. striiformis* f. sp. *tritici* (“*Pst* 134E”). In each panel, the top track shows the d_S_ values (“d”) of allele pairs along chromosome 4. Each dot corresponds to the d value of a single allele pair. The second and third track shows the averaged TE (“TE”) and gene (“gene”) density along chromosome 4 in 10 kbp-sized windows, respectively. The *HD* genes (*bW-HD1* and *bE-HD2*) are highlighted with a red line and red shading indicates a 0.4 mbp-sized window around the *HD* locus. Predicted centromeric regions are marked with blue shading. The two lower tracks (d_S_ values and gene locations) provide a detailed zoomed in view of red shaded area around the *HD* locus. Species-specific background coloring is the same as for Fig 1.

Lastly, we investigated overall nucleotide and gene coding regions conservation for the *HD* locus including proximate regions containing 40 neighboring genes on either side of the locus. We combined fine scale nucleotide synteny with gene conservation analysis. We specifically tested if cereal rust fungi contain syntenic blocks with conserved protein coding genes surrounding *MAT* loci as it was found for species in the genus *Trichosporonales* (64). We used blastn to identify conserved nucleotide sequences within dikaryotic genomes and between species. We plotted genes, transposons, nucleotide and protein coding gene conservation for the *HD* loci proximate regions for all four cereal rust fungi (Fig 4A). Protein coding genes and their order are mostly conserved within dikaryotic genomes of the same species with 58/80 genes in *Pca 203*, 40/80 in *Pgt 21-0,* 57/80 in *Pt 76*, and 62/80 in *Pst 134E* being conserved. Similarly, we observed considerable nucleotide conservation and synteny within dikaryotic genomes with the exception of *Pgt 21-0*, which is consistent with our initial mummer-based analysis (S7 Fig). Overall, synteny and protein coding gene conservation was very limited between species for *HD* loci proximate regions. We could only identify three conserved genes across all four cereal rust fungi. These three genes code for an integral membrane protein (PTHR12459), D-2-hydroxyglutarate dehydrogenase (PTHR43716) and Glucose-6-phosphate 1-sepimerase (IPR025532) with all three clustering downstream of the *HD* locus (Fig 4A). Our TE analysis revealed that their coverage was overall consistent at the order level within species but varied across species (S12 Fig). For example, TIR and other undetermined class II DNA transposons dominated the *HD* locus of *Pst 134E,* while *HD* loci of other rust fungi displayed higher coverage of class I RNA transposons including LTRs.

**Fig 4.**
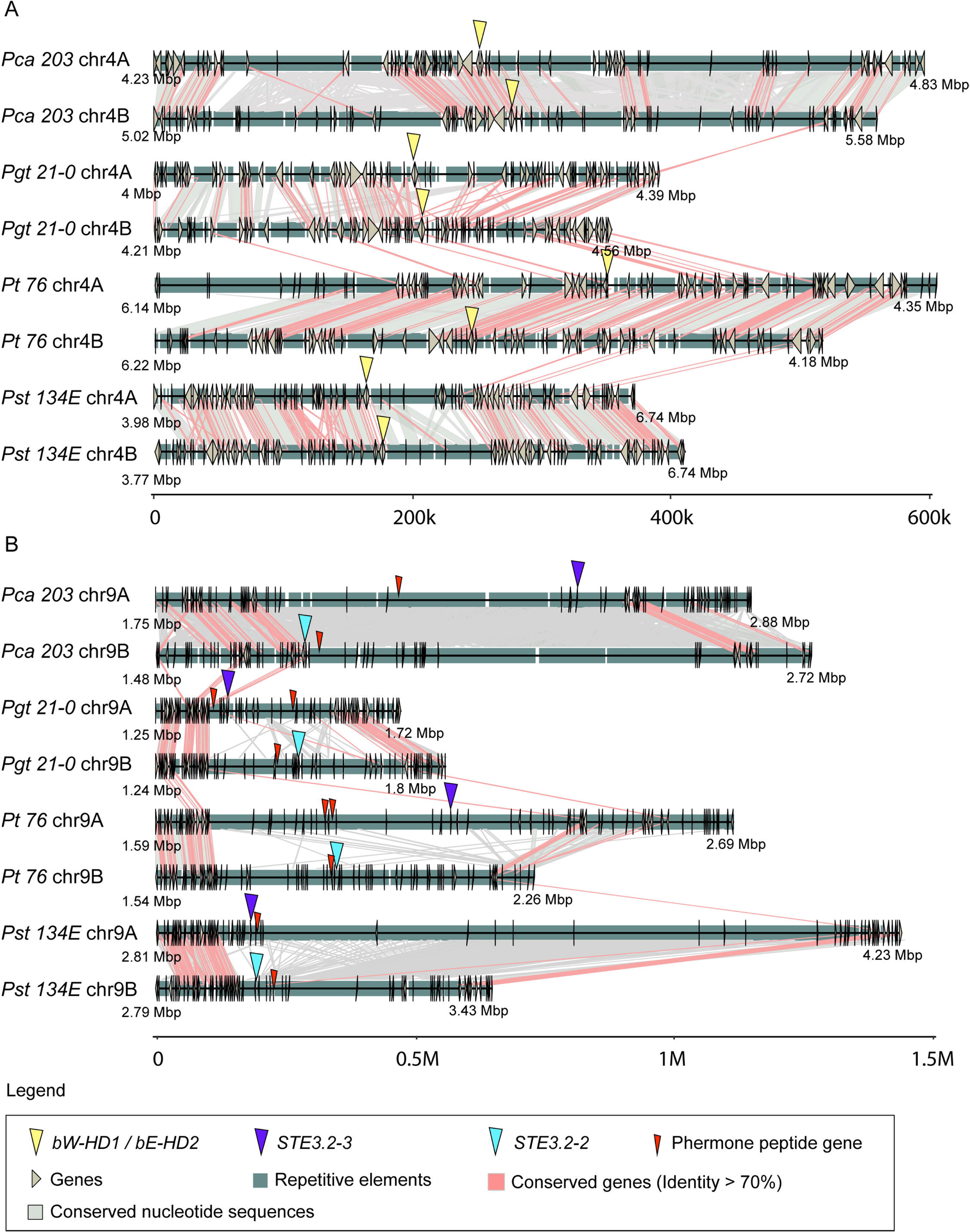
*HD* loci are partially conserved within species whereas *PR* loci are highly heterozygous and display strong signals of genomic degeneration. Synteny graphs of *HD* locus (A) and (B) *PR* locus including proximal regions in *P. coronara* f. sp. *avenae* (“*Pca* 203”)*, P. graminis* f. sp. *tritici* (“*Pgt* 210”), (C) *P. triticina* (“*Pt* 76”), and (D) *P. striiformis* f. sp. *tritici* (“*Pst* 134E”). Proximal regions are defined as 40 genes downstream and upstream of *HD* and *Pra* genes, respectively. *HD* loci are relatively syntenic within each dikaryotic genome but less conserved between species. *PR* loci display little synteny in *Pgt 21-0, Pt 76* and *Pst 134E* and show strong signs of transposon accumulation (see also S13 Fig). There is very little conservation of the *PR* loci across species. Red lines between chromosome sections represent gene pairs with identity higher than 70% and grey shades represent conserved nucleotide sequences (>=1000 bp and identity >=90%). For additional annotations please refer to the included legend (“Legend”).

Overall, our analysis of the *HD* locus suggests that it is overall conserved within each dikaryotic genome on a species level, while conservation has been eroded between species over longer evolutionary timescales.

### *PR* loci display strong signs of genomic deterioration

To investigate organization and synteny of the *PR* loci, we performed the same approach as for the *HD* loci. Initial mummer based synteny analysis revealed clear macro- and micro-synteny breaks around the *PR* loci in all cereal rust fungi (S13 Fig). It also revealed accumulation of highly repetitive sequences closely associated with *PR* loci, especially in *Pca 203* and *Pst 134E*. This increase in TEs correlates with a reduction in gene density around *PR* loci (Figs 4B and 5). The loss in synteny was specific to *PR* loci, as overall macro-synteny of chromosome 9 was conserved in all species (S13 Fig).

**Fig 5.**
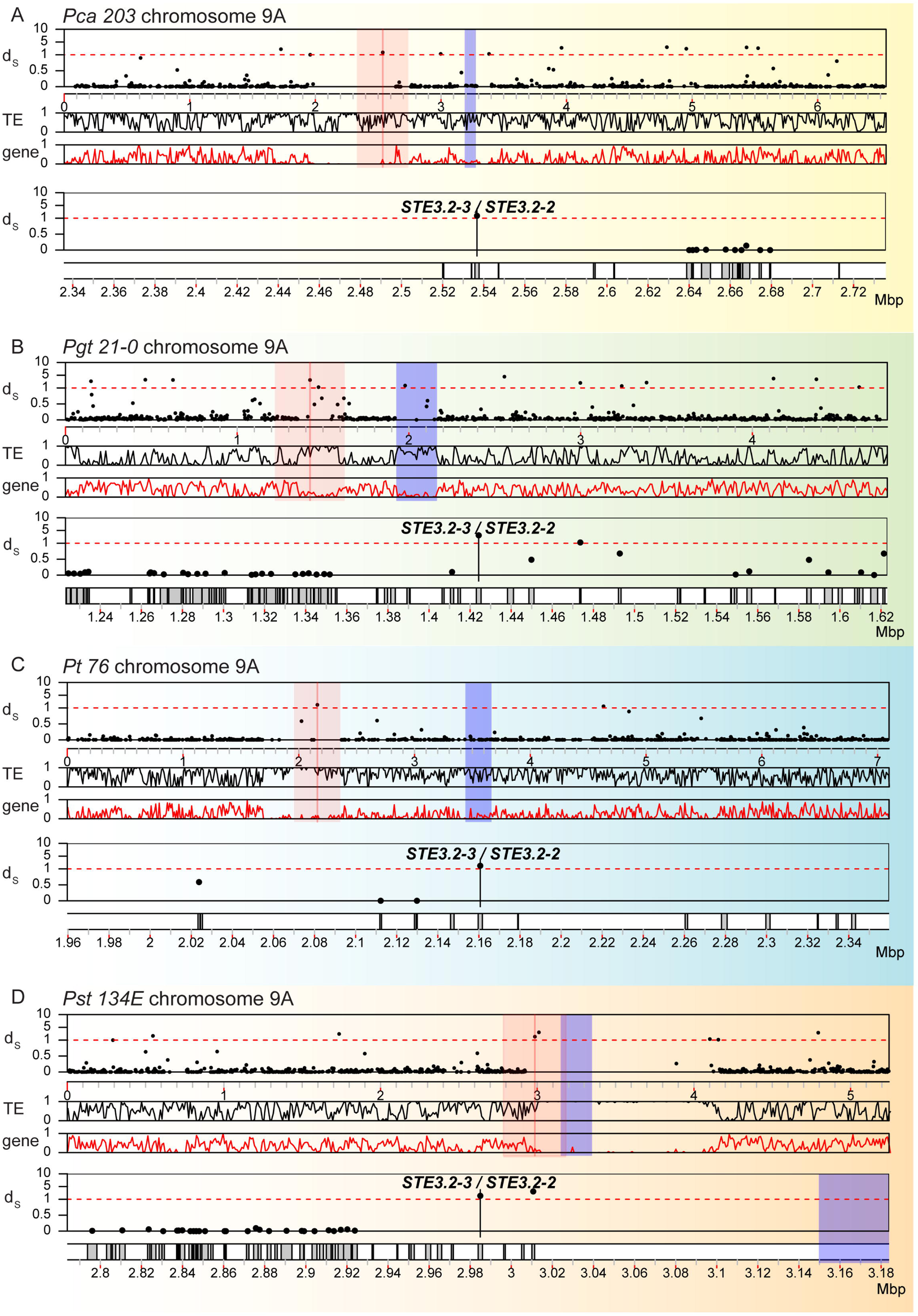
*PR* loci display elevated synonymous divergence (d_S_) values, accumulation of transposable elements and depletion of genes. Synonymous divergence values (d_S_) for all allele pairs are plotted along chromosome 9A for (A) *P. coronara* f. sp. *avenae* (“*Pca* 203”), (B) *P. graminis* f. sp. *tritici* (“*Pgt* 210”), (C) *P. triticina* (“*Pt* 76”), and (D) *P. striiformis* f. sp. *tritici* (“*Pst* 134E”). In each panel, the top track shows the d_S_ values (“d_S_”) of allele pairs along chromosome 9. Each dot corresponds to the d_S_ value of a single allele pair. The second and third track shows the averaged TE (“TE”) and gene (“gene”) density along chromosome 9 in 10 kbp-sized windows, respectively. The *Pra* alleles (*STE3.2-2* and *STE3.2-3*) are highlighted with a red line and red shading indicates a 0.4 mbp-sized window around the *PR* locus. Predicted centromeric regions are marked with blue shading. The two lower tracks (d_S_ values and gene locations) provide a detailed zoomed in view of red shaded area around the *PR* locus. Species-specific background coloring is the same as for Fig 1.

Consistent with the macro-synteny analysis, the d_S_ values for the *Pra* alleles *STE3.2-2*-*STE3.2-3* belonged to the 99.9% quantile of d_S_ values of all allele pairs found on chromosome 9 in all species (Fig 5 and S7 Fig). Gene density dropped significantly around *PRA* genes with *STE3.2-2*/*STE3.2-3* located within or adjacent to ‘gene deserts’ that extended over 1 mbps in the case of *Pst 134E*. Concomitant with gene depletion leading to gene deserts, *PR* loci are enriched in TEs. One exception was the *PR* locus of *Pgt 21-0*, which might be a biological feature or a technical artefact (see Discussion). None of the *Pra* alleles was closely linked to centromeres except for *Pst 134E STE3.2-3,* which was found in close proximity to the centromere at a distance of 164 kbp, embedded in a > 1 Mb long stretch of TEs depleted of genes (Fig 5).

Detailed nucleotide synteny and protein coding gene conservation analysis confirmed signals of extended recombination suppression, loss of synteny and accumulation of transposable elements around *PR* loci (Fig 4B). We observed little conservation of nucleotide sequences and protein coding genes in *Pra* proximate regions within dikaryotic genomes or between species. Indeed, we could not identify any conserved protein-coding gene in *Pra* proximate regions across the four cereal rust fungi. The TE composition in *Pra* proximate regions varied within dikaryotic genomes and between species (S12B Fig). This indicates the accumulation of distinct transposable element families around the *PR* locus in different cereal rust species (see more detail for *Pt* below).

Overall, the *PR* locus appears highly plastic with little conservation within and between species.

### *STE3.2-1* loci are highly conserved within and among cereal rust fungi

We analyzed *STE3.2-1* as above for the *MAT* loci. The three independent analyses revealed that *STE3.2-1* and its surrounding regions are highly conserved at the nucleotide and protein coding gene level within dikaryotic genomes while being overall syntenic between species (S14-16 Figs). This clearly sets apart the *STE3.2-1* locus from the two *MAT* loci which further supports the hypothesis that this locus is not involved in mate compatibility in rust fungi.

### *HD* genes are multiallelic in cereal rust fungi while *Pra genes* display little variation at the species level

We aimed to extend our species level analysis of variation in *MAT* genes (Figs 1 and 2) using publicly available genomic data for all four species (S1 Table). We identified between 12 and 15 representative isolates per species that were assigned to as many distinct genomic lineages as possible based on previous genealogical studies to avoid repeated sampling from clonal lineages that are predicted to show no genetic variation at *MAT* genes (30, 40-42, 44-46, 50-53) (S1 Table). We quality controlled whole genome short read datasets and mapped them against the respective reference genomes to estimate the genetic variation in *MAT* genes (S17-20 and 22-25 Figs). We identified high levels of polymorphisms at the *HD* loci for all analyzed cereal rust fungi. *Pca* showed the highest level of variation of *HD* loci at the species level (S18 Fig). When mapping Illumina short read data of the *Pca* isolates against the *Pca* 203 reference, the *HD* loci in the *Pca* isolates displayed high levels of polymorphism including heterozygous SNPs and multiple unaligned regions and gaps. This suggested that the *HD* loci present in our additionally selected *Pca* isolates were not well reflected in the *Pca 203* reference assembly. This is likely due to the fact that the selected isolates were highly diverse sexual populations of *Pca* sampled in the USA (40). We therefore implemented a novel *de novo* reconstruction approach for *HD* loci and *HD* coding sequences directly from whole genome Illumina short read datasets (see Methods for detail). We confirmed our approach by reconstructing the *Pca 203 HD* locus from publicly available Illumina short read datasets. We confirmed that the two reconstructed *HD* alleles were identical to the ones derived from the dikaryotic reference genome assembly (Fig 1) via a nucleotide alignment of the two *bW-HD1* and *bE-HD2* coding sequences to the reconstructed *HD* alleles (S21A Fig). In total, we were able to reconstruct eleven *bW-HD1* and twelve *bE-HD1* alleles for *Pca* (S21B Fig). Using this novel approach, we identified extensive genetic variation for *bW-HD1* and *bE-HD2* in *Pgt* and *Pt* (S21C-S21D Figs) while making use of recently published *bW-HD1* and *bE-HD2* alleles in case of *Pst* (21). We identified six, nine, and ten *bW-HD1* and six, eight, and ten *bE-HD1* alleles for *Pgt, Pt,* and *Pst*, respectively. Overall, this suggests that the *HD* genes are multiallelic in all four cereal rust fungi, which is consistent with other recent reports (21, 32).

Compared to the *HD* genes, the *Pra* genes are reported to be far less polymorphic in many basidiomycetes (9, 69, 70). Hence, we next focused our analysis on the *Pra* genes using the same whole genome Illumina short read datasets (S1 Table). As expected, the *Pra* genes were far less polymorphic within each species compared to *HD* loci (S22-26 Figs), which is consistent with our initial analysis of *Pra* genes extracted from genome assemblies (Figs 1 and 2). In contrast to the highly polymorphic *HD* genes, we were able to identify only two SNPs in the coding regions of *STE3.2-2* and *STE3.2-3* in *Pca* with a single SNP being non-synonymous (S26 Fig). Similarly, *STE3.2-2* and *STE3.2-3* copies of all *Pst* isolates had identical coding regions. Our short-read mapping analysis of *Pra* in *Pt* identified four SNPs in coding sequence regions of *STE3.2-2* and *STE3.2-3*. These SNPs gave rise to a single additional STE3.2-2 variant with two amino acid substitutions close to the C-terminus (S26 Fig). *Pgt* was the most polymorphic for *Pra*. We identified multiple distinct copies of *STE3.2-2* and *STE3.2-3* in the global *Pgt* population. We identified one STE3.2-2 variant with several amino acid changes (S26B Fig). *STE3.2-3* was the most polymorphic in *Pgt* including two isolates, *TTTSK* and *UVPgt60,* which contain potential nonsense mutations leading to pre-mature stop codons (S26B Fig). Overall, our analysis suggests that *Pra* is most likely biallelic in *Pca*, *Pt*, and *Pst* while *Pgt* may have more than two functional *Pra* alleles.

Lastly, we also investigated the nucleotide sequences of all *mfa* alleles at the species level. *Mfa1* and *Mfa2* alleles were fully conserved in *Pca* and *Pst*, while *Pt* had only a single non-synonymous change in *mfa1/3* (S3 Fig). In contrast, *mfa1* and *mfa3* in *Pgt* had several non-synonymous variations (S27 Fig). This species level variation in *Pgt* at the *mfa* loci is consistent with significant variation observed in *STE3.2-2* and *STE3.2-3*.

### *MAT* loci are conserved at the species level in *Puccinia triticina* while showing transposable element expansion specific to the *STE3.2-3* locus

We made use of four chromosome scale and fully phased genome assemblies for *Pt* (Table 1) to explore the structural conservation or plasticity of the *MAT* loci within a cereal rust fungal species. We performed macro-synteny, detailed nucleotide synteny, protein coding gene conservation and TE analysis for the *HD* and *PR* loci within and between *Pt 76* (43), *Pt 15* (55), *Pt 19NSW04* and *Pt 20QLD87* (32). A recent study suggested that *Pt 19NSW04* arose via somatic hybridization between *Pt 76* and *Pt 20QLD87* or close relatives in Australia mid-2010 (32). Hence, *Pt 19NSW04* and *Pt 76* share one complete nuclear haplotype B, which contains *STE3.2-2,* and *Pt 19NSW04* and *Pt 20QLD87* share one complete nuclear haplotype C, which contains *STE3.2-3*. The *Pt15* isolate is unrelated and was sampled in China in 2015.

Macro-synteny plots of the *HD* locus containing chromosomes suggested that the *HD* loci of shared nuclear haplotypes e.g. *Pt 19NSW04* and *Pt 76* hapB or *Pt 19NSW04* and *Pt 20QLD87* hapC, are more conserved when compared to *HD* loci contained within each dikaryotic genome (S6 and S28 Figs). Detailed nucleotide synteny and protein coding gene conservation analysis revealed that gene order and function is highly conserved in the *HD* locus between all tested nuclear haplotypes (S29 Fig). To investigate whether any specific TE families cluster around the *HD* locus, we built a TE database based on the *Pt 76* reference genome and used it to annotate TEs in all four strains. We identified similar TE coverage at the order level around the *HD* locus in all dikaryotic genomes (S30A Fig).

We performed identical analysis for the *PR* locus to get a species level understanding of its evolution in *Pt*. We compared the overall structural relationship between chromosomes containing *STE3.2-2* or *STE3.2-3* using macro-synteny plots. Like the *HD* locus, chromosomes of shared nuclear haplotypes, e.g. *Pt 19NSW04* vs *Pt 76* hapB with *STE3.2-2*, were more similar to each other than non-shared haplotypes, e.g. *Pt 76* hapB vs *Pt15* hapA with *STE3.2-3* (S13 and S31A Figs). The same applied for *STE3.2-2* (S13 and S31B Figs). We could clearly observe clustering of repetitive sequences around the *STE3.2-2* and *STE3.2-3* locus with the latter being more pronounced (S31 Fig). These repetitive sequences appeared to be allele specific as they were not visible in macro-syntenty plots comparing *STE3.2-2* and *STE3.2-3* containing chromosomes (S13 Fig). Detailed nucleotide synteny and protein coding gene analysis showed that genes around either *Pra* allele were mostly conserved while some haplotypes appeared to have inversions downstream of *STE3.2-2* relative to the *Pt 76* reference (Fig 6A and 6B). Gene synteny was more conserved for genes surrounding *STE3.2-3* yet gene distance appeared to be especially variable between the two *mfa1/3s* and *STE3.2-3* (Fig 6B). We explored the TE composition around each *Pra* allele and tested if TE expansions could be an explanation for the difference in intergenic distances around the *STE3.2-3* locus. TE composition around *STE3.2-2* and *STE3.2-3* were markedly different with the *STE3.2-3* locus displaying an increase of LTR retrotransposons (Fig 6C and S30B Fig.). TE coverage and composition around *STE3.2-2* was overall consistent between the different haplotypes (S30B Fig). In the case of *STE3.2-3* we found a single LTR *Ty3* (also known as Gypsy (71)) TE family (Ty3_Pt_*STE3.2-3*, Fig 6D) highly expanded at the locus with varying coverage between haplotypes ranging from 14.7% to 25.4% with *Pt 20QLD87* having the highest coverage (Fig 6C). The percentage identity relative to the consensus sequence was > 99% (Fig 6E) and most copies were around 8 kb (Fig 6F) which indicates active transposition of Ty3_Pt_*STE3.2-3* in recent history. To investigate whether this TE family was specific to the *STE3.2-3* locus, we compared its abundance at the locus with its abundance across the whole genome. We found that Ty3_Pt_*STE3.2-3* only clustered on *STE3.2-3* containing chromosomes in all four strains with only a very limited number of single copies found on other chromosomes (S32 Fig). Indeed, Ty3_Pt_*STE3.2-3* had a strong preference to insert between *Ptmfa1/*3s and *STE3.2-3* (S33 Fig). To further investigate the structure of Ty3_Pt_*STE3.2-3*, we used LTR_FINDER (72) to identify 5’ and 3’ LTR regions and other features related to retrotransposons. Ty3_Pt_*STE3.2-3* was found to have a short LTR, around 150 bp in all four strains, with target site repeat (TSR) region of ‘AAGT’, *pol* genes containing reverse transcriptase (RT), ribonuclease H (RH), integrase (INT). However, we failed to identify any group-specific antigen (*gag*) in these TEs (Fig 6D).

**Fig 6.**
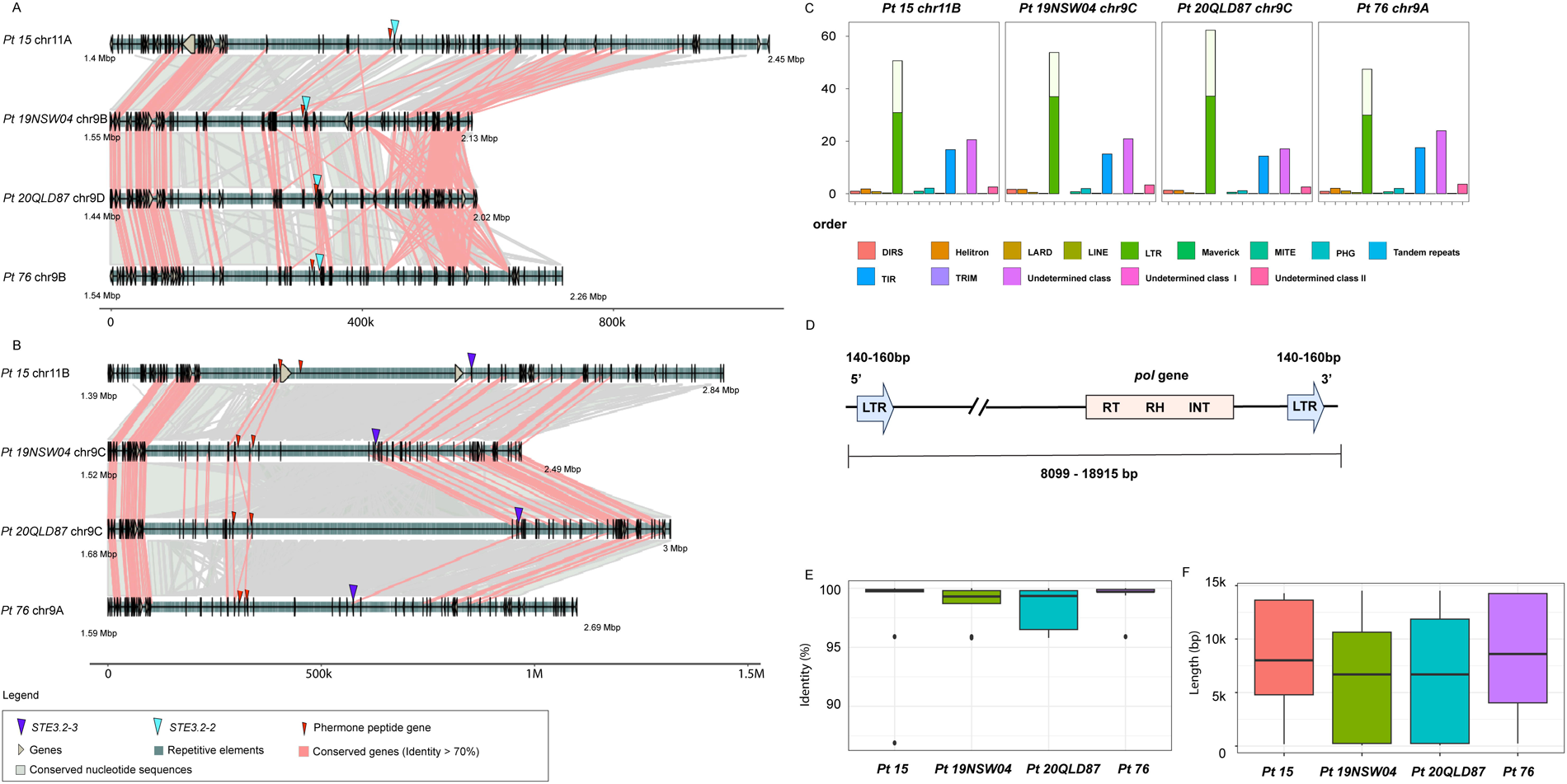
The *PR* locus shows an *STE3.2-3* allele specific accumulation of a unique Ty3-transposon family in *Puccinia triticina*. Synteny graphs of (A) *STE3.2-2* and (B) *STE3.2-3* including proximal regions in four different *P. triticina* isolates *Pt* 15, *Pt 19NSW04*, *Pt 20QLD87,* and *Pt 76*. Both loci are overall syntenic while the *STE3.2-3* locus displays an extension of the intergenic distance especially between the pheromone receptor and the two *mfa* genes. Red lines between chromosome sections represent gene pairs with identity higher than 70% and grey shades represent conserved nucleotide sequences (>=1000 bp and identity >=90%). For additional annotations please refer to the included legend (“Legend”). (C) Transposable element coverage at the order level at the *STE3.2-3* locus in the four different *P. triticina* isolates. (D) Structure of a representative Ty3_Pt_*STE3.2-3* copy. LTR – Long Terminal Repeat, RT – Reverse transcriptase, RH – RNAse H, and INT – Integrase. Distributions of (E) percentage identity relative to the consensus sequence and (F) lengths of individual Ty3_Pt_*STE3.2-3* insertions at the *STE3.2-3* locus in four different *P. triticina* isolates.

### *MAT* genes are expressed late in the asexual infection cycle during urediniospore production

Importance of *MAT* loci in sexual reproduction were well established for *Pt* and *Pst* (18, 73), yet it is unclear if they are expressed and functional during the asexual reproduction on cereal hosts as found in *Pt* (18). Hence, we investigated the expression patterns of *MAT* genes in *Pca*, *Pgt* and *Pst* using publicly available RNA-seq infection time series of their cereal hosts (38, 41, 48, 74) (S2 Table). We applied the trimmed mean of M-values (TMM) normalization to read counts and assessed quality of the RNA-seq datasets by multidimensional scaling (MDS) plots (S34 Fig). The MDS plots confirmed the suitability of the datasets for detailed expression analysis based on technical replicates clustering closely together and dimension 1 separating samples according to their infection progress. The stable expression of two house keeping genes for each species across all samples further confirmed the suitability of the datasets for detailed *MAT* gene expression analysis (S35 Fig.).

*STE3.2-2* and *STE3.2-3* were upregulated during the asexual infection process and always displayed highest expression at the latest time point available, which coincides with sporulation and production of urediniospores (Fig 7). We confirmed the differential expression at later infection timepoints with likelihood ratio test with a p-value cut-off of < 0.05 (S36 Fig). Similarly, *HD* genes are upregulated during asexual infection of the cereal host (Fig 7). We did not observe any expression of *STE3.2-1* in any of the samples. Lastly, we were also interested to see if we could detect RNA of *MAT* genes in cereal rust spores before infection and in specialized infection structures called haustoria. We used an additional *Pst* RNA-seq dataset and compared the expression of *MAT* genes in ungerminated spores, germinated spores, 5 days post infection, 9 days post infection and haustoria enriched samples (S37 Fig) (48). *HD* genes displayed some expression in spores albeit lower than observed in later stages of infection during spore production in the asexual cycle. In contrast, we could not detect expression of *Pra* genes in ungerminated and germinated spores. Yet, *Pra* genes were expressed and upregulated at later stages of the asexual infection consistent with other publicly available datasets (Fig 7C and S37 Fig).

**Fig 7.**
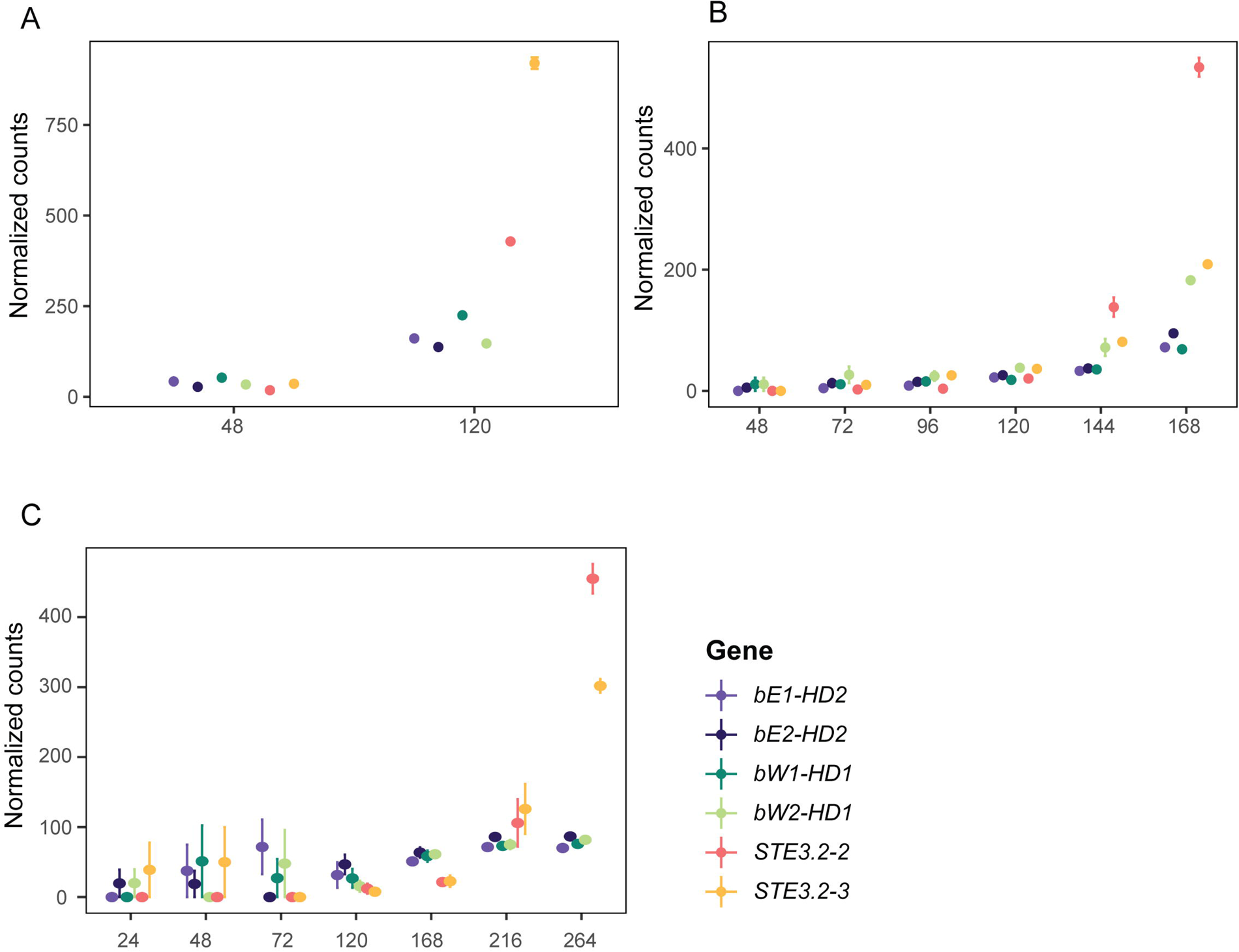
*MAT* gene expression is upregulated in the late stage of asexual infection cycle of the cereal host. (A) Trimmed mean of M-values (TMM)-normalized values of *MAT* genes in *P. coronara* f. sp. *avenae* (“*Pca 12NC29”)* at 48 and 120 hour post infection (hpi). (B) TMM-normalized values of *MAT* genes in *P. graminis* f. sp. *tritici* (“*Pgt 21-0*”) at 48, 72, 96, 120, 144 and 168 hpi. (C) TMM-normalized values of *MAT* genes in *P. striiformis* f. sp. *tritici* (“*Pst 87/66*”) at 24, 48, 72, 120, 168, 216 and 264 hpi.

## Discussion

Our comparative analysis of predicted *MAT* genes in four cereal rust fungi provides novel insight into the evolution of these genes and their proximal regions within and between species.

First, we confirmed that *HD* and *PR* loci are unlinked and located on two distinct chromosomes in all four cereal rust fungi, which confirms previous analysis on more fragmented genome assemblies or for individual rust fungal species (18, 21, 30). Further, both loci were heterozygous in all twelve dikaryotic genome assemblies. Taken together, these observations strongly support the hypothesis of tetrapolar mating types in rust fungi. Though this genome biology informed hypothesis requires experimental validation via crosses and genetic analysis of resulting offspring. As previously observed for *Pt* (18), the *Pra* homologs *STE3.2-1* were highly conserved in all cereal rust fungi and had no *mfa* genes closely associated. This suggests that *STE3.2-1* does not regulate mate compatibility.

In *P. coronata* f. sp. *avenae*, *P. striiformis* f. sp. *graminis* and *P. striiformis* f. sp. *tritici*, *STE3.2-1* is flanked by anaphase promoting complex *APC10/Doc1* which regulates important events in mitosis (75) and Elongation factor *EF1B* (76). Together, these findings suggest that *STE3.2-1* might play an important role during the cell cycle or different developmental programs as reported for other basidiomycete fungi (77, 78). However as previously reported for *P. triticina* (18), expression for *STE3.2-1* was absent or very low during the infection cycle of the cereal host.

Heterothallic fungi with tetrapolar mating types are most often multiallelic at least at one of the two *MAT* loci. Multiallelism is beneficial under outcrossing conditions, because it increases the compatibility between randomly selected mates. At the same time, outcrossing selects for new and rare alleles because these are compatible with existing and more common alleles (9). For example, *U. maydis* has many distinct transcription factor alleles at the *HD* locus (79, 80). We performed extensive analysis of *HD* loci extracted from dikaryotic genome assemblies and developed a workflow that permitted us to *de novo* reconstruct loci directly from Illumina short read sequencing data. This enabled us to investigate the allele status of *bW-HD1* and *bE-HD2* in 12-15 additional isolates for each cereal rust fungal species without the need for long-read genome assemblies or targeted amplicon sequencing. In addition, the analyzed isolates were selected based on them being the best available representation of global rust fungal lineages without repeated sampling within clonal lineages to facilitate a more in-depth analysis of the allele status of *MAT* loci. This extensive analysis demonstrated that the *HD* locus is also multiallelic in the four cereal rust fungal species. The four species had between six to twelve alleles of *bW-HD1* and *bE-HD2*. This supports previous analyses of the allele status of these genes in *P. striiformis* f. sp. *tritici,* which identified nine alleles of *bW-HD1* and *bE-HD2* (21). The number of *bW-HD1* and *bE-HD2* alleles is likely a gross underestimate giving limited public datasets. The majority of analyzed isolates were retrieved from the cereal host during the asexual reproduction cycle and from regions that are known to be deprived of sexually recombining populations. The only exception was *P. coronata* f. sp. *avenae* isolates, which are reported to be derived from sexual recombining populations in the United States of America (40). Indeed, we identified the highest allelic variation of *HD* genes (eleven *bW-HD1* and twelve *bE-HD2*) in *P. coronata* f. sp. *avenae*. This is similar to previous studies of isolates from likely sexual populations of *P. striiformis* f. sp. *tritici* in China, India, and Pakistan, which showed the highest diversity in *HD* alleles compared to other clonal lineages in Western wheat growing regions (21). In contrast to the multiallelic status of *HD* genes, the *Pra* and associated *mfa* genes were clearly biallelic in *P. coronata* f. sp. *avenae*, *P. triticina* and *P. striiformis* f. sp. *tritici*. In all cases, one *Pra* allele was linked to one specific invariant *mfa* allele whereby *P. triticina* carried two *mfa1/3* genes that were nearly identical in the sequence of the predicted mature pheromone peptide. This biallelism of *Pra/mfa* is similar to other tetrapolar dimorphic basidiomycetes like *U. maydis* and *Cryptococcus sp.* (81, 82). The notable exception was *P. graminis* f. sp. *tritici* which encodes two *mfa* genes (*mfa1* and *mfa3*) in proximity to *STE3.2-3*. The two encoded peptides are highly variable and are likely processed into two distinct mature pheromone peptides based on their sequence divergence, which might lead to distinct receptor activation specificities. Interestingly, *Pgt STE3.2-3* was also the most variable *Pra* allele in all species. We identified three *STE3.2-3* variants that have at least four or more non-synonymous nucleotide changes relative to the most common *Pgt STE3.2-3* variant. Moreover, the *STE3.2-3* variants in *Pgt 126-6711*, *Pgt ME-02*, and *Pgt PK-01* shared four distinctive amino acid changes in the C-terminus while being linked to an *mfa1* allele that gives rise to MFA1 with four amino acid substitutions. At the same time all analyzed *P. graminis* f. sp. *tritici* isolates carried an invariant *STE3.2-2* allele with only one *mfa* gene in close proximity and we did not identify a clearly distinct additional *Pra* allele. Hence the functional significance of the observed variation at the *STE3.2-3* locus is currently difficult to assess without more extensive sampling of isolates, more phased chromosome scale genome assemblies and in the absence of direct functional crossing studies in *P. graminis* f. sp. *tritici*. Yet, the organization of the *STE3.2-3* locus is reminiscent of the *PR* locus of *Sporisorium reilianum* in the order Ustilaginales. In *S. reilianum,* the *PR* locus is at least triallelic with each allele encoding one pheromone receptor and two distinct pheromone peptides. Each of the mature pheromones derived from one allele specifically activates only one of the receptors encoded by the two other alleles (83). Taken together, more studies are needed to assess if the *PR* locus in *P. graminis* f. sp. *tritici* is more than biallelic.

We initially predicted that biallelic *MAT* loci would show stronger signatures of recombination suppression and genomic degeneration than multiallelic loci based on published mathematical models (2) and observations in other basidiomycete species (9, 84). Indeed, the multiallelic *HD* loci were mostly syntenic within dikaryotic genomes of the four cereal rust species. This initial analysis with a limited set of genomes per species is likely to hold true at the population level. In the case of *P. triticina,* the eight *HD* alleles of four isolates were all highly syntenic with most of the genes being conserved between the different alleles. Consistently, the TE composition at the order level was highly similar between *HD* alleles of the same species. In contrast, *PR* loci showed strong signs of recombination suppression and genomic degeneration. However, the *PR* loci did not follow the exact same trends in all four species. The *PR* loci in *P. coronata* f. sp. *avenae*, *P. triticina* and *P. striiformis* f. sp. *tritici* were the most extended with the *Pra* genes being located at the edge or within large gene deserts and TE islands of ∼0.7 mbp, ∼0.7 mbp, and ∼1.2 mbp, respectively. At least some of these repeats were shared between the two *PR* alleles within each dikaryotic genome of *P. coronata* f. sp. *avenae* and *P. striiformis* f. sp. *tritici* based on whole chromosome alignments. This was not the case for *PR* alleles in *P. triticina*. In all three cases, there appeared to be differences in TE composition and coverage at the order level between the two *PR* alleles. This was most obvious for *P. triticina,* where we analyzed four dikaryotic genome assemblies and identified the *STE3.2-3* specific TE family Ty3_Pt_*STE3.2-3.* Ty3_Pt_*STE3.2-3* displayed preferential insertion for the *STE3.2-3* allele and insertions between the pheromone receptor and the two *mfa* genes. Ty3_Pt_*STE3.2-3* is likely a currently active TE based on its sequence conservation, length of individual TE copies, and the suggested nuclear exchange between the studied Australian isolates within the last decade (32). These initial observations of partial allele specific TE composition and coverage in *P. coronata* f. sp. *avenae*, *P. triticina* and *P. striiformis* f. sp. *tritici* contrast with observations of TE analysis of biallelic *MAT* loci in 15 *Microbotryum* spp.. The *MAT* loci in *Microbotryum* spp. did not display allele specific TE composition patterns, yet were likely reservoirs for TE families that drove TE expansion at a genome scale at discrete time points associated with the extension of evolutionary strata around the *MAT* loci (37).

Compared to these observations at biallelic *PR* loci, the *PR* locus in *P. graminis* f. sp. *tritici* contained fewer TEs, more genes, and was much shorter with ∼300 kbp. In addition, the TE composition and coverage was very similar between the two alleles within the dikaryotic genome of *Pgt 21-0*. These observations could be biological or a technical artifact of the assembly process of this specific genome. The *Pgt 21-0* assembly is based on older error prone PacBio long-read technologies, included extensive manual curation and relied on gene synteny between contigs and haplotypes for scaffolding. This gene based scaffolding might have been broken by long-stretches of TEs around the *PR* locus as observed for the three other cereal rusts and thereby introduced scaffolding errors in *Pgt 21-0* (30). Several of the other genomes generated with older error prone PacBio long-read technologies had issues assembling the *Pra* locus correctly. Alternatively, these marked differences of the *P. graminis* f. sp. *tritici PR* locus might reflect biological reality, as models predict that recombination suppression and genomic degeneration is more pronounced at biallelic loci (2). If the *PR* locus in *P. graminis* f. sp. *tritici* is not truly biallelic as potentially indicated by the gene-based analysis of *Pra* and *mfa* alleles, the organization and level of genomic degeneration at the *PR* locus as observed would fulfill these predictions. However, more and better high-quality phased chromosome scale genome assemblies are necessary to differentiate these alternative hypotheses concerning the *PR* locus in *P. graminis* f. sp. *tritici*.

The functions of *MAT* genes in rust fungi are currently unknown, beyond their likely role in mediating mate compatibility. However, in *P. triticina* and *M. larici-populina* (35, 85), *MAT* genes are expressed during the sexual and asexual infection cycle. Suppression of *MAT* gene expression in *P. triticina* during the asexual infection cycle on wheat reduces infection severity and spore production (18). Here, we show that *MAT* genes are also upregulated during the late stage of the asexual infection cycle of the cereal host in *P. coronata* f. sp. *avenae*, *P. graminis* f. sp. *tritici,* and *P. striiformis* f. sp. *tritici*. This suggests a broader conservation of *MAT* gene expression during the late stages of the asexual infection cycle of rust fungi. Rust fungal cells carry more than two nuclei in the same cytosol during the asexual infection cycle (86). At the same time, ureodiniospores always carry two nuclei of compatible mating types and never two nuclei of identical mating types. Studies from the early 20^th^ century show that the two nuclei of ureodiniospore mother cells undergo synchronous nuclear division within the same cytoplasm giving rise to four nuclei within the same ureodiniospore precursor cells. This is followed by nuclear movement to restore appropriate nuclear pairing and planar cell division that enables ureodiniospore maturation and reconstitutes the dikaryotic state (86). The observed upregulation of *MAT* genes aligns with the development of ureodiniospore primordia development and spore production. Hence, we hypothesize that *MAT* genes are involved in the appropriate nuclear pairing at this stage of fungal development.

Understanding mating types in cereal rust fungi has also significant implications for agriculture and for the prediction of the evolution of new virulent isolates. Recent reports suggest that nuclear exchange between isolates during the asexual infection cycle (also known as somatic hybridization (87)), in addition to sexual reproduction, can lead to nuclear assortment (87). Nuclear exchange and the viability of the resulting offspring is likely regulated by *MAT* genes because all proposed events have given rise to isolates with dikaryotic genomes that carry opposing *MAT* gene pairs. We confirmed these observations with our *MAT* gene analysis in the case of *Pgt Ug99* (*Pgt 21-0* and unknown nuclear donor) and *Pt 19NSW04* (*Pt 76* and *Pt 20QLD87* as nuclear donors). In reverse, our analysis showed that nuclear exchange has not occurred for the analyzed isolates of *P. coronata* f. sp. *avenae*, and *P. striiformis* f. sp. *tritici*, because none of the isolates share identical *HD* and *Pra* alleles that could be carried in the same nucleus. In cases of nuclear exchange, these nuclei behave like entire linkage groups with virulence (also known as effectors) allele complements being tied to specific *MAT* alleles if somatic hybridization occurs in the absence of parasexual reproduction as currently proposed. This implies that the nuclear *MAT* alleles define the possible virulence allele combinations that can arise via nuclear exchange. We hence can predict the most likely novel virulence allele combinations that might arise given circulating nuclear genomes in existing cereal rust fungal populations on their cereal hosts. This observation requires us to generate phased and chromosome scale genome assemblies of all cereal rust fungi at the population level. These will improve disease management strategies and the durability of characterized resistance loci in the global wheat germplasm leading to smarter rust resistance breeding programs.

## Acknowledgments

We thank Dr. P. Tobias and Dr. M. Moeller for critical comments on manuscript, Dr. M. Bui for advice in iqtree2 usage, Prof. C. Linde for useful suggestions, and R. Tam for providing a snakemake pipeline for trimming raw sequencing data and centromere information of *Pst 134E*. We thank J. Lin for suggestions on data visualization. We acknowledge the excellent and critical suggestions of the editor and peer reviewers that significantly improved the manuscript.

## Material and methods

### Assessment of used genomic resources

Detailed information on isolates and genomes used in this study can be found in Table 1. Completeness of genome assemblies were assessed by BUSCO v5.5.0 (88) in the genome mode with a basidiomycota_odb10 set of genes.

### Identification of *MAT* genes

*MAT* genes including the *HD* and *PRA* genes previously identified in *P. triticina* (Table 2) (18) were used as blastp and blastn (89) query to identify orthologs in *P. graminis* f.sp *tritici Ug99* and *21-0* isolates (30), *P. striiformis* f.sp *tritici 134E* (47), *104E* (48) and *DK0911* (49), *P. triticina 76* (43), *19NSW04*, *20QLD87* (32) and *15* (55), *P. polysora GD1913* (63), *P. coronate* f.sp *avenae 203* (39), *12NC29* and *12SD80* (38) isolates. Missed *STE3.2-3* in *Pst 104E* was fixed by mapping short reads to reference genome *Pst 134E* with bwa-mem2 (90). *Mfa genes* were identified by using *mfa* of *Pt BBBD*, *Pgt SCCL* and *Pst 78* (18) as a custom motif database to search similar pattern in all studied species with Geneious Prime v. 2022.1.1 (91), using a function built from fuzznuc (92). Karyograms of *Pca 203*, *Pgt 21-0*, *Pt 76* and *Pst 134E* were made with RIdeogram 0.2.2 (93).

### Genealogical analysis of the *HD* and *Pra* genes

To construct a species tree for cereal rust fungi, protein sequences encoded in one nuclear genome from each cereal rust fungus species under study was used as input data. The input data was processed by Orthofinder 2.5.5 (61, 94) to infer maximum likelihood trees from multiple sequence alignments (MSA), fasttree v2.1 (95) was used to calculate likelihood-based local support values. Coding sequences of identified *HD* and *Pra* genes were aligned with MACSE v2.07 (96) with default setting. Coding-sequence based alignments were trimmed by using TrimAl v1.2 (97) with option (-gc 0.9) to remove sequences which have gaps in more than 10% of samples, trimmed alignments were realigned with MACSE again. Nucleotide alignments were imported into BEAUti v2.7.6 (98) to generate .xml files, bModelTest 1.3.3 (99) was used to test the best nucleotide substitution model separately with a Markov chain length by 15 million. HKY+G was chosen for inferring two *HD* genes, whereas TN93+I was chosen for *Pra* genes and *STE3.2-1* individually. Then BEAST 2 MCMC runs under a strict molecular clock with the Yule Model tree, a 15 million length Markov chains was used to provide a sufficient effective sample size (ESS) of independent samples. Tracer v.1.7.2 (100) was used to check if ESS values of all parameters are above 200 for each run. TreeAnnotator (98) was used to generate a maximum clade credibility tree with 10% burn in. Figtree v.1.4.4 (101) was used to visualize the final output.

Nucleotide sequences of identified *mfa* genes were aligned with MACSE v2.07 (96) with default settings, amino acid alignment generated was used to construct maximum likelihood tree via IQ-TREE 2 (version 2.1.4-beta) (102), the best model was estimated by ModelFinder (-m MFP) (103) based on Akaike Information Corrected Criterion (AICc) score (104), 10000 replicates were set for ultrafast bootstrap (105). JTT+G4 substitution model was chosen by ModelFinder (103).

To compare topologies of *bW-HD1* and *bE-HD2* gene trees, IQ-TREE 2 (version 2.1.4-beta) (102) was used to build unrooted maximum likelihood trees for each species and both genes separately, the best model for each alignment estimated by ModelFinder (-m MFP) (103) based on AICc score (104), 10000 replicates were set for ultrafast bootstrap (105). Top 100 optimal gene trees of *bW-HD1* and *bE-HD2* were used as alternate topologies after removing branch length values to perform AU tests (62) with IQ-TREE2 (-keep-ident –zb 10000 −n 0 - au), p_AU_ <0.05 suggesting topological incongruence between the two *HD* genes.

### Estimation of centromeric regions

Hi-C reads of *Pca 203* and *Pt 76* were mapped back to their corresponding haploid genome assembly with Juicer 1.6 (68) and processed with the 3d-dna (106) pipeline. Juicerbox 1.8.8 (68) was used to visualize Hi-C maps. The approximate positions of centromeric regions were estimated by zooming in on the Hi-C heatmap in Juicerbox and selecting the region corresponding to strong interaction signals, which has been validated and widely used in previous studies (107, 108).

### d_s_ value estimation for allele pairs and plotting

Inspired by the approach used by Branco et al. (8) for identifying evolutionary strata on mating-type chromosomes of *Microbotryum* species, proteinortho 6.0.22 (109) was used to pair orthologs one-to-one with-synteny and −singles tag. Muscle 3.8.1551 (110) was applied to generate coding sequence based protein alignments with default settings. PAML 4.9 (111) was used to calculate synonymous divergence values from alignments generated with Muscle. Genomes were split in 10 kbp per window, then density of genes and repeats were calculated with bedtools v 2.29.2 (112). Results were visualized with karyoploteR v3.17 (113).

### Synteny analysis of *MAT* loci and flanking regions

Self-alignment analyses between two haplotypes of each species were done with Nucmer (version 3.23) (65) with –maxmatch option, alignments with a size of less than 1000 bp or less than 90 % identity were removed, filtered data were visualized via Matplotlib (114).

Nucmer (version 3.23) (65) was used to align nucleotide sequences of *MAT* regions.

Blastn 2.10.0 (89) was used to identified conserved genes (identity higher than 70%, e-value < 0.05) across and within species. Results were visualized with gggenomes 0.9.9.9000 (115).

### General repeat annotations

REPET v3.0 pipeline (116, 117) was applied to annotate repeats in all genomes used in this study, together with Repbase v22.05 (118), config files were based on default settings. TEdenovo was used to predict novel repetitive elements and to construct specific custom databases from each genome individually as described in the manual instructions. Constructed custom databases were used for repeat annotation with TEannot. The reference TE database built for *Pt 76* from TEdenovo with the REPET v3.0 pipeline was used as reference for annotating repeats in *Pt 15* (55), *Pt 19NSW04* and *Pt 20QLD87* (32) with RepeatMasker 4.1.5 (119) To make analyses consistent, the *Pt 76* genome was reannotated in the same way as the other three isolates.

To calculate the coverage of each TE family, classification information from TEannot was used. For TEs with more than one potential classification, hits from Repbase which have more than 70% of similarity were used as evidence for reclassification.

LTR_FINDER v1.07 (72) with ps_scan (120, 121) was used to identify detailed structures of LTRs of Ty3_Pt_*STE3.2-3* in default setting. Reverse transcriptase (A0A0C4EV49_PUCT1), Retrotransposon gag domain (A0A0C4EKM8_PUCT1), Integrase, catalytic domain (A0A0C4EMG6_PUCT1) and Retropepsins (A0A0C4F2V2_PUCT1) (122), were additionally used as query to search for motifs in identified LTR retrotransposons using tblastn 2.12.0+ (89).

### Identification of variants in *MAT* loci

Sequence data of *Pca* were published by (40), data of *Pgt* were published by (30, 41, 42, 53) and under PRJNA39437, data of *Pt* used in this study have been published earlier by (44-46) and under PRJNA39803 and PRJNA39801, data of *Pst* were published by (50-52) and under PRJNA60743, all above data were downloaded from NCBI SRA database (URL: https://www.ncbi.nlm.nih.gov/sra) (Table S1). Four *Pca* isolates, *Pca 203* (39), *Pgt 21-0* (30), *Pt 76* (43) and *Pst 134E* (47), with published chromosome-scale haplotype-phased genome assemblies were used as reference for mapping. Trimmomatic 0.39.2 (123) was used to remove adapters from raw reads whereas FastQC v0.11.8 (124) was applied for assessing the quality before and after trimming. Bwa-mem2 (90) was used to map trimmed reads to reference genomes, MarkDuplicates (Picard) 1.70 (125) was applied to remove PCR-generated duplicates. Samtools 1.12 (126) was used to remove mapped reads with mapping quality lower than 30. IGV 2.16.2 (127) and Qualimap 2 (128) were both used to check mapping quality.

For reconstruction of *Pra* alleles, freebayes-parallel 1.3.6 (129, 130) was applied to detect variants, bcftools 1.12 (126) was used to exclude low quality variants with ‘QUAL<40’. Samtools 1.12 (126) was used to filter out regions of interest. The Ensembl Variant Effect Predictor (VEP) 88.9 (131) was used to annotate sequence variants. Bcftools 1.12 (126) was used to generate consensus sequences, nucleotide and protein alignments were generated and visualized with Geneious Prime v. 2022.1.1 (91).

For *de novo* reconstruction of *HD* alleles, Spades (132), a de novo assembler, was used for building draft genome assemblies from available whole genome Illumina short read datasets (Table S1). For isolates with average read lengths > 101 bp, k-mer size of 101 was chosen for building draft genomes. For isolates with average read lengths < 101 bp, k-mer size of 51 bp was chosen. *HD* genes used in genealogical analysis were used as reference genes, blastn 2.12.0+ (89) was used to search and identify *HD* allele containing contigs of the draft genome assemblies. KAT (133) was used to compute a k-mer matrix from *de novo* assembled *HD* contigs and reference *HD* loci sequences with default setting (k-mer size=27). KAT filter was used to obtain all paired-end reads which contained *HD* k-mers based on the created *HD* k-mer matrix (-T 0.2). Spades was used to reassemble all *HD* related paired-end reads of the specific isolate. Final outputs were visualized in Geneious, CDS regions of reconstructed *HD* alleles were predicted by mapping the CDS of reference *HD* alleles to reconstructed contigs, ORFs were predicted with Geneious.

### Comparison of *MAT* loci among *Pt* isolates

Fully phased genome assemblies of *Pt 15* (55), *Pt 19NSW04* and *Pt 20QLD87* (32) were downloaded from NCBI SRA database (URL: https://www.ncbi.nlm.nih.gov/sra), annotation of *Pt 19NSW04* and *Pt 20QLD87* were downloaded from CSIRO Data Access Portal (URL: https://data.csiro.au/). CDS and protein data of *Pt 19NSW04* and *Pt 20QLD87* were generated with Gffread (134), *MAT* genes were identified with blastn 2.12.0 (89) using *MAT* genes in *Pt 76* as query. Nucmer 3.2.3 was applied to make pair-wise comparisons between *Pt* isolates, -- maxmatch tag was used to allow identification of maximum hits, macrosynteny plots were generated with Matplotlib (114) after removing matches with < 90% identity and shorter than 1000 bp. Blastn 2.10.0 (89) was used to identify orthologs with at least 70 % identity. Microsynteny plots were generated with gggenome2 (115).

### Comparison of expression of *MAT* genes in asexual reproduction

RNA-seq reads of rust fungi on infected plants were obtained from NCBI SRA database (URL: https://www.ncbi.nlm.nih.gov/sra). The datasets included: RNA-seq data of *Pst* 87/66 on infected wheat (PRJEB12497 (74), *Pca* 1*2NC29* on *Brachypodium distachyon* (PRJNA398546 (38), *Pgt* 21-0 on infected wheat (PRJNA415866 (41), *Pst 104E* (PRJNA396589 (48). Raw reads were quality checked with FastQC v0.11.8 (124), trimmomatic 0.50 (123) was used to trim low quality reads with parameters: ILLUMINACLIP:adapter.fa:2:30:10 LEADING:3 TRAILING:3 SLIDINGWINDOW:4:15 MINLEN:36. Kallisto 0.44.0 (135) was then applied to align and count transcripts with −b 100 parameter for all paired-end reads and −b 50 −l 200 −s 20 --single --single-overhang for single-end reads. *Pst 104E* cDNA data was used as reference for *Pst 104E* infected tissues. Since no reference genome of *Pst 87/66* is available, cDNA data of *Pst 134E* was used as reference instead with *HD* genes of *Pst 87/66* supplemented to obtain better quantification of the specific *HD* alleles. *Pgt 21-0* and *Pca 12NC29* were used as reference for mapping reads from *Pgt 21-0* and *Pca 12NC29* infected tissues, respectively. Gene names of *Pca 12NC29* and *Pst 104E* were added with funannotate annotate v 1.8.5 (136, 137) with default parameters. Kallisto outputs were imported to EdgeR 3.17 (138) with tximportdata (139) following user instruction, followed by normalizing with the TMM method, significant upregulation of *MAT* genes was assessed with the LRT method and ggplot2 (140) was used to visualize the result. House keeping genes were identified in each dataset based on the following conditions: *p-value*>0.1 and log(FC) <0.5, functional annotations of candidate house keeping genes were retrieved from interproscan (137).

### Data Availability

Analysis code used in this study is available at github repository: https://github.com/ZhenyanLuo/codes-used-for-mating-type. Alignments of genealogical studies, d_s_ value of gene pairs in studied cereal rust species, CDS of reconstructed *HD* alleles, *Pra* alleles, TE annotation and classification files are available at Dryad (https://doi.org/10.5061/dryad.w0vt4b8zm).

## Notes

https://github.com/ZhenyanLuo/codes-used-for-mating-type

https://datadryad.org/stash/share/XCyaQ2j8q8yI28amg3ct1WplAOlh-yTUHfaz_jyDYK4

### Competing Interest Statement

The authors have declared no competing interest.

### Summary of Updates

The content is the same but the "Full Text" version 2 on Biorxiv had the panels for Fig 6 and Fig 7 swapped. Not sure why. The Preview pdf had the right order.

