## Supplemental Figures for "Genome biology and evolution of mating type loci in four cereal rust fungi"

**Table S1 List of whole genome Sequence Read Archive data used in the present study.**

The table provides information on all whole genome Sequence Read Archive (SRA) data used in the present study. Metadata includes species abbreviation ("Species"), isolate name ("Isolate"), SRA identifier ("SRA ID"), and the initial reference ("Citation"). *Pca* - *P. coronata* f. sp. *avenae*, *Pgt* - *Puccinia graminis* f. sp. *tritici*, *Pt* - *P. triticina* and *Pst* - *P. striiformis* f. sp. *tritici.*

| Species | Isolate | SRA ID | Citation |
|---|---|---|---|
| *Pca* | *15Fl1-4* | SRR7062959 | (1) |
| | *15MN16-3* | SRR7062966 | |
| | *15MN18-1* | SRR7062962 | |
| | *15MN27-3* | SRR7062951 | |
| | *15NE9-3* | SRR7062952 | |
| | *15OH12-3* | SRR7062922 | |
| | *15SD30-1* | SRR7062937 | |
| | *90GA16-1* | SRR7062947 | |
| | *90MN1B-1* | SRR7062928 | |
| | *90MN5B-1* | SRR7062929 | |
| | *90MN7B-1* | SRR7062924 | |
| | *90MN149-2* | SRR7062917 | |
| | *90TX45-1* | SRR7062945 | |
| | *90TX52-1* | SRR7062943 | |
| | *90TX58-1* | SRR7062940 | |
| *Pgt* | *126-6711* | SRR1593673 | (2) |
| | *326-12356* | SRR1593681 | |
| | *AF4NF* | SRR5883047 | (3) |
| | *IR-01* | ERR2099124 | (4) |
| | *IS-03* | ERR2099132 | |
| | *ME-02* | ERR2099134 | |
| | *PK-01* | ERR2099136 | |
| | *SA-01* | ERR2099140 | |
| | *TTTSK* | SRR039024 | PRJNA39437 |
| | *UVPgt55* | SRR9024811 | (5) |
| | *UVPgt59* | SRR039039 | |
| | *UVPgt60* | SRR9024809 | |
| | *UVPgt61* | SRR9024810 | |
| *Pt* | *9* | SRR035305 | PRJNA39803 |
| | *161* | SRR035303 | PRJNA39801 |
| | *60-L-2* | SRR4254457 | (6) |
| | *630846* | SRR4254442 | |
| | *890155* | SRR4254451 | |
| | *900084* | SRR4254449 | |
| | *ETH-4114-7* | SRR11070891 | (7) |
| | *FRA-5.3* | SRR11070888 | |
| | *ISR-850* | SRR11070886 | |
| | *MCTSB_11US116_1* | SRR11479798 | |
| | *S423-wlr* | SRR11177218 | (8) |

| | | | |
|---|---|---|---|
| | *S459-wlr* | SRR11177217 | |
| | *S472-wlr* | SRR11177215 | |
| | *TBBGJ_11US136_1* | SRR11071601 | (7) |
| *Pst* | *87/27-s2* | SRR1534348 | (9) |
| | *87/66-s1* | SRR1534347 | |
| | *98/83-s3* | SRR1534208 | |
| | *110B* | SRR15496042 | (10) |
| | *s414* | SRR13781820 | |
| | *s572* | SRR13799080 | |
| | *s674_WYR* | SRR13823727 | |
| | *SA1* | ERR3856826 | (11) |
| | *SA2* | ERR3856827 | |
| | *SA3* | ERR3856828 | |
| | *SA4* | ERR3856829 | |
| | *W010* | SRR15496040 | (10) |

**Table S2 List of RNAseq Sequence Read Archive data used in the present study.**

The table provides information on all RNAseq Sequence Read Archive (SRA) data used in the present study. Metadata includes species abbreviation ("Species"), isolate name ("Isolate"), timepoint of infection (hpi, hours post infection) or sample type ("Sample Type"), timepoint/status of spores ("Time point/Status"), replicate number ("Replicate"), SRA identifier ("SRA ID"), and the initial reference ("Citation"). *Pca* - *P. coronata* f. sp. *avenae*, *Pgt* - *Puccinia graminis* f. sp. *tritici*, and *Pst* - *P. striiformis* f. sp. *tritici.*

| Species | Isolate | Sample Type (hpi) | Replicates | SRA ID | Citation |
|---|---|---|---|---|---|
| *Pca* | *12NC29* | 48 | 1 | SRR5976045 | (12) |
| | | | 2 | SRR5976121 | |
| | | | 3 | SRR5976122 | |
| | | 120 | 1 | SRR5976046 | |
| | | | 2 | SRR5976047 | |
| | | | 3 | SRR5976048 | |
| *Pgt* | *21-0* | 48 | 1 | SRR6242049 | (3) |
| | | | 2 | SRR6242050 | |
| | | | 3 | SRR6242047 | |
| | | 72 | 1 | SRR6242051 | |
| | | | 2 | SRR6242052 | |
| | | | 3 | SRR6242054 | |
| | | 96 | 1 | SRR6242055 | |
| | | | 2 | SRR6242056 | |
| | | | 3 | SRR6242053 | |
| | | 120 | 1 | SRR6242057 | |
| | | | 2 | SRR6242058 | |
| | | | 3 | SRR6242061 | |
| | | 144 | 1 | SRR6242062 | |
| | | | 2 | SRR6242059 | |

| | | | | | |
|---|---|---|---|---|---|
| | | | 3 | SRR6242060 | |
| | | 168 | 1 | SRR6242063 | |
| | | | 2 | SRR6242035 | |
| | | | 3 | SRR6242037 | |
| *Pst* | *87/66* | 24 | 1 | ERR1224556 | (13) |
| | | | 2 | ERR1224557 | |
| | | | 3 | ERR1224558 | |
| | | 48 | 1 | ERR1224559 | |
| | | | 2 | ERR1224560 | |
| | | | 3 | ERR1224561 | |
| | | 72 | 1 | ERR1224562 | |
| | | | 2 | ERR1224563 | |
| | | | 3 | ERR1224564 | |
| | | 120 | 1 | ERR1224565 | |
| | | | 2 | ERR1224566 | |
| | | | 3 | ERR1224567 | |
| | | 168 | 1 | ERR1224568 | |
| | | | 2 | ERR1224569 | |
| | | | 3 | ERR1224570 | |
| | | 216 | 1 | ERR1224571 | |
| | | | 2 | ERR1224572 | |
| | | | 3 | ERR1224573 | |
| | | 264 | 1 | ERR1224574 | |
| | | | 2 | ERR1224575 | |
| | | | 3 | ERR1224576 | |
| | *104E* | Ungerminated spore | 1 | SRR6043978 | (14) |
| | | | 2 | SRR6043979 | |
| | | | 3 | SRR6043971 | |
| | | Germinated spores | 1 | SRR6043973 | |
| | | | 2 | SRR6043972 | |
| | | | 3 | SRR6043975 | |
| | | 144 | 1 | SRR6043970 | |
| | | | 2 | SRR6043968 | |
| | | | 3 | SRR6043967 | |
| | | 216 | 1 | SRR6043966 | |
| | | | 2 | SRR6043965 | |
| | | | 3 | SRR6043969 | |
| | | Haustoria enriched samples | 1 | SRR6043974 | |
| | | | 2 | SRR6043977 | |
| | | | 3 | SRR6043976 | |

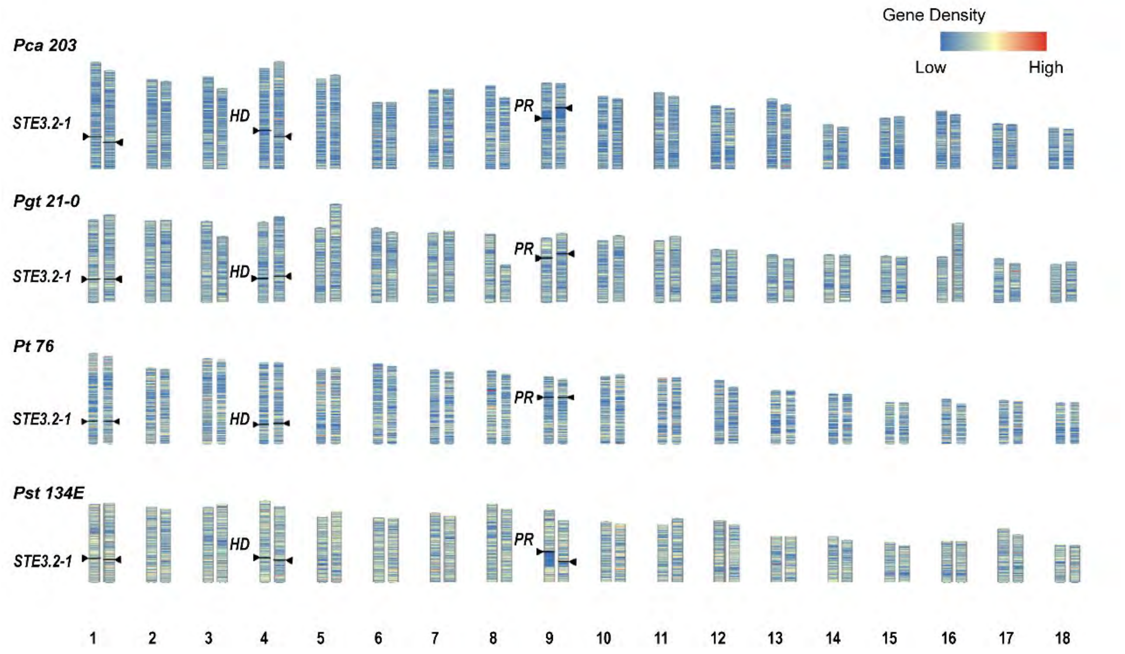


**S1 Fig. Two unlinked *MAT* loci suggest tetrapolar mating types in four *Puccinia spp*.** Karyograms of *P. coronata* f. sp. *avenae* (“*Pca 203*”), *P. graminis* f. sp. *tritici* (“*Pgt 21-0*”), *P. triticina* (“*Pt 76*”) and *P. striiformis* f. sp. *tritici* (“*Pst 134 E*”) with the positions of *HD, PR* and *STE3.2-1* loci marked by black arrow heads. *HD*, *PR* and *STE3.2-1*, located on chromosome 4, chromosome 9 and chromosome 1, respectively, suggest tetrapolar mating types in these four species.

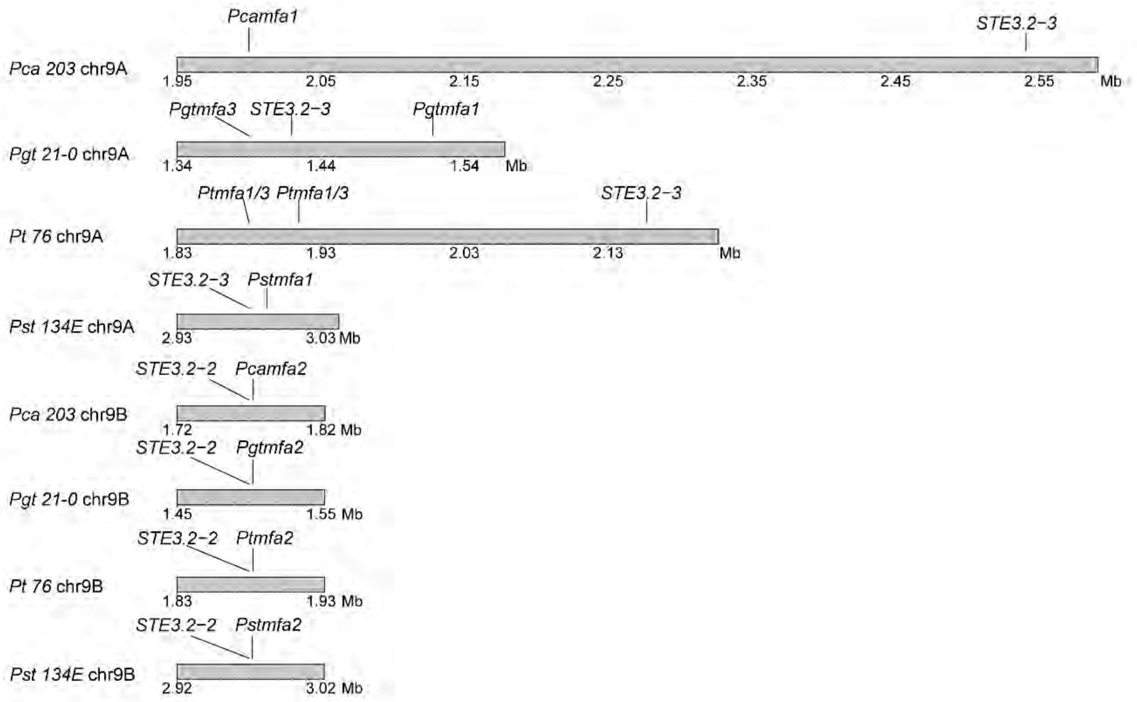


**S2 Fig. Genetic distance between pheromone precursor genes (*mfa*) and pheromone receptor (*Pra*) genes varies from 100 bp to 100 kb.** The diagrams display the location of *STE3.2-2* and *STE3.2-3* and their linked *mfa* genes on chromosome 9A and 9B in *P. coronata* f. sp. *avenae* (*"Pca 203"*), *P. graminis* f. sp. *tritici* (*"Pgt 21-0"*), *P. triticina* (*"Pt 76"*) and *P. striiformis* f. sp. *tritici* (*"Pst 134 E"*). The grey bar represents chromosome subsections containing the genes of interest with the numbers indicating the absolute location in mega base pairs on chromosome 9A and 9B. The genetic distance between *mfa1/mfa3* and *STE3.2-3* are highly variable between species, whereas *STE3.2-2* and *mfa2* are tightly linked in all species.

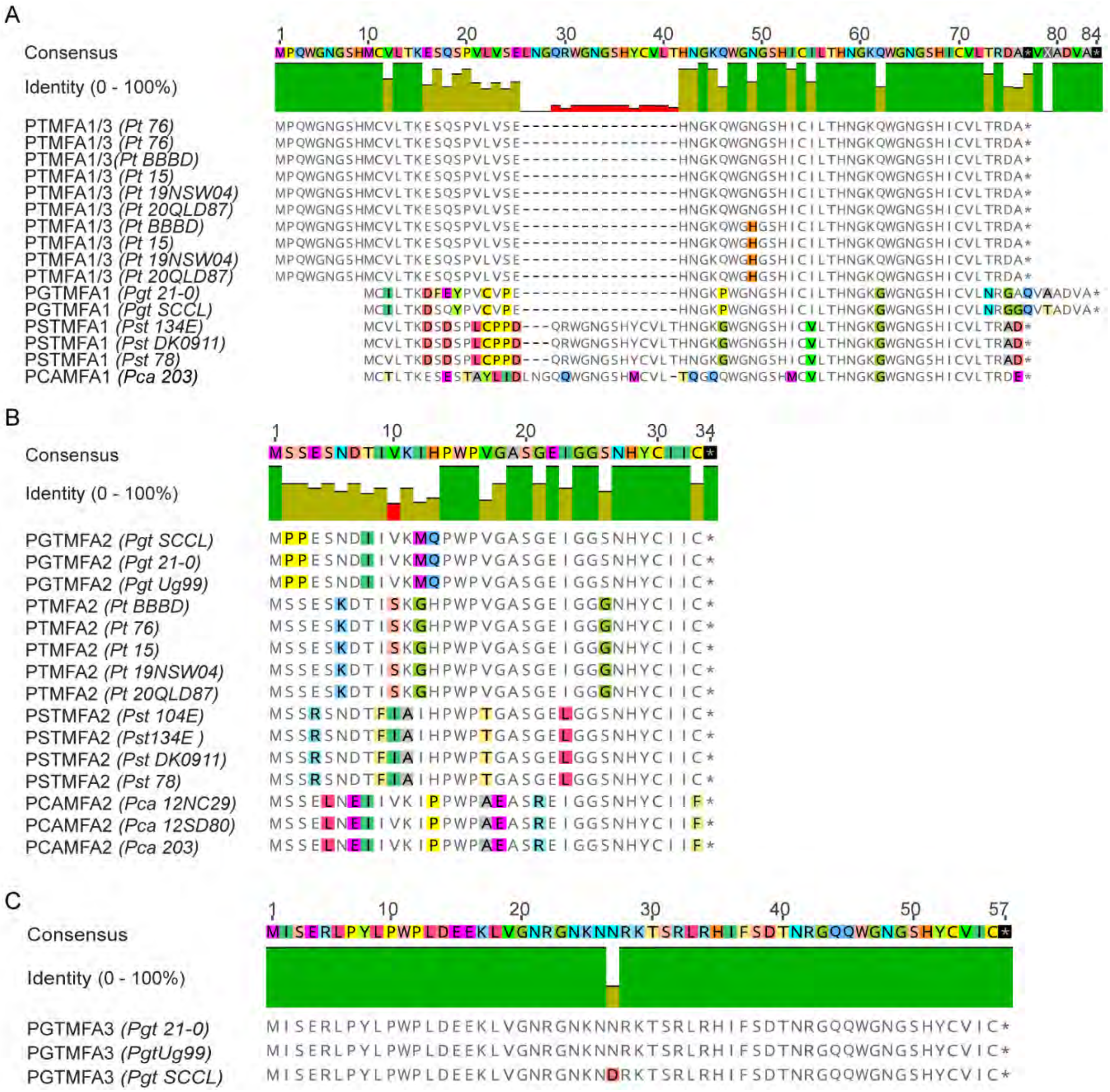


**S3 Fig. MFA protein alignments indicate mating factor a proteins are overall conserved within species.** (A) Alignment of MFA1 protein sequences from four different rust fungal species. The MFA1, which is linked to *STE3.2-3*, is mostly conserved within species. Two near identical MFA1/3 copies can be identified in four *P. triticina* isolates. In *P. graminis* f. sp. *tritici* MFA1 has four amino acid substitutions in *Pgt SCCL* versus *Pgt 21-0*. (B) Alignment of MFA2 protein sequences from four different rust fungal species. MFA2 is fully conserved within each species. (C) Alignment of MFA3 protein sequences from three *P. graminis* f. sp. *tritici* isolates. *Mfa3* is only present *P. graminis* f. sp. *tritici* in close proximity to *STE3.2-3*.

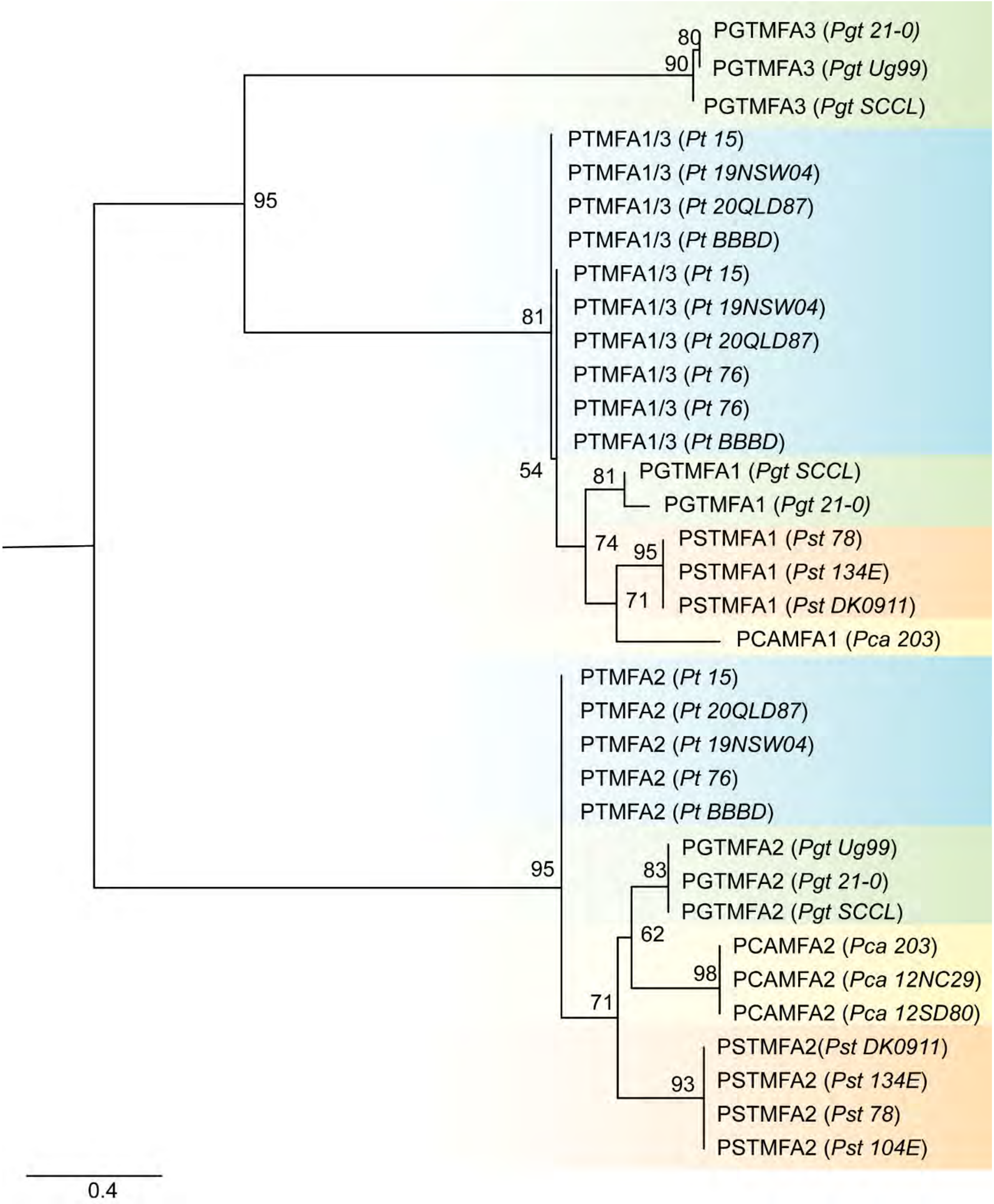


**S4 Fig. Pheromone peptide mating factor a *(*MFA) proteins group according to their proximate *Pra* alleles and display strong signals of trans-specific polymorphisms.** Maximum likelihood tree of MFA identified in four cereal rust fungi including multiple isolates per species. Tips are labelled with the species abbreviation, protein names, and isolate names are provided in parentheses. Branch support was assessed by 1000 replicates. The scale bar represents 0.4 substitutions per site. *Pca* - *P. coronata* f. sp. *avenae*, *Pgt* - *Puccinia graminis* f. sp. *tritici*, *Pt* - *P. triticina* and *Pst* - *P. striiformis* f. sp. *tritici*.

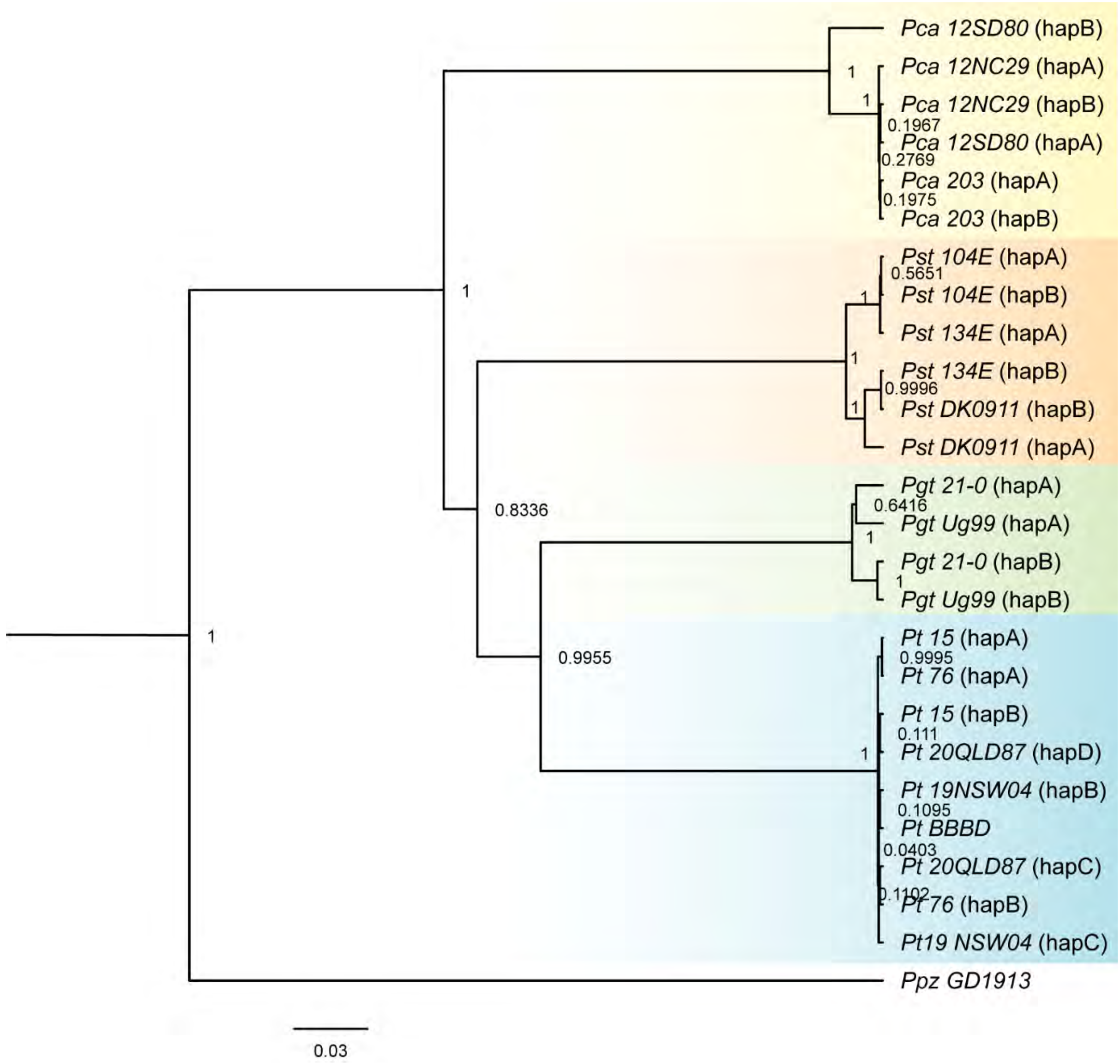


**S5 Fig. Genealogy of *STE3.2-1* in rust fungi indicates sequence of *STE3.2-1* are conserved within species.** Bayesian rooted gene tree built from *STE3.2-1* coding-based sequence alignment from four cereal rust fungi: *P. coronata* f. sp. *avenae* (*Pca*), *P. graminis* f. sp. *tritici* (*Pgt*), *P. triticina* (*Pt*) and *P. striiformis* f. sp. *tritici* (*Pst*) and *P. polysora* f. sp. *zeae* (*Ppz*) *GD1913* was included as outgroup. Trees are based on a TN93+I model of molecular evolution. Each node is labelled with its values of posterior probability (PP). PP values above 0.95 are considered to have strong evidence for monophyly of a clade and PP values of identical alleles are not displayed. The scale bar represents the number of nucleotide substitutions per site. Alleles of the same species are colored with identical background: *Pca* (yellow), *Pgt* (green), *Pt* (blue), *Pst* (orange).

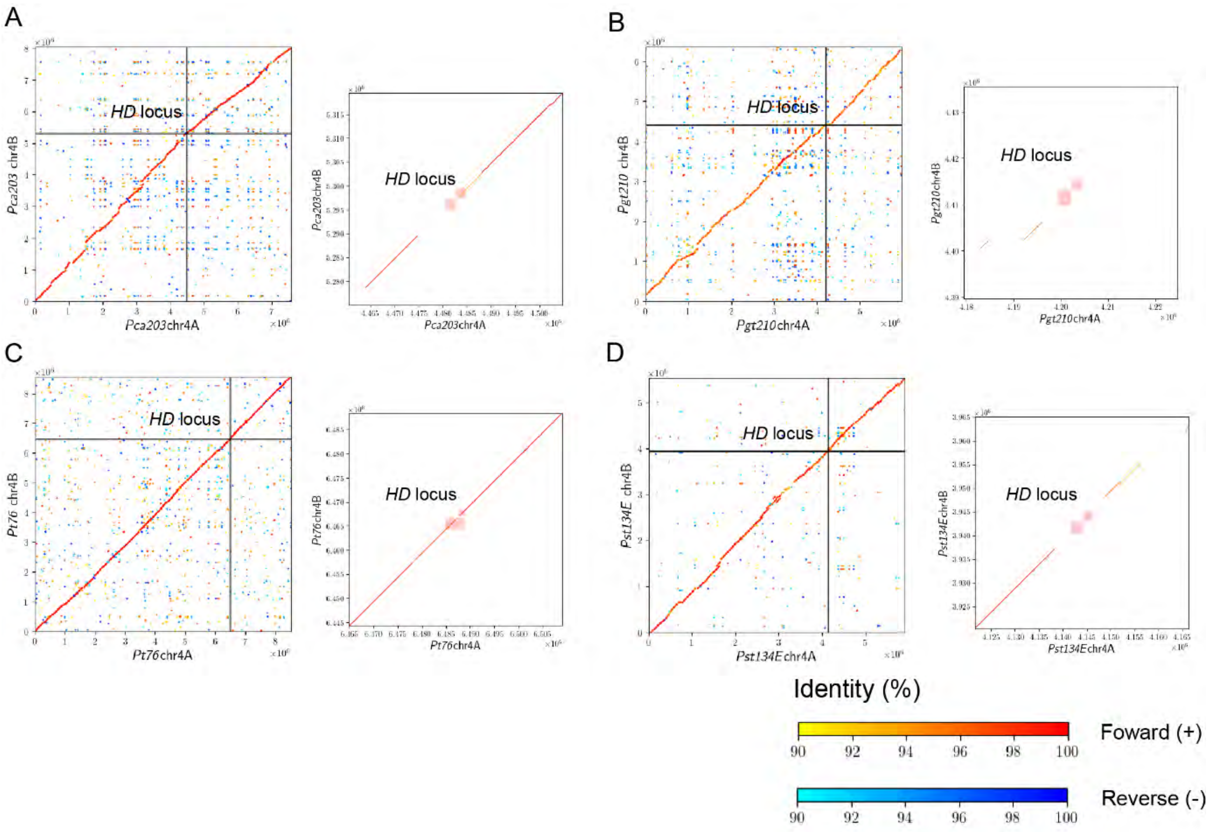


**S6 Fig. Whole chromosome alignments of *HD* loci containing chromosomes between the two haplotypes of each dikaryotic genome exhibit slightly reduced synteny around *HD* locus.** The figure shows dots plots of whole chromosome alignments between the two *HD* loci containing chromosomes from dikaryotic genome assemblies. Each panel consists of dot plots of the whole chromosome and subset dot plot zooming into the *HD* locus. The *HD* locus is labelled and line colors show the nucleotide percentage identity and nucleotide orientation as indicated in the figure legend. Subfigure A to D show *P. coronata* f. sp. *avenae* ("*Pca 203*"), *P. graminis* f. sp. *tritici* ("*Pgt 21-0*"), *P. triticina* ("*Pt 76*") and *P. striiformis* f. sp. *tritici* ("*Pst 134 E*"), respectively.

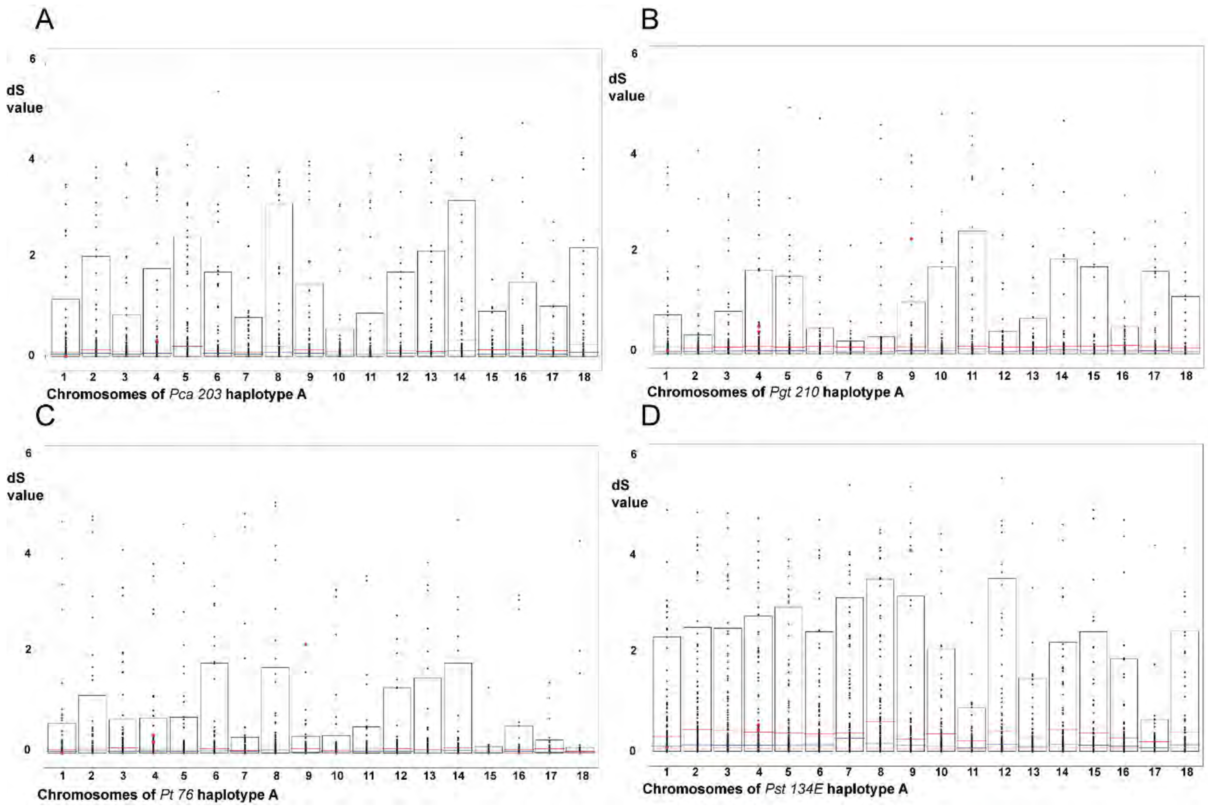


**S7 Fig. Allele pairs on chromosomes 4 and 9 are not overall more diverged when compared to other chromosomes.** The plots show the distribution of $d_S$ values of allele pairs on sister chromosomes in four cereal rust fungal species. Bar plots show the distribution of $d_S$ values up to the 99% quantile. Red lines represent threshold of $d_S$ values of 95% of all alleles. Blue lines represent threshold of $d_S$ values of 90% of all alleles. Black points show $d_S$ values of individual allele pairs. *HD* and *STE3* allele pairs are highlighted as red points. Subfigures A to D show *P. coronata* f. sp. *avenae* (“*Pca 203*”), *P. graminis* f. sp. *tritici* (“*Pgt 21-0*”), *P. triticina* (“*Pt 76*”) and *P. striiformis* f. sp. *tritici* (“*Pst 134 E*”), respectively.

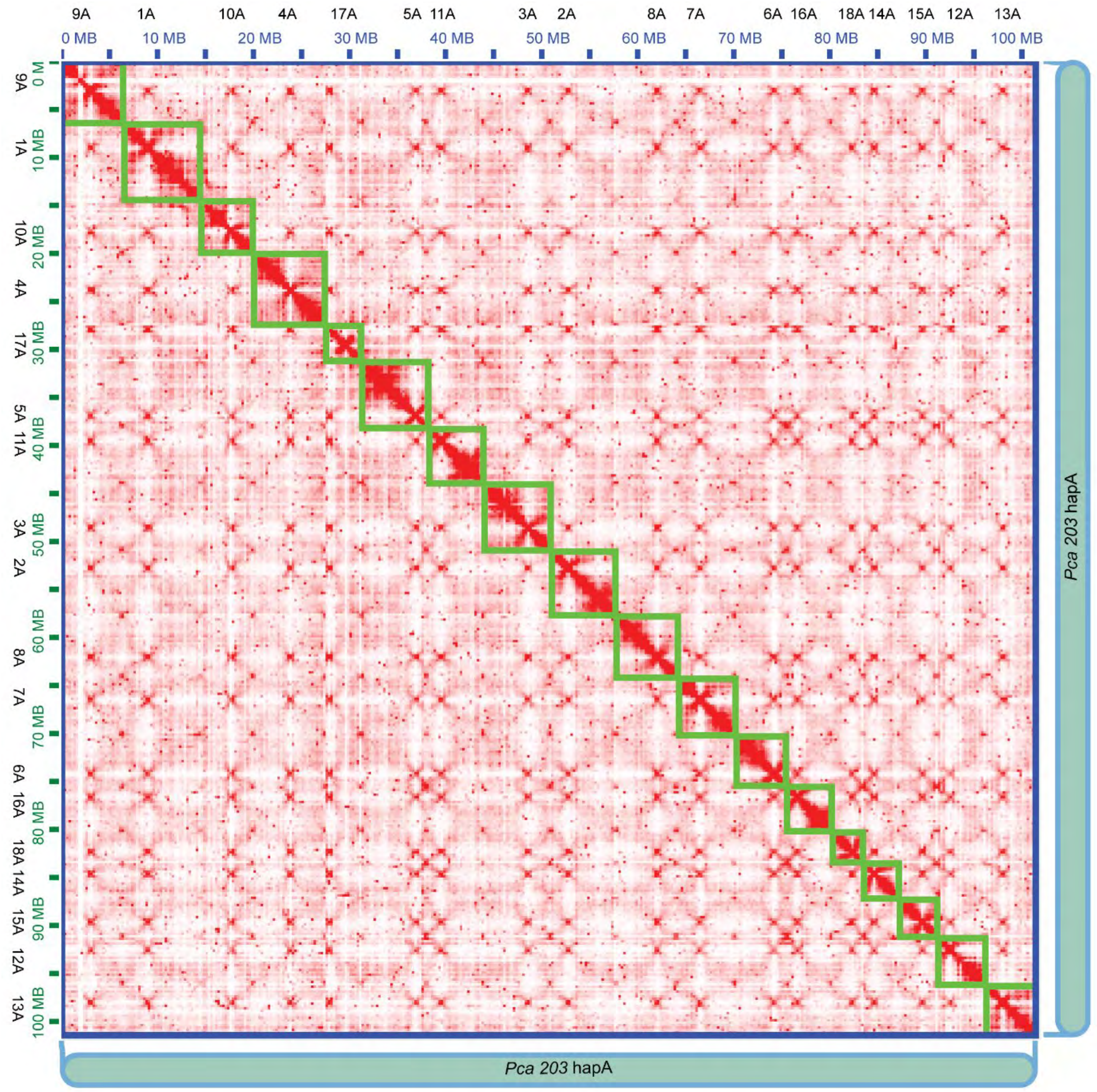


**S8 Fig. Hi-C heatmap of *P. coronata* f. sp. *avenae* ("*Pca 203*") hapA.**

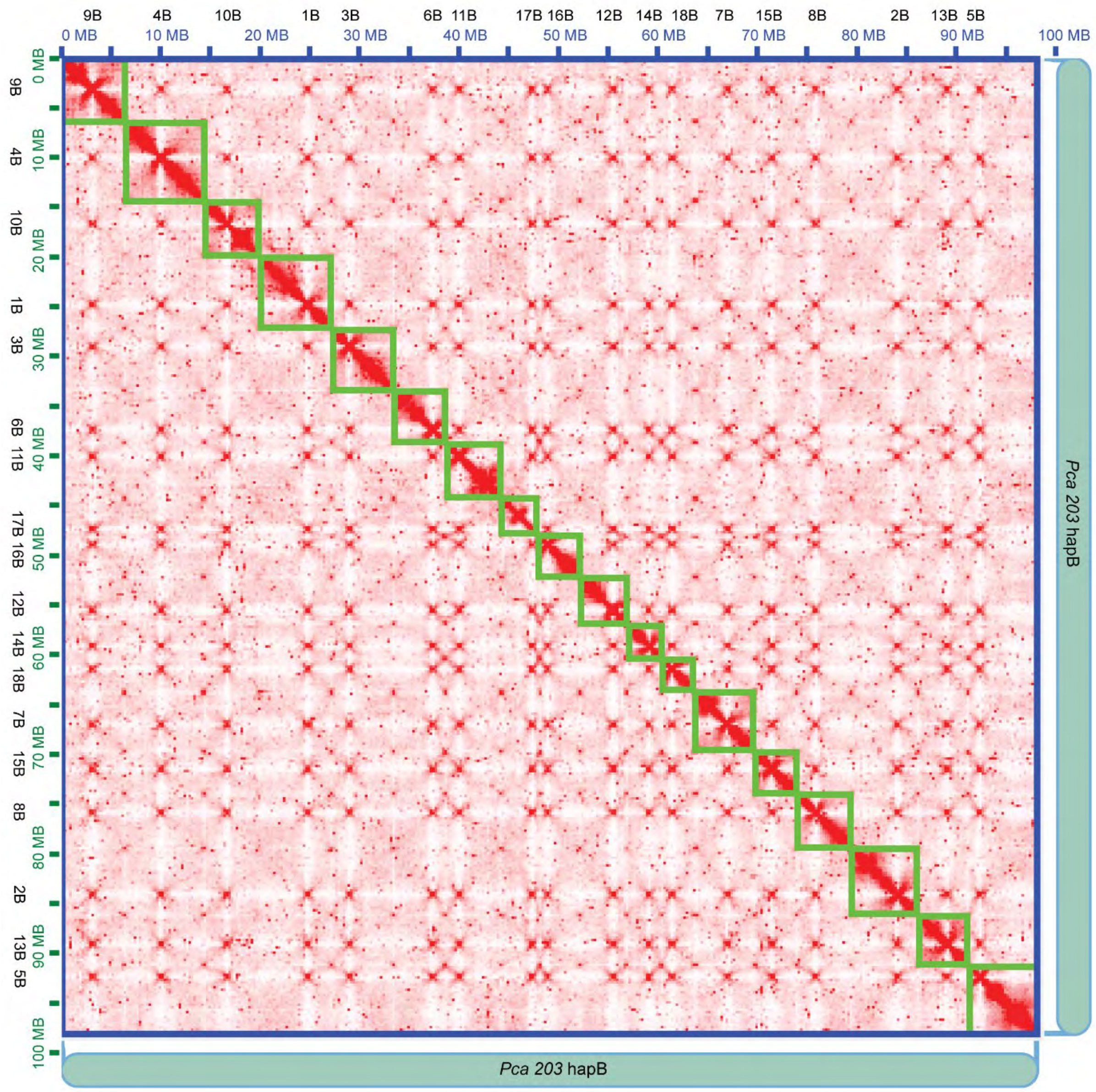


**S9 Fig. Hi-C heatmap of *P. coronata* f. sp. *avenae* ("*Pca 203*") hapB.**

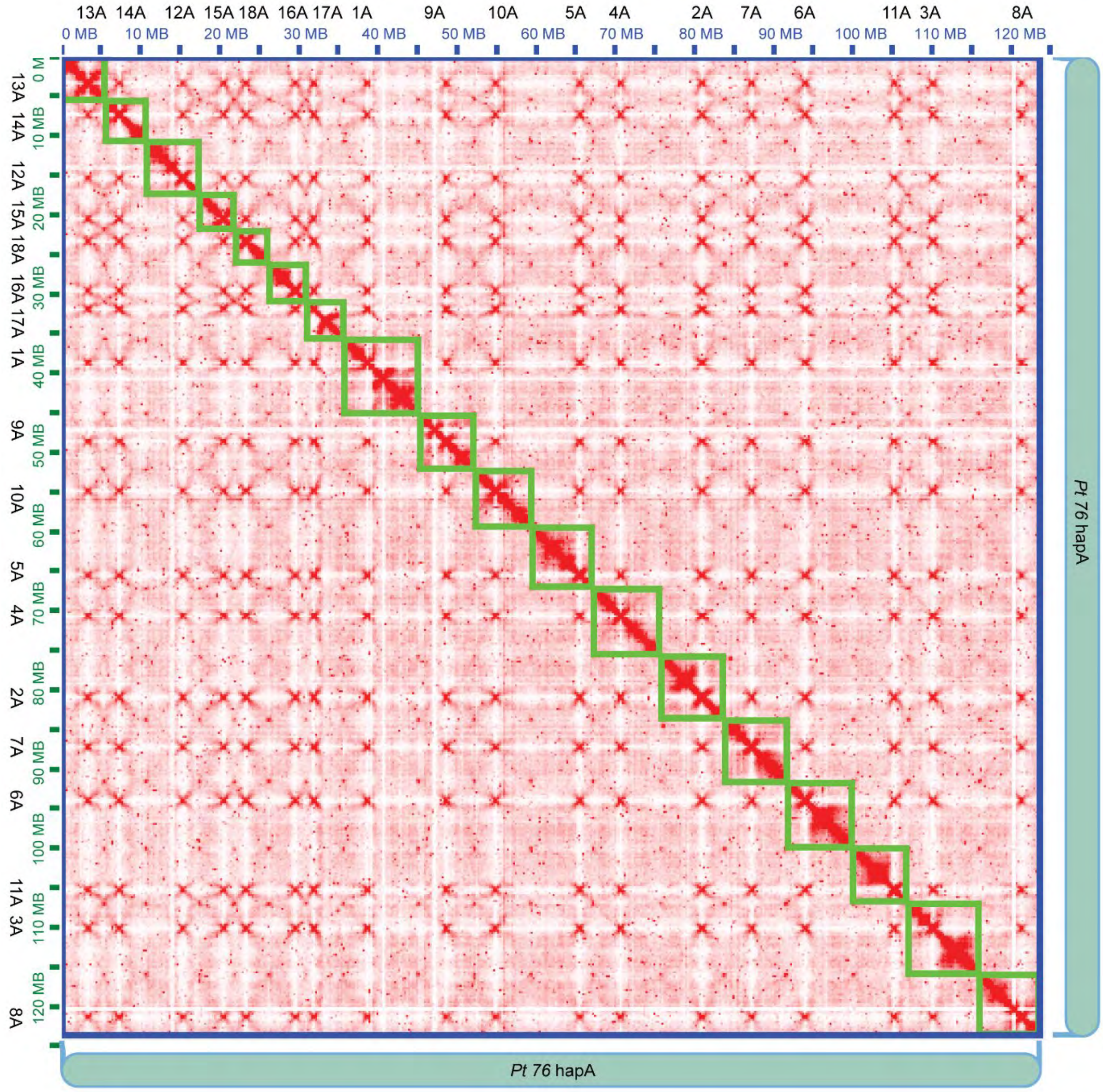


**S10 Fig. Hi-C heatmap of *P. triticina* ("*Pt 76*") hapA.**

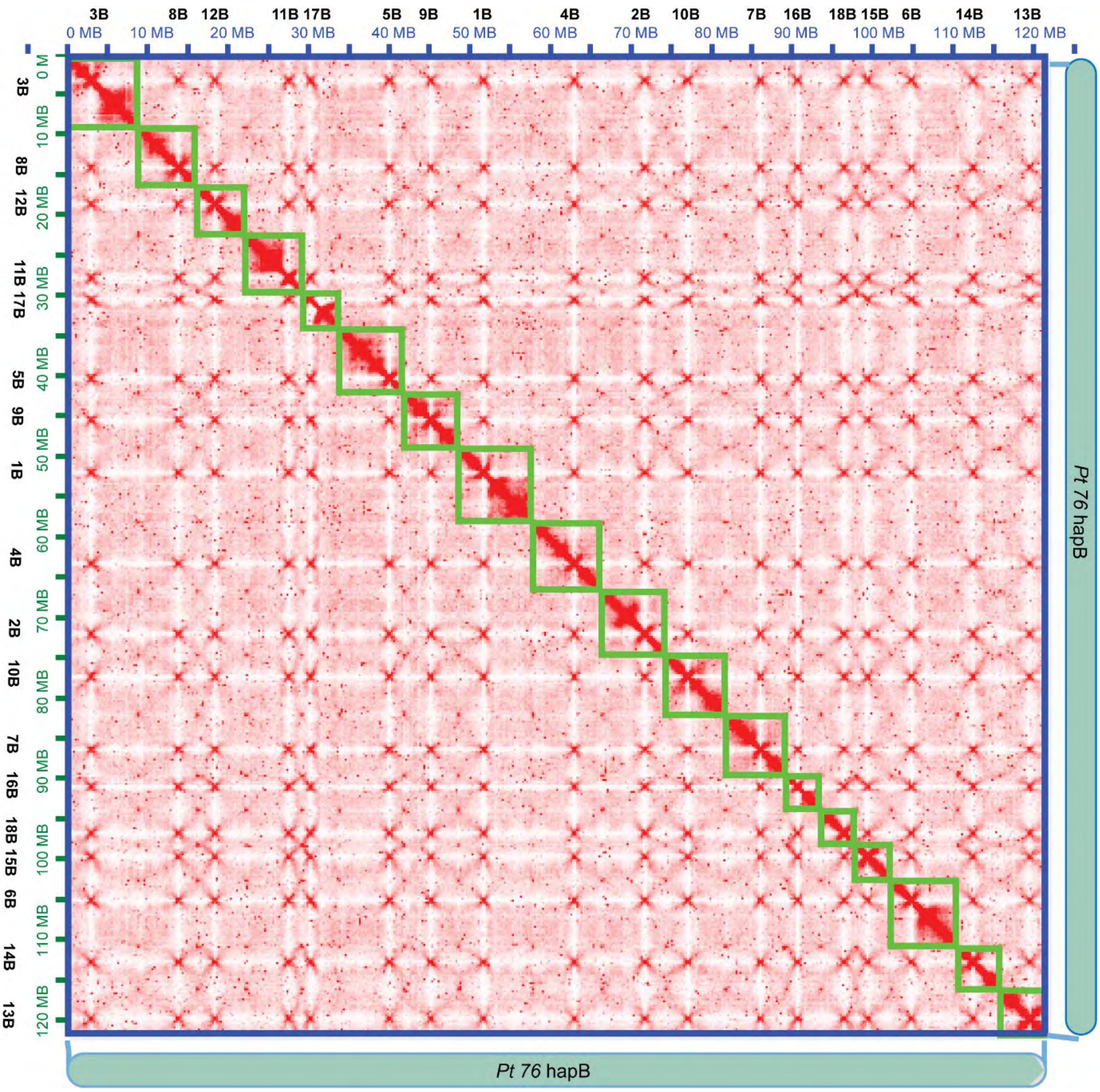


**S11 Fig. Hi-C heatmap of *P. triticina* ("*Pt 76*") hapB.**

108

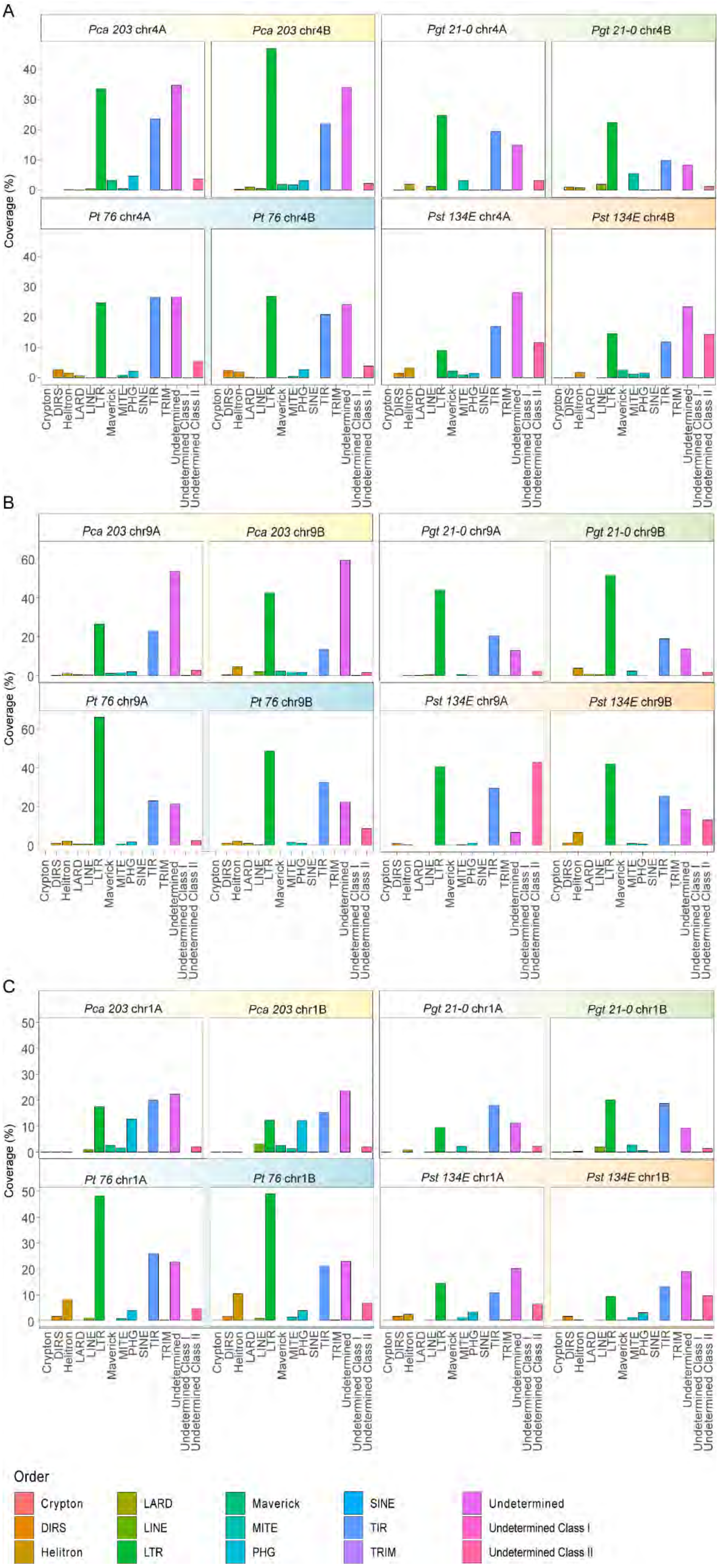
A
Pca 203 chr4A
Pca 203 chr4B
Pgt 21-0 chr4A
Pgt 21-0 chr4B
Pt 76 chr4A
Pt 76 chr4B
Pst 134E chr4A
Pst 134E chr4B
B
Pca 203 chr9A
Pca 203 chr9B
Pgt 21-0 chr9A
Pgt 21-0 chr9B
Pt 76 chr9A
Pt 76 chr9B
Pst 134E chr9A
Pst 134E chr9B
C
Pca 203 chr1A
Pca 203 chr1B
Pgt 21-0 chr1A
Pgt 21-0 chr1B
Pt 76 chr1A
Pt 76 chr1B
Pst 134E chr1A
Pst 134E chr1B
Coverage (%)
Order
Crypton
DIRS
Helitron
LARD
LINE
LTR
Maverick
MITE
PHG
SINE
TIR
TRIM
Undetermined
Undetermined Class I
Undetermined Class II

**S12 Fig. Coverage of different transposable element orders at the *HD, PR,* and *STE3.2-1* locus.** The plots show the percentage of nucleotides covered by different transposable element orders at (A) *HD* locus (B) *PR* locus (C) *STE3.2-1* locus. Each subfigure A to C shows the coverage in each haplotype of the dikaryotic genomes of *P. coronata* f. sp. *avenae* (*"Pca 203"*), *P. graminis* f. sp. *tritici* (*"Pgt 21-0"*), *P. triticina* (*"Pt 76"*) and *P. striiformis* f. sp. *tritici* (*"Pst 134 E"*). Different TE orders are color coded as shown in the legend. TEs with no assigned class are labelled "Undetermined". TEs with no assigned order but belonging to Class I (RNA retrotransposons) or Class II (DNA transposons) are labelled "Undetermined Class I" or "Undetermined Class II", respectively.

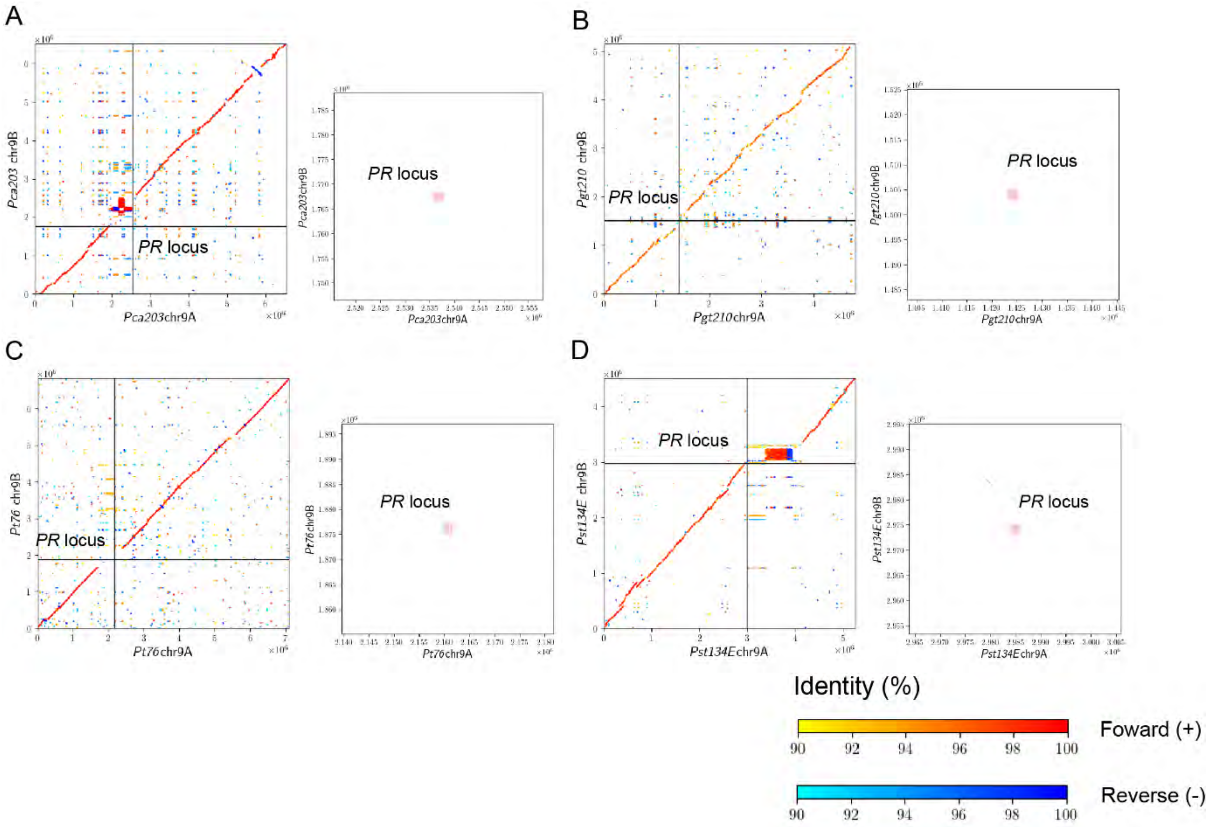


**S13 Fig. Whole chromosome alignments of *PR* loci containing chromosomes between two haplotypes of each dikaryotic genome exhibit strong signs of synteny loss.** The figure shows dot plots of whole chromosome alignments between the two *PR* loci containing chromosomes from the dikaryotic genome assemblies. Each panel consists of dot plots of the whole chromosome and subset dot plots zooming into the *PR* locus. The *PR* locus is labelled and line colors show the nucleotide percentage identity and nucleotide orientation as indicated in the figure legend. Subfigures A to D show *P. coronata* f. sp. *avenae* (“*Pca 203*”), *P. graminis* f. sp. *tritici* (“*Pgt 21-0*”), *P. triticina* (“*Pt 76*”) and *P. striiformis* f. sp. *tritici* (“*Pst 134 E*”), respectively.

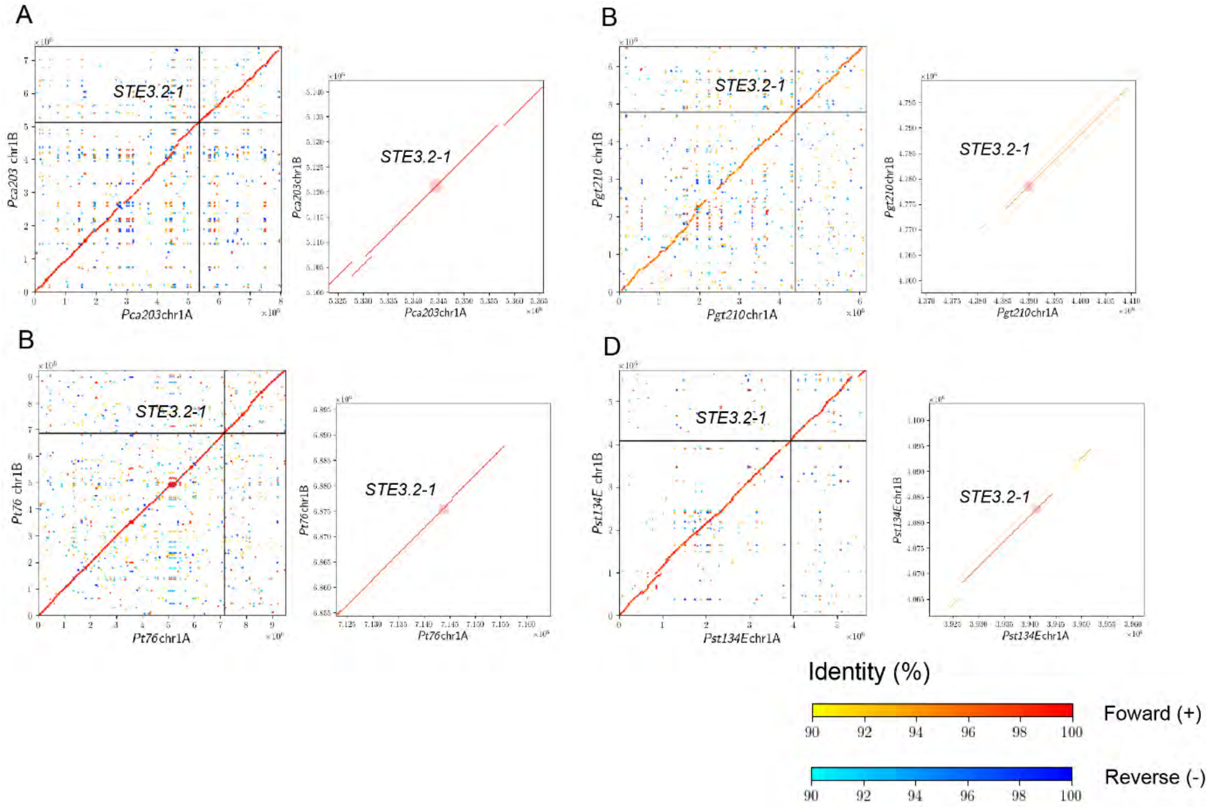


**S14 Fig. Whole chromosome alignments of *STE3.2-1* loci containing chromosomes between two haplotypes of each dikaryotic genome suggest high conservation of *STE3.2-1*.** The figure shows dot plots of whole chromosome alignments between the two *STE3.2-1* loci containing chromosomes from dikaryotic genome assemblies. Each panel consists of dot plots of the whole chromosome and subset dot plots zooming into the *STE3.2-1* locus. The *STE3.2-1* locus is labelled and line colors show the nucleotide percentage identity and nucleotide orientation as indicated in the figure legend. Subfigures A to D show *P. coronata* f. sp. *avenae* (“*Pca 203*”), *P. graminis* f. sp. *tritici* (“*Pgt 21-0*”), *P. triticina* (“*Pt 76*”) and *P. striiformis* f. sp. *tritici* (“*Pst 134 E*”), respectively.

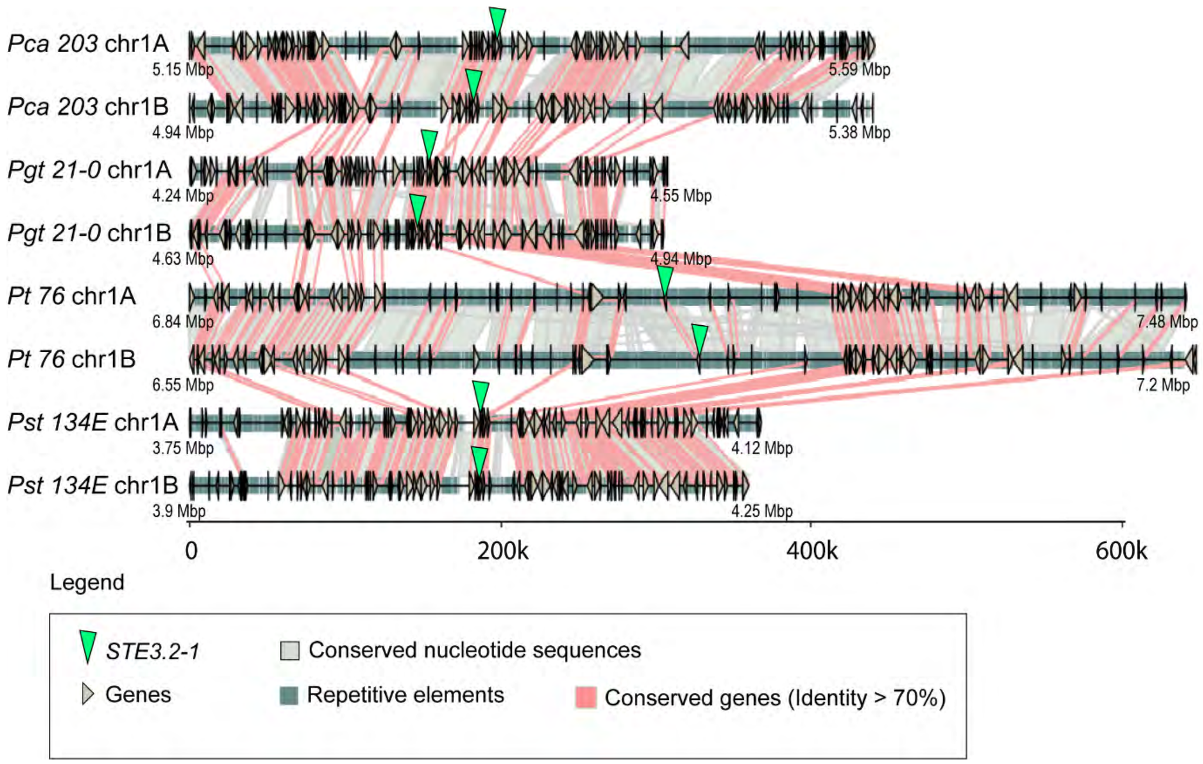


**S15 Fig. Synteny analysis of *STE3.2-1* loci and their flanking regions reveal no evidence of genetic degeneration.** Synteny graphs of the *STE3.2-1* locus including proximal regions in *P. coronara* f. sp. *avenae* ("*Pca 203*"), *P. gramminis* f. sp. *tritici* ("*Pgt 21-0*"), *P. triticina* ("*Pt 76*"), and *P. striiformis* f. sp. *tritici* ("*Pst 134E*"). Proximal regions are defined as 40 genes downstream and upstream of the *STE3.2-1* alleles, respectively. *STE3.2-1* loci are highly syntenic within each dikaryotic genome and conserved between species. Red lines between chromosome sections represent gene pairs with sequence identity higher than 70% and grey shades represent conserved nucleotide sequences (>=1000 bp and identity >=90%). For additional annotations please refer to the provided legend ("Legend").

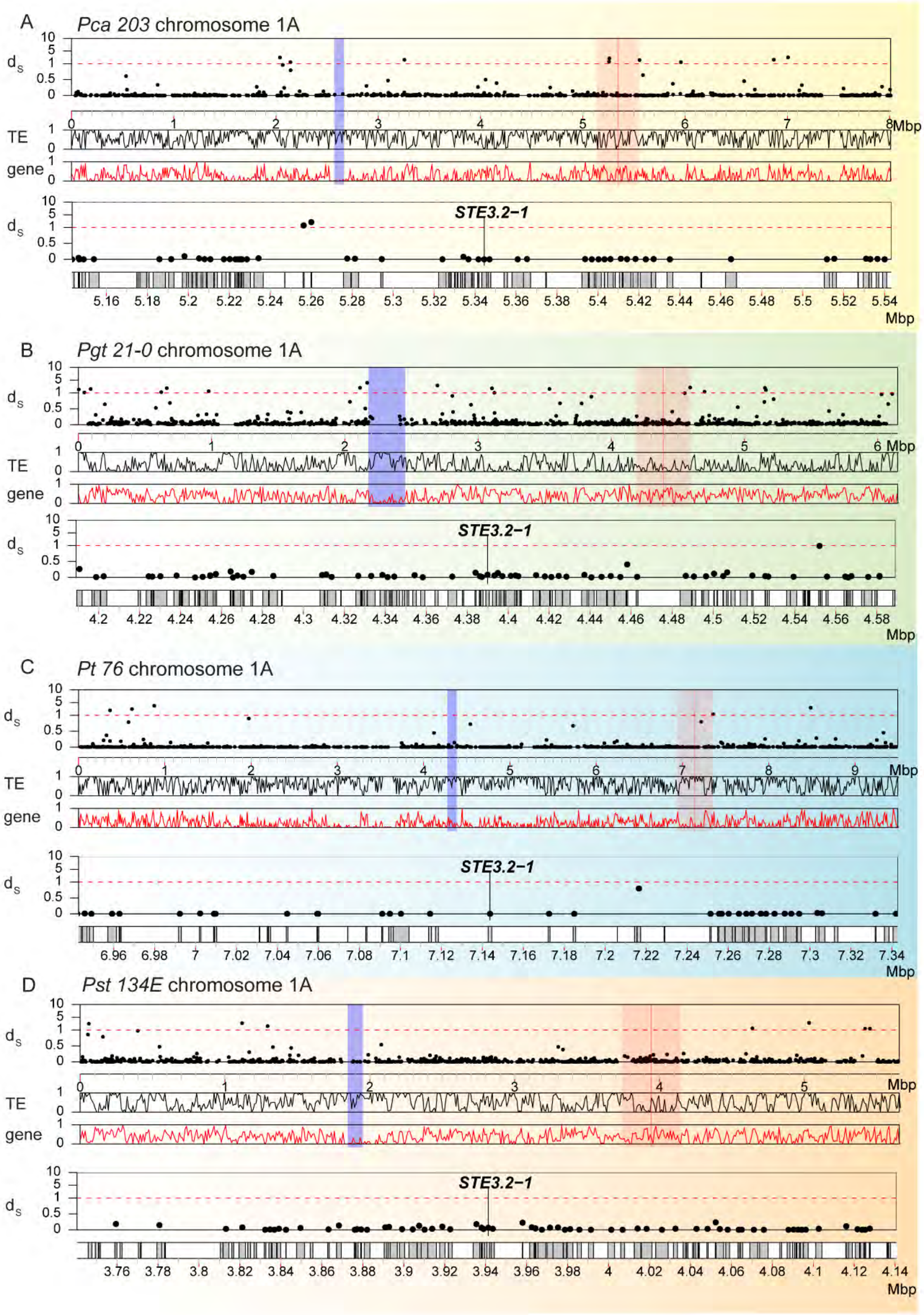


**S16 Fig. Synonymous divergence ($d_S$) values between *STE3.2-1* alleles along chromosome 1A suggest conservation of this locus.** Synonymous divergence values ($d_S$) for all allele pairs are plotted along chromosome 1A for (A) *P. coronara* f. sp. *avenae* (“*Pca 203*”), (B) *P. gramminis* f. sp. *tritici* (“*Pgt 21-0*”), (C) *P. triticina* (“*Pt 76*”), and (D) *P. striiformis* f. sp. *tritici* (“*Pst 134E*”). In each panel, the top track shows the $d_S$ values (“$d_S$”) of allele pairs along chromosome 1. Each dot corresponds to the $d_S$ value of a single allele pair. The second and third track show the averaged TE (“TE”) and gene (“gene”) density along chromosome 1 in 10 kbp-sized windows, respectively. The *STE3.2-1* alleles are highlighted with a red line and red shading indicates a 0.4 mbp-sized window around the *STE3.2-1* locus. Predicted centromeric regions are marked with blue shading. The two lower tracks ($d_s$ values and gene locations) provide a detailed zoomed in view of red shaded area around the *STE3.2-1* locus. Species-specific background coloring is the same as for Figure 1.

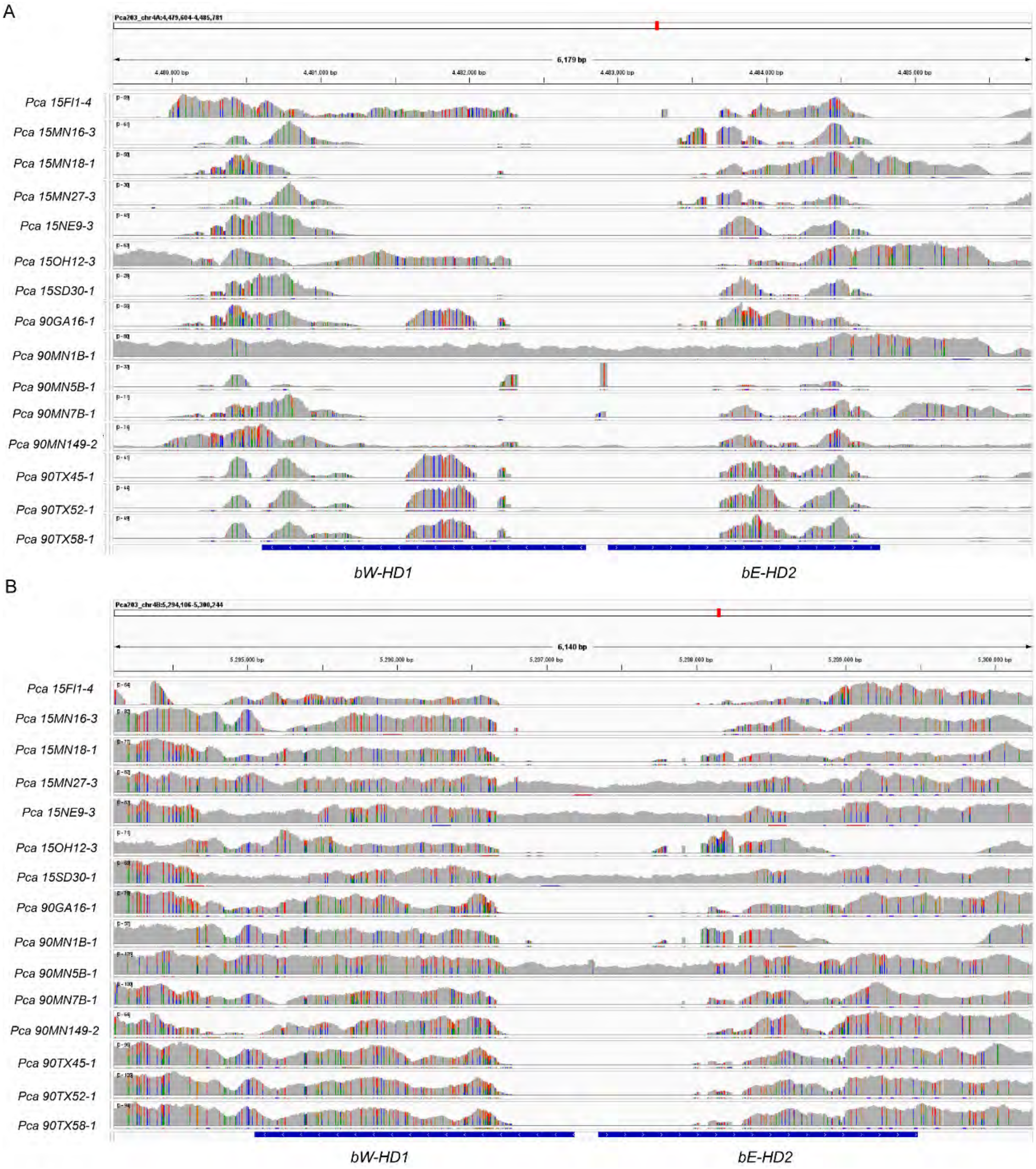


**S17 Fig. IGV screen shots of short read Illumina mapping of various *P. coronata* f. sp. *avenae* isolates against the *HD* locus in the *Pca 203* reference.**

(A) shows mapping against the *HD* locus on chromosome 4A and (B) chromosome 4B, respectively.

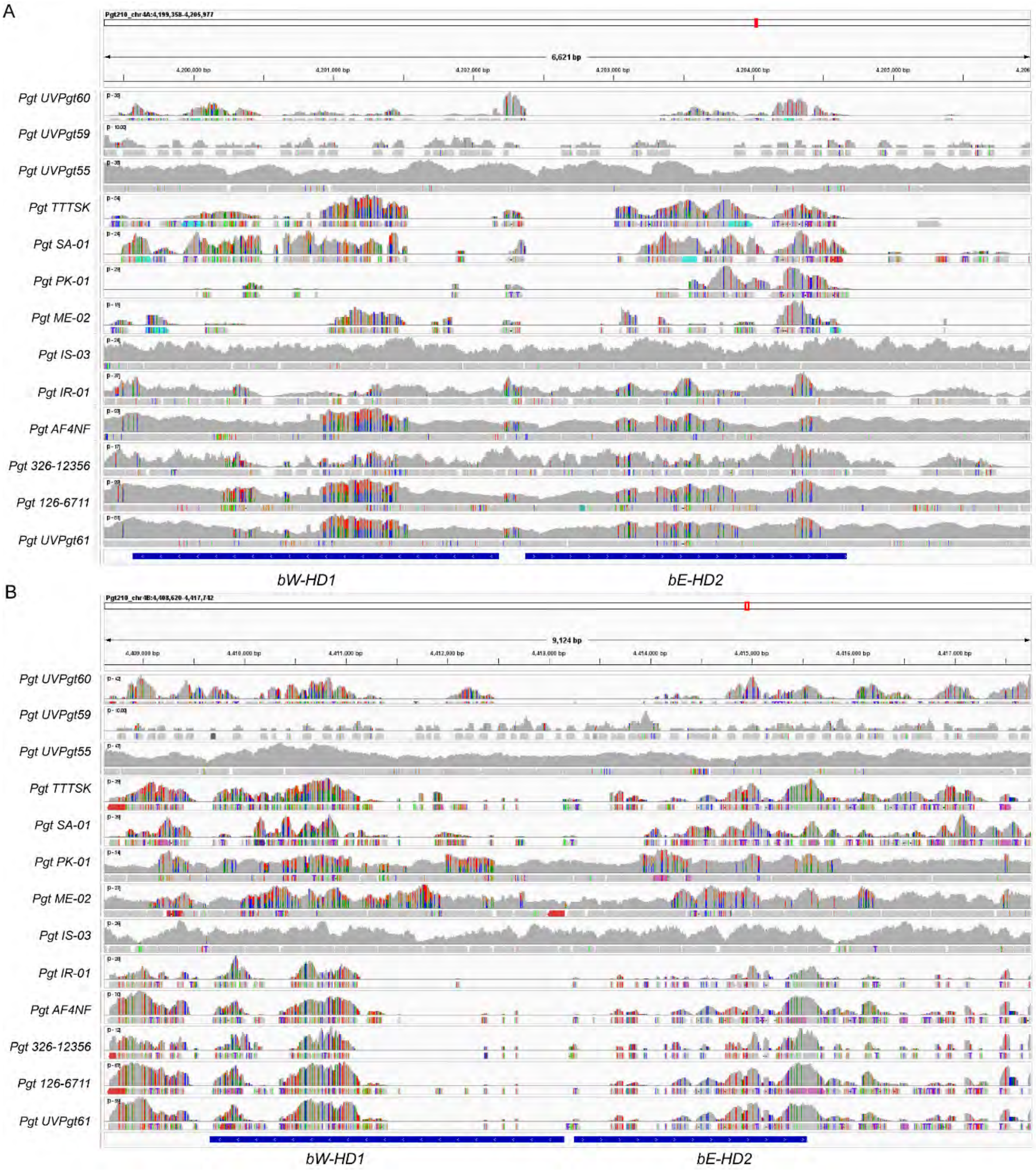


**S18 Fig. IGV screen shot of short read Illumina mapping of various *P. gramminis* f. sp. *tritici* isolates against the *HD* locus in the *Pgt 21-0* reference.**

(A) shows mapping against the *HD* locus on chromosome 4A and (B) chromosome 4B, respectively.

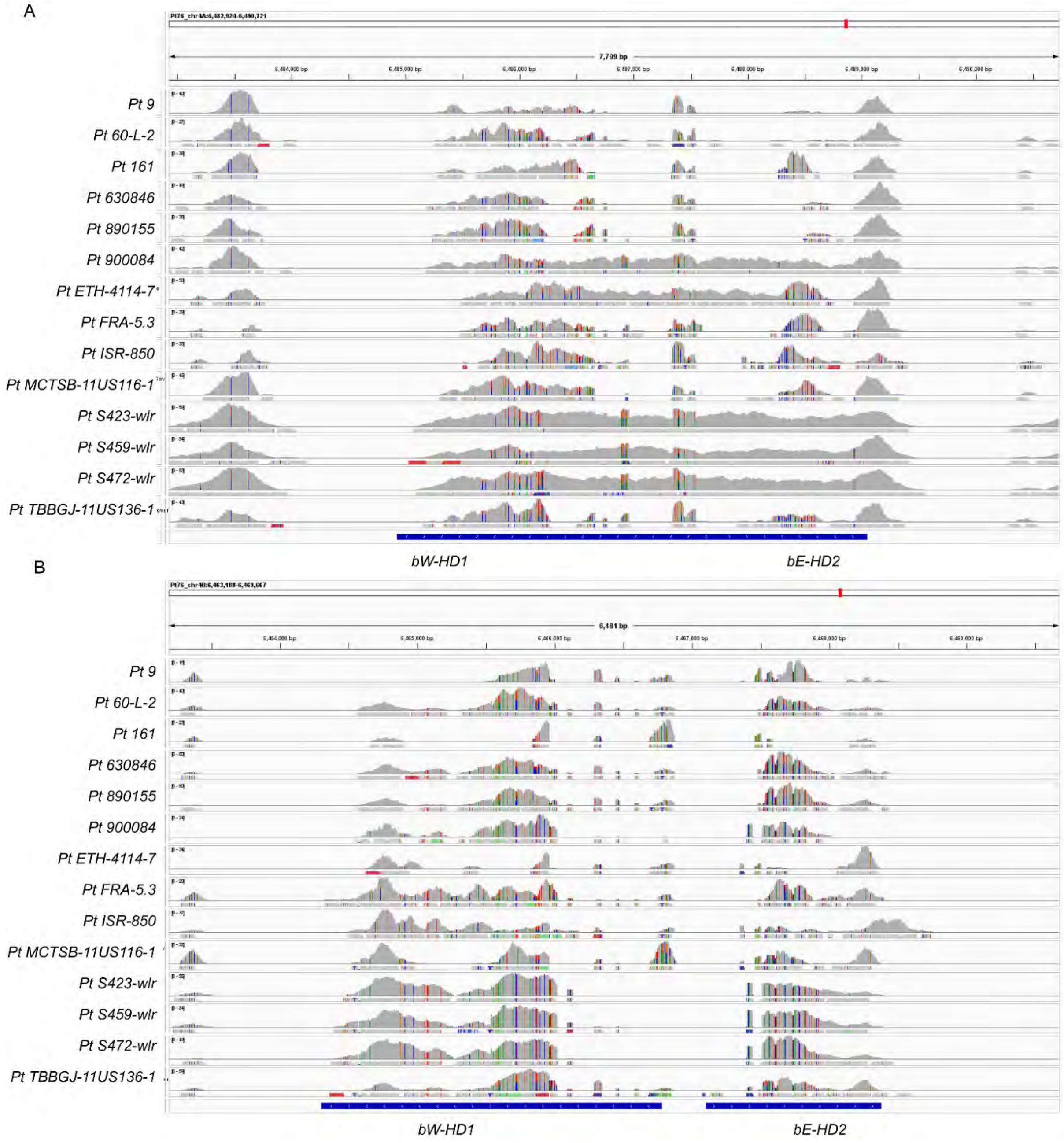


**S19 Fig. IGV screen shots of short read Illumina mapping of various *P. triticina* against the *HD* isolates locus in the *Pt 76* reference.**

(A) shows mapping against the *HD* locus on chromosome 4A and (B) chromosome 4B, respectively.

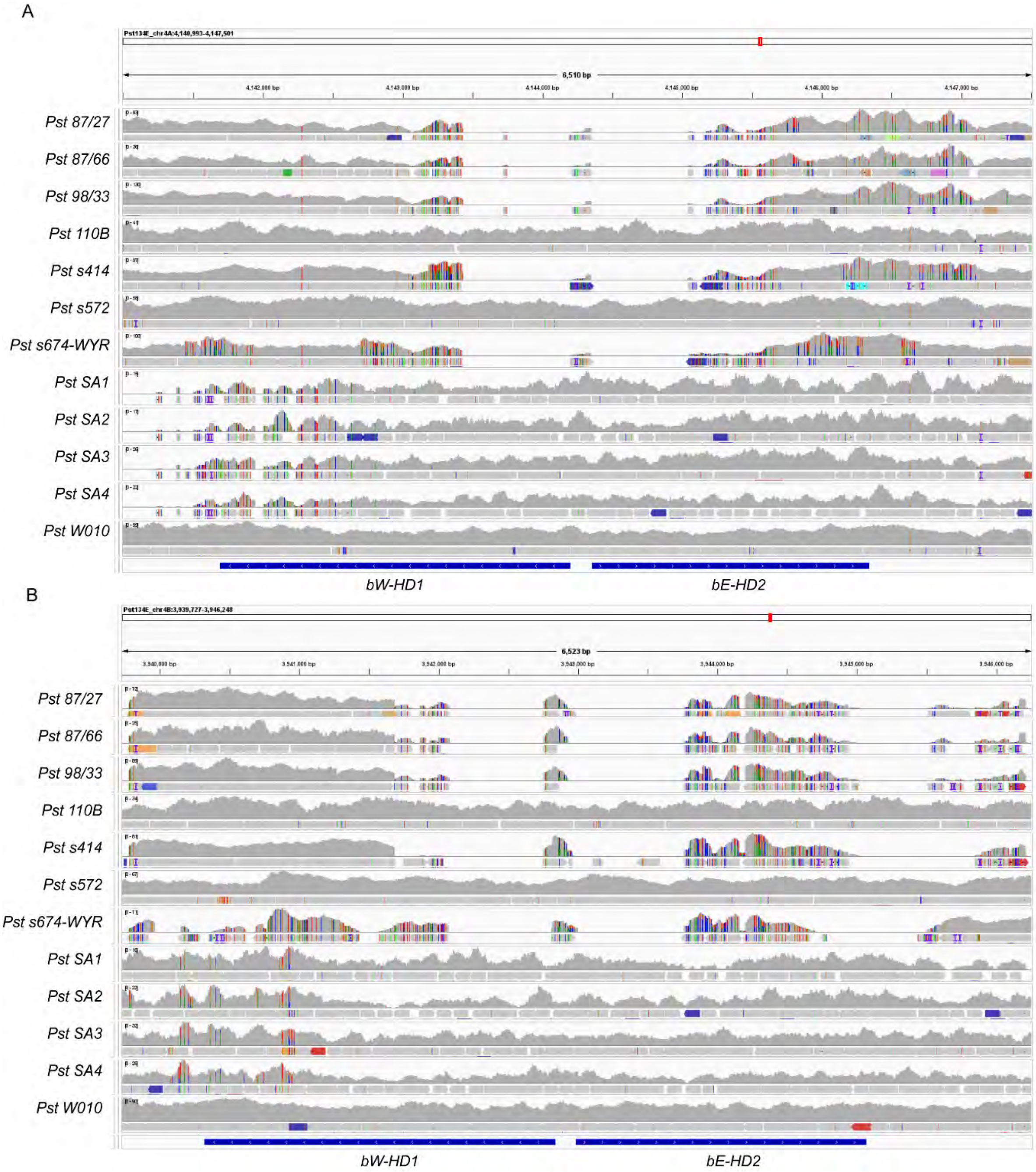


**S20 Fig. IGV screen shots of short read Illumina mapping of various *P. striiformis* f. sp. *tritici* isolates against the *HD* locus in the *Pst 134E* reference.**

(A) shows mapping against the *HD* locus on chromosome 4A and (B) chromosome 4B, respectively.

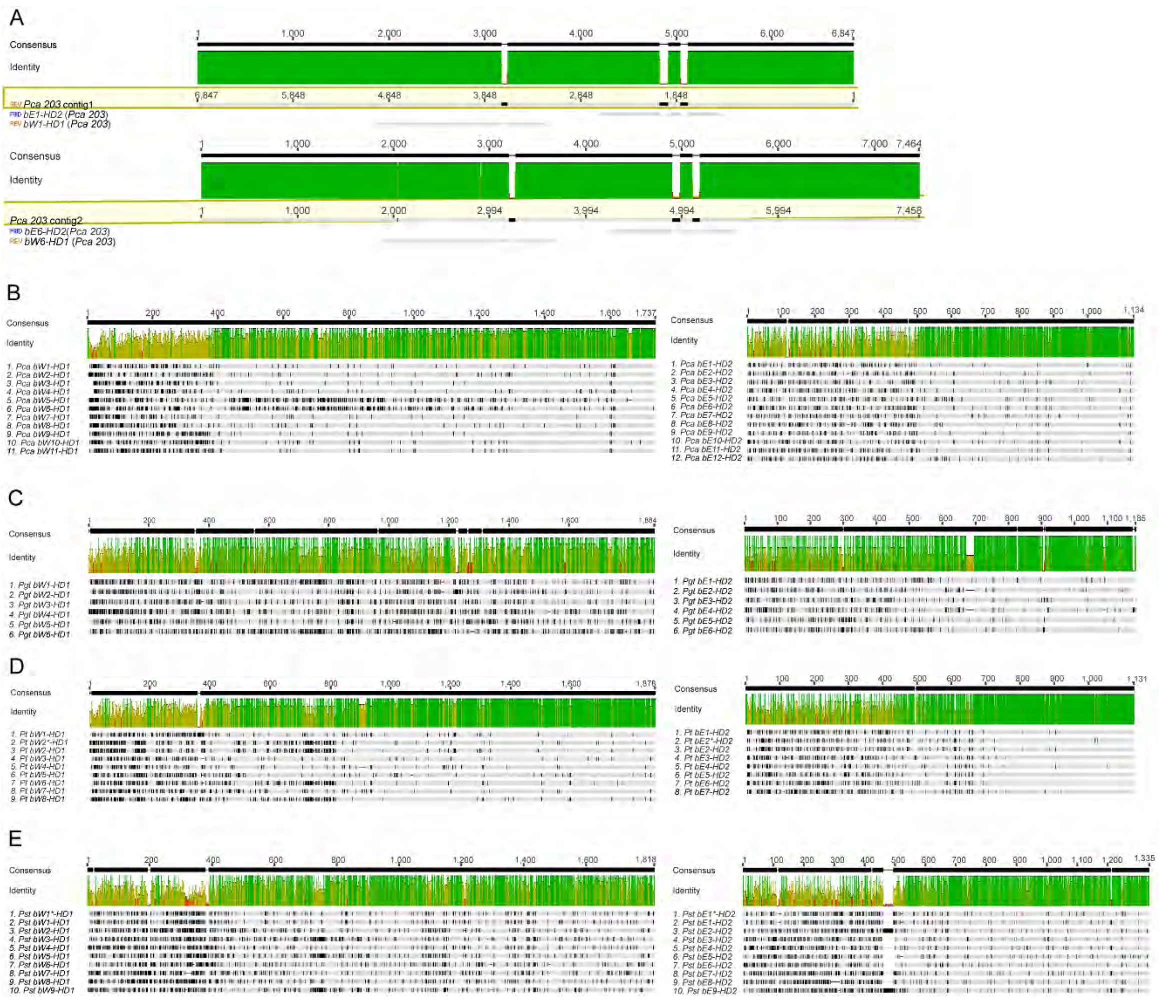


**S21 Fig. Nucleotide alignments of the *HD* gene coding sequence in four cereal rust fungi indicate *bW-HD1* and *bE-HD2* are multiallelic in each species.** (A) Nucleotide alignment of the *de novo* reconstructed *HD* locus from *Pca 203* Illumina short read data with the coding sequences of *bW-HD1* and *bE-HD2* alleles from *Pca 203* dikaryotic reference genome. (B) to (D) multiple sequence alignments of *de novo* reconstructed *HD* gene coding sequences, (E) multiple sequence alignments of *Pst bW-HD1* and *bE-HD2* alleles (15). In each subfigure (B) to (E), the top two track shows the consensus sequence length and relative sequence identity, respectively. Subfigure (B) to (E) show *P. coronata* f. sp. *avenae* (“*Pca*”), *P. graminis* f. sp. *tritici* (“*Pgt*”), *P. triticina* (“*Pt*”) and *P. striiformis* f. sp. *tritici* (“*Pst*”), respectively. The *bW-HD1* and *bE-HD2* are numbered in accordance with Fig 1.

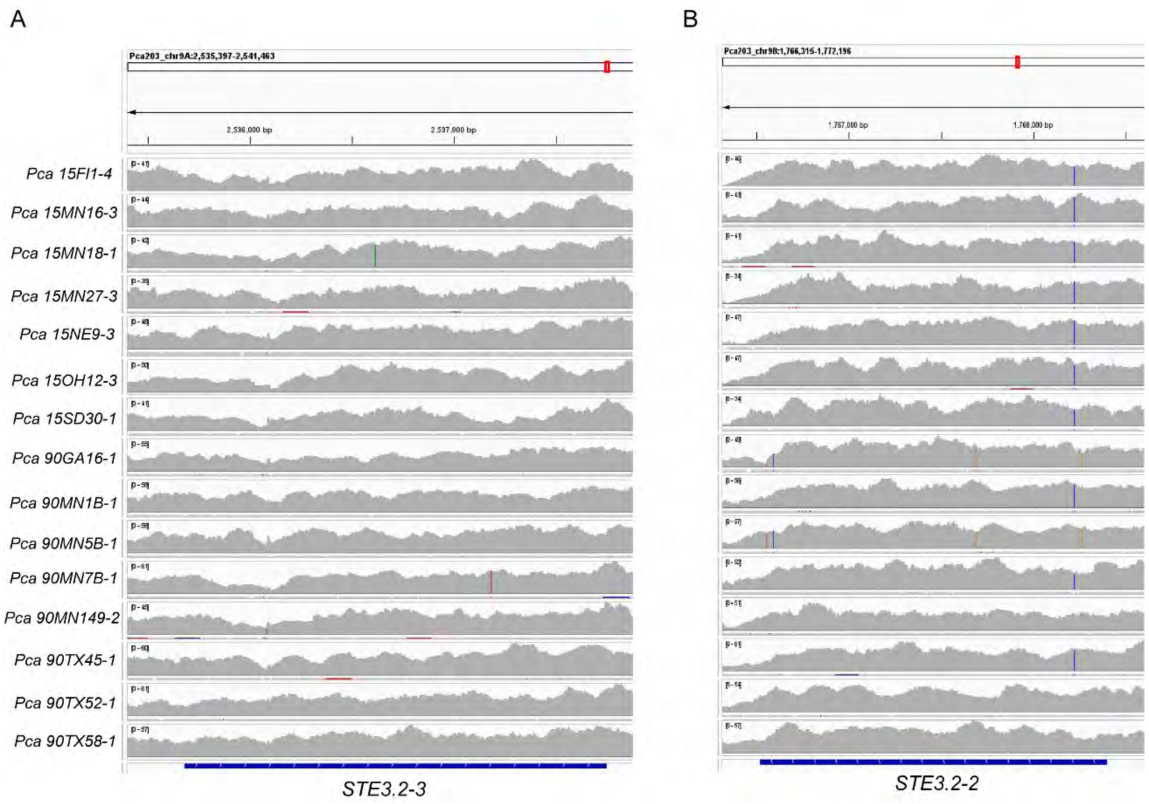


**S22 Fig. IGV screen shots of short read Illumina mapping of various *P. coronata* f. sp. *avenae* isolates against the *PR* locus in the *Pca 203* reference.**

(A) shows mapping against the *PR* locus on chromosome 9A and (B) chromosome 9B, respectively.

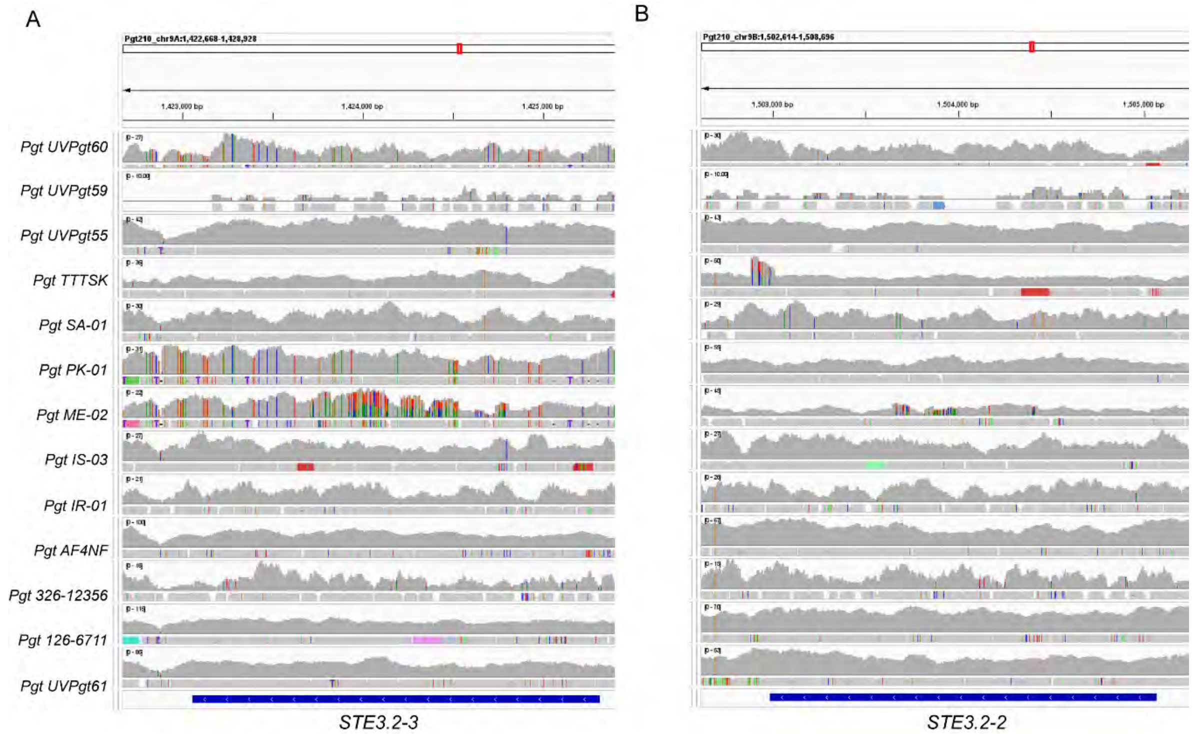


**S23 Fig. IGV screen shot of short read Illumina mapping of various *P. graminis* f. sp. *tritici* isolates against the *PR* locus in the *Pgt 21-0* reference.**

(A) shows mapping against the *PR* locus on chromosome 9A and (B) chromosome 9B, respectively.

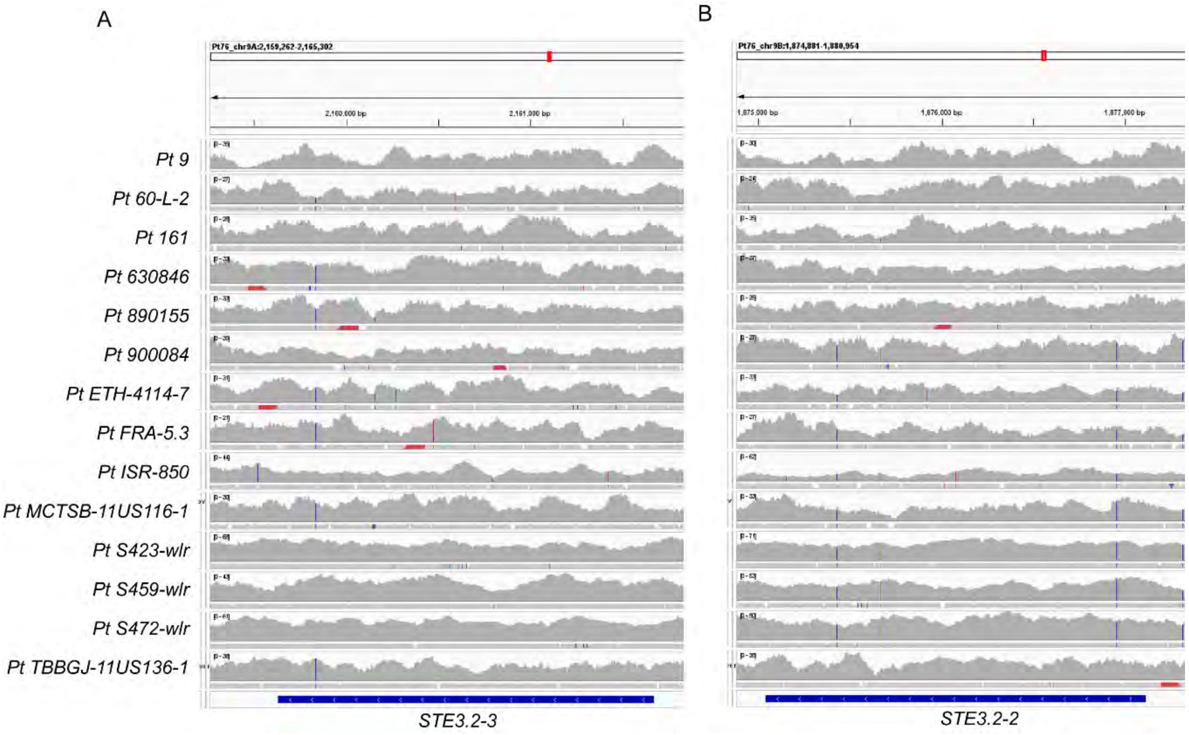

**S24 Fig. IGV screen shots of short read Illumina mapping of various *P. triticina* isolates against the *PR* locus in the *Pt 76* reference.**

(A) shows mapping against the *PR* locus on chromosome 9A and (B) chromosome 9B, respectively.

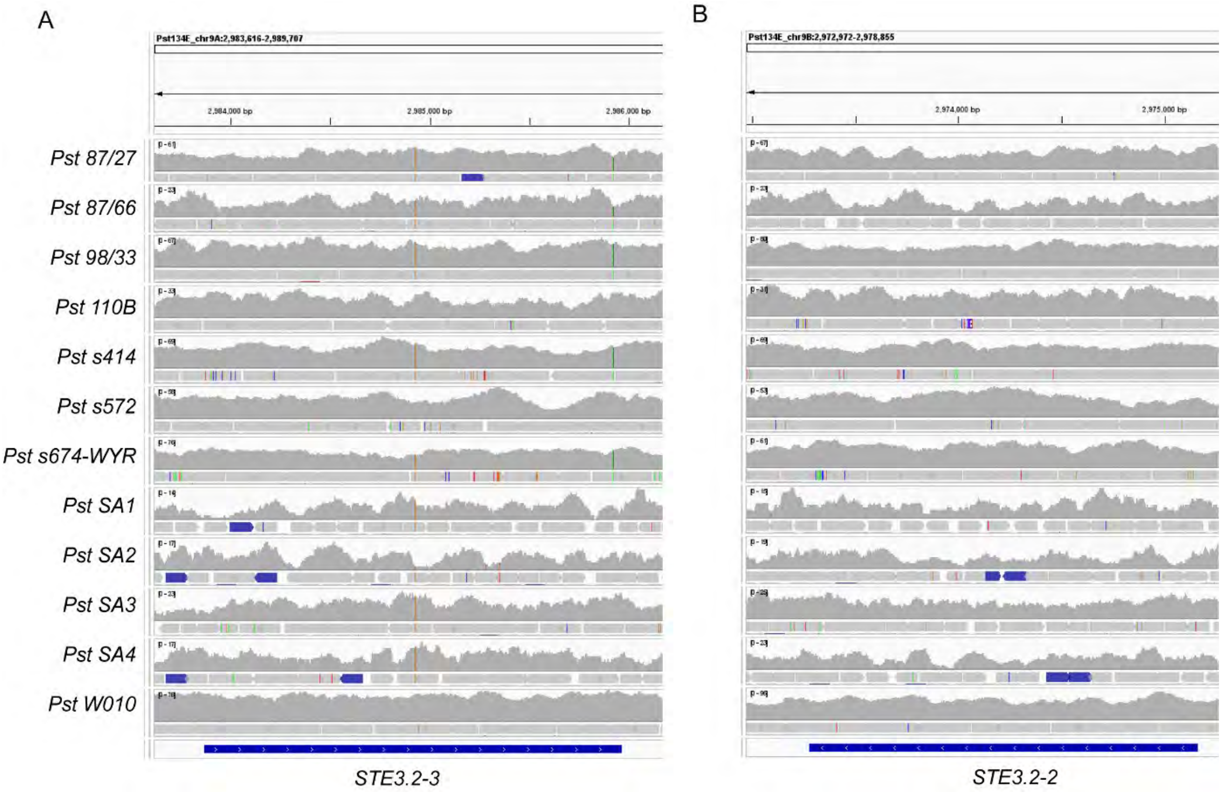


**S25 Fig. IGV screen shots of short read Illumina mapping of various *P. striiformis* f. sp. *tritici* isolates against the *PR* locus in the *Pst 134E* reference.**

(A) shows mapping against the *PR* locus on chromosome 9A and (B) chromosome 9B, respectively.

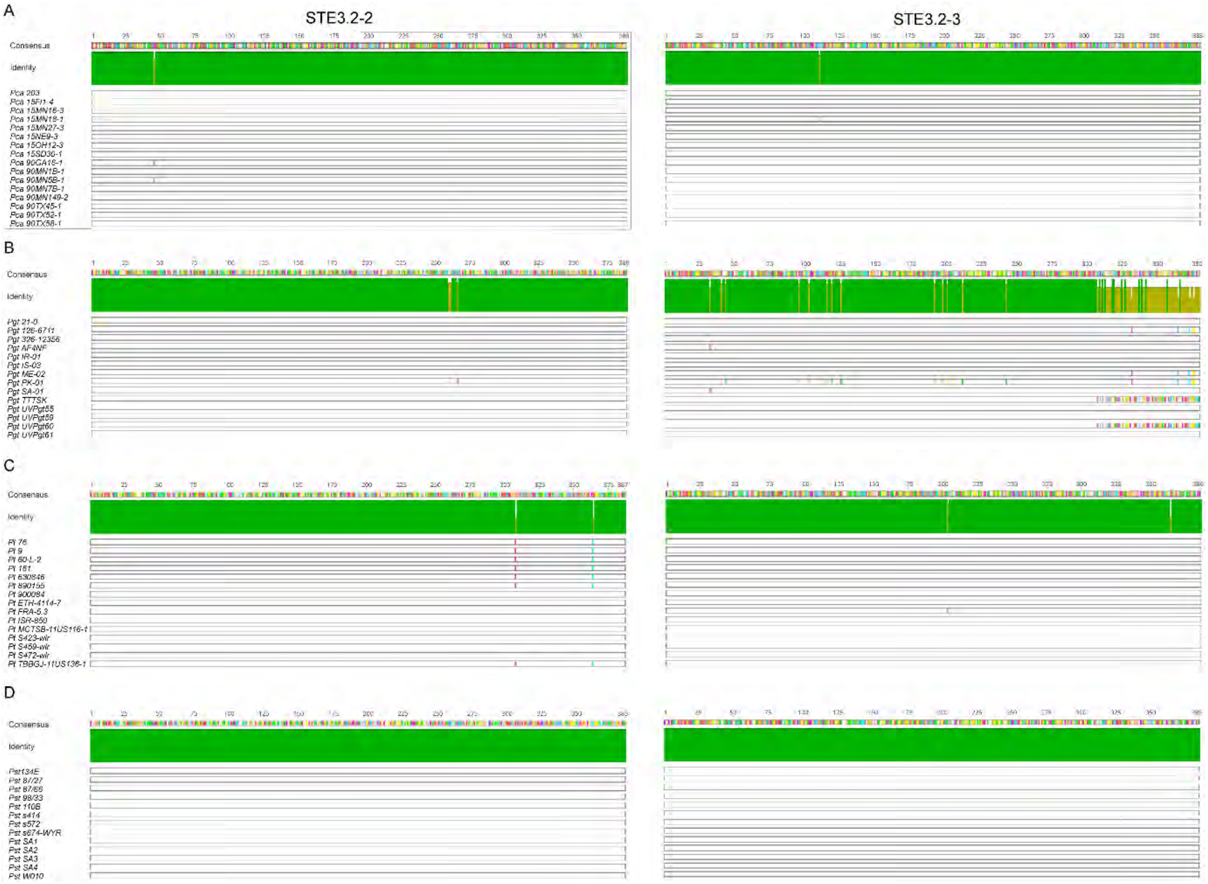


**S26 Fig. Amino acid alignments of STE3.2-2 and STE3.2-3 alleles of four cereal rust fungi.** Multiple sequence alignment of *de novo* reconstructed STE3.2-2 and STE3.2-3 protein sequences. Subfigure contains MSAs; one for STE3.2-2 and one for STE3.2-3. Subfigure A to D show *P. coronata* f. sp. *avenae* ("*Pca*"), *P. graminis* f. sp. *tritici* ("*Pgt*"), *P. triticina* ("*Pt*") and *P. striiformis* f. sp. *tritici* ("*Pst*"), respectively.

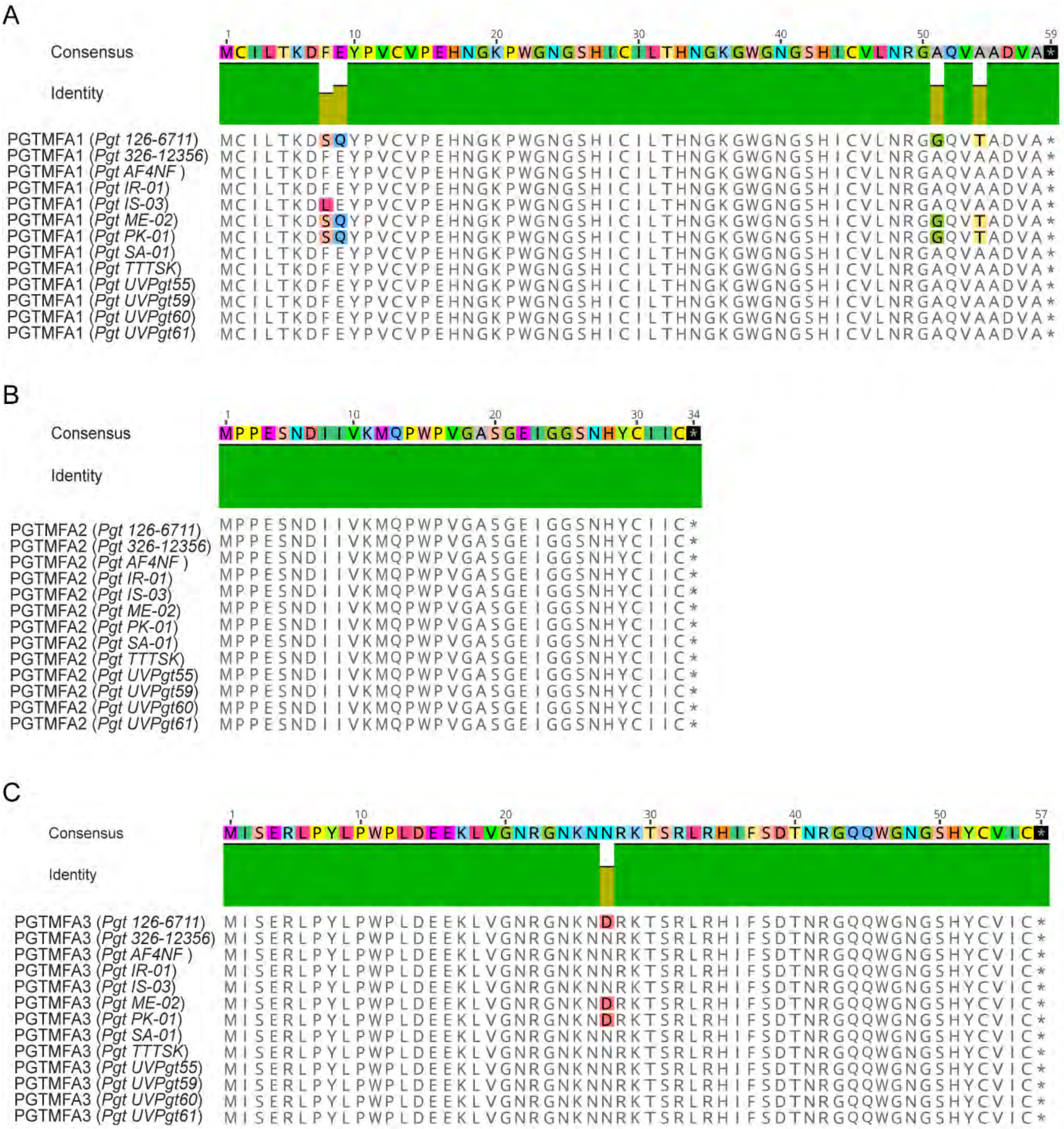


**S27 Fig. Amino acid alignments of the three MFA alleles from various *P. graminis* f. sp. *tritici* isolates.** Amino acid substitutions are highlighted by color.

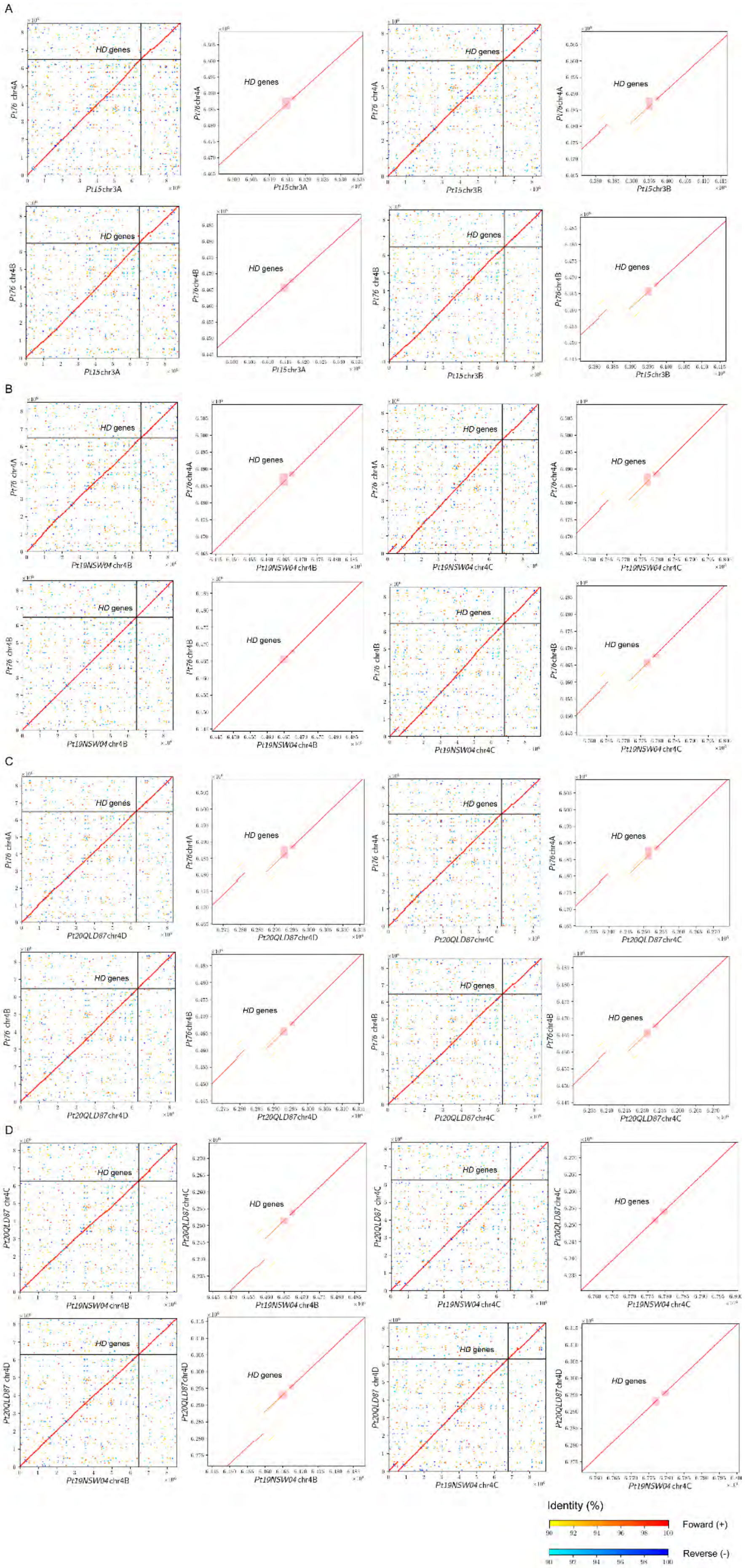
A
B
C
D
HD genes
Pt76 chr4A
Pt76 chr4B
Pt15chr3A
Pt15chr3B
Pt19NSW04 chr4B
Pt19NSW04 chr4C
Pt20QLD87chr4D
Pt20QLD87chr4C
Pt20QLD87 chr4C
Pt20QLD87 chr4D
Identity (%)
Foward (+)
Reverse (-)

**S28 Fig. Whole chromosome alignments of *HD* genes containing chromosomes between haplotypes of four different *P. triticina* isolates.** The figure shows dots plots of whole chromosome alignments of *HD* loci containing chromosomes derived from distinct dikaryotic genomes of four different *P. triticina* isolates including *Pt 15, Pt 19NSW04, Pt 20QLD87* against *Pt 76*. Each panel consists of a dot plot of the whole chromosome and a subset dot plot zooming into the *HD* locus. The *HD* locus is labelled and line colors show the nucleotide percentage identity and nucleotide orientation as indicated in the figure legend. (A) Comparison of nucleotide sequence of chromosome 4s of *Pt 15* and *Pt 76*. (B) Comparison of nucleotide sequence of chromosome 4s of *Pt 19NSW04* and *Pt 76*. (C) Comparison of nucleotide sequence of chromosome 4s of *Pt 20QLD87* and *Pt 76*. (D) Comparison of nucleotide sequence of chromosome 4s of *Pt 19NSW04* and *Pt 20QLD87*.

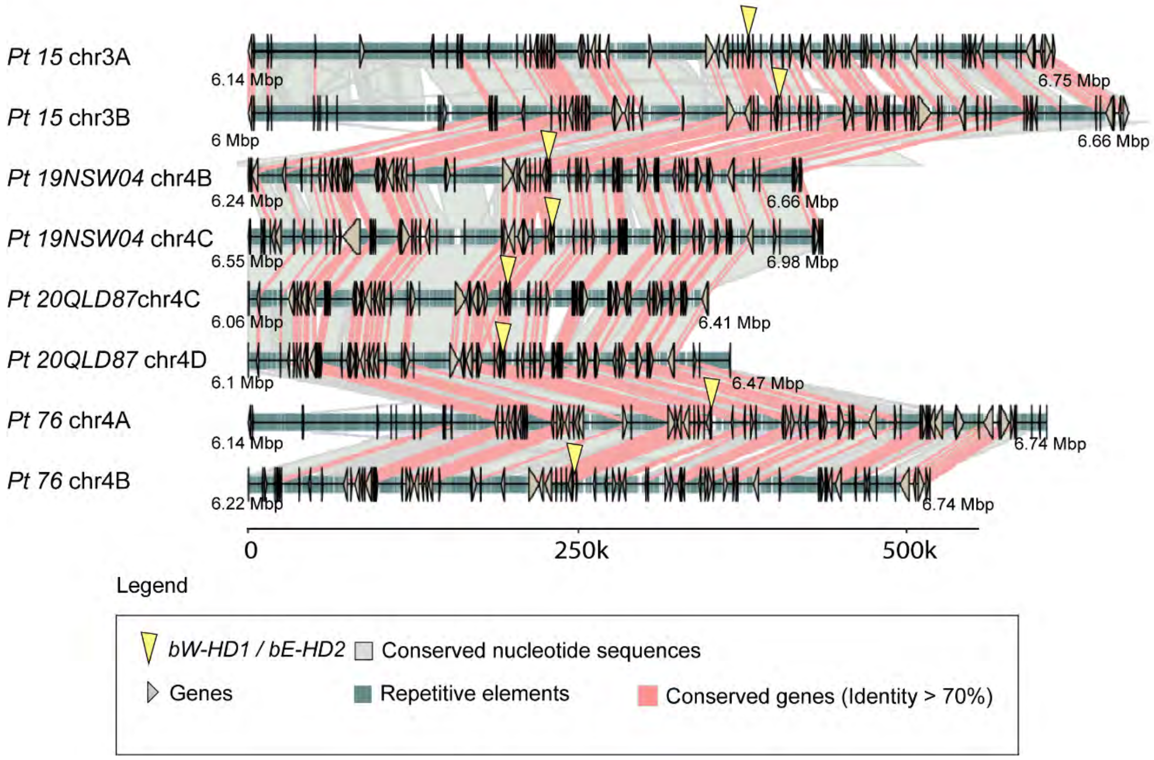


**S29 Fig. The *HD* locus is highly conserved in four *P. triticina* isolates.** Synteny graphs of *HD* loci including proximal regions in the four *P. triticina* isolates *Pt 15, Pt 19NSW04, Pt 20QLD87* and *Pt 76*. Red lines between chromosome sections represent gene pairs with nucleotide sequence identity higher than 70% and grey shades between conserved nucleotide sequences (>=1000 bp and identity >=90%). For additional annotations please refer to the provide legend (“Legend”).

280

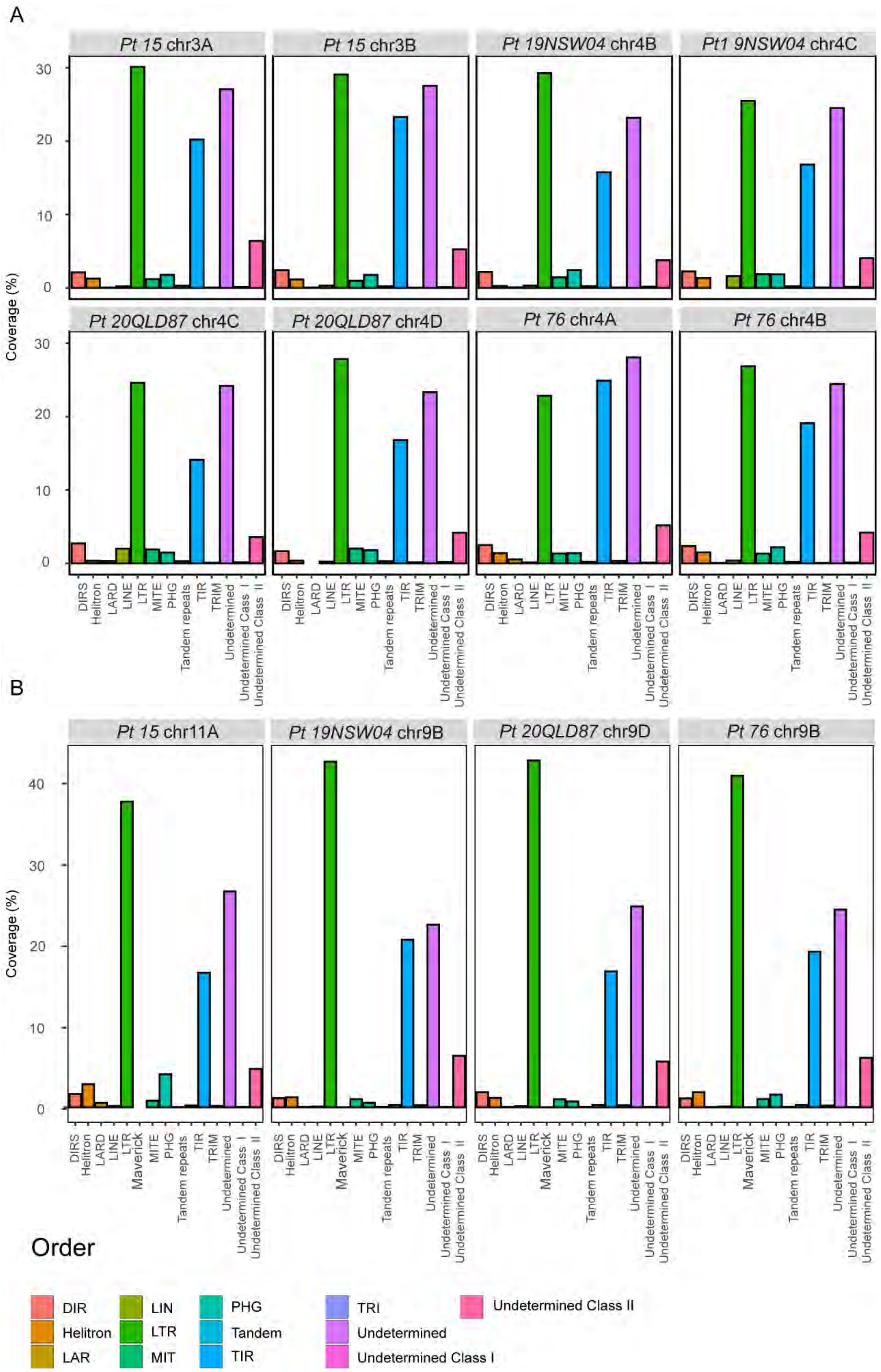
A
Pt 15 chr3A
Pt 15 chr3B
Pt 19NSW04 chr4B
Pt1 9NSW04 chr4C
Pt 20QLD87 chr4C
Pt 20QLD87 chr4D
Pt 76 chr4A
Pt 76 chr4B
Coverage (%)
DIRS
Helitron
LARD
LINE
LTR
MITE
PHG
Tandem repeats
TIR
TRIM
Undetermined
Undetermined Cass I
Undetermined Class II
B
Pt 15 chr11A
Pt 19NSW04 chr9B
Pt 20QLD87 chr9D
Pt 76 chr9B
Maverick
Order
DIR
Helitron
LAR
LIN
LTR
MIT
PHG
Tandem
TIR
TRI
Undetermined
Undetermined Class I
Undetermined Class II

281

**S30 Fig. Coverage of different transposable element orders at the *HD* and *STE3.2-2* locus in four *P. triticina* isolates.** The plots show the percentage of nucleotides covered by different transposable element orders at the (A) *HD* locus and (B) *STE3.2-2* locus. Each subfigure A and B shows the coverage in each haplotype of the dikaryotic genomes of *P. triticina* isolates *Pt 15, Pt 19NSW04, Pt 20QLD87* and *Pt 76*. Different TE orders are color coded as shown in the legend. TEs with no assigned class are labelled "Undetermined". TEs with no assigned order but belonging to Class I (RNA retrotransposons) or Class II (DNA transposons) are labelled "Undetermined Class I" or "Undetermined Class II", respectively.

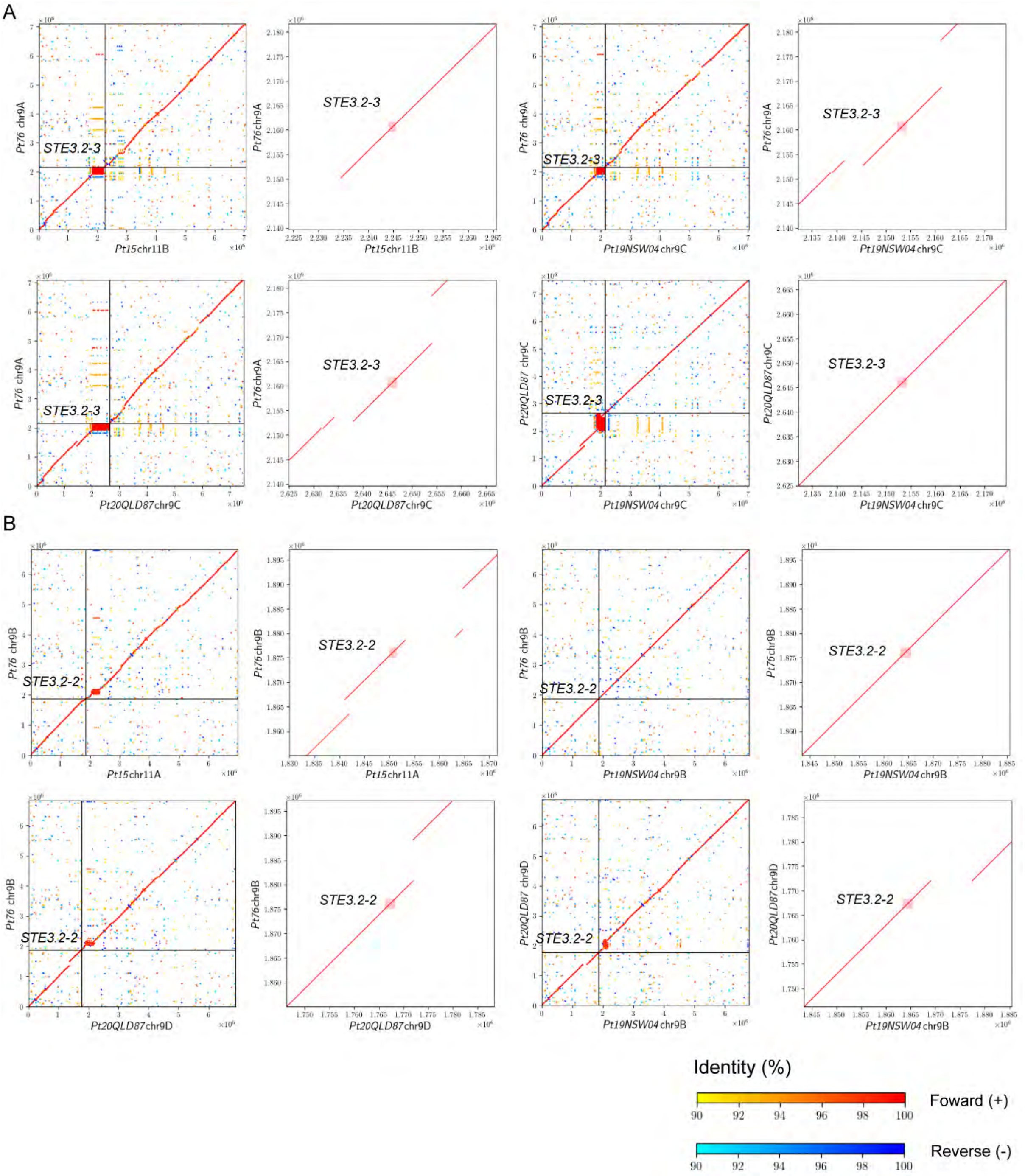


**S31 Fig. Whole chromosome alignments of *Pra* gene containing chromosomes between haplotypes of four different *P. triticina* isolates.** The figure shows dot plots of whole chromosome alignments of *STE3.2-2* or *STE3.2-3* containing chromosomes derived from distinct dikaryotic genomes of four different *P. triticina* isolates including *Pt 15*, *Pt 19NSW04*, *Pt 20QLD87* against *Pt 76*. Each panel consists of a dot plot of the whole chromosome and a subset dot plot zooming into the *STE3.2-2* or *STE3.2-3* locus. The *STE3.2-2* or *STE3.2-* locus is labelled and line colors show the nucleotide percentage identity and nucleotide orientation as indicated in the figure legend. (A) Comparison of nucleotide sequence of chromosome 9s containing *STE3.2-2* loci (B) Comparison of nucleotide sequence of chromosome 9s containing *STE3.2-3*.

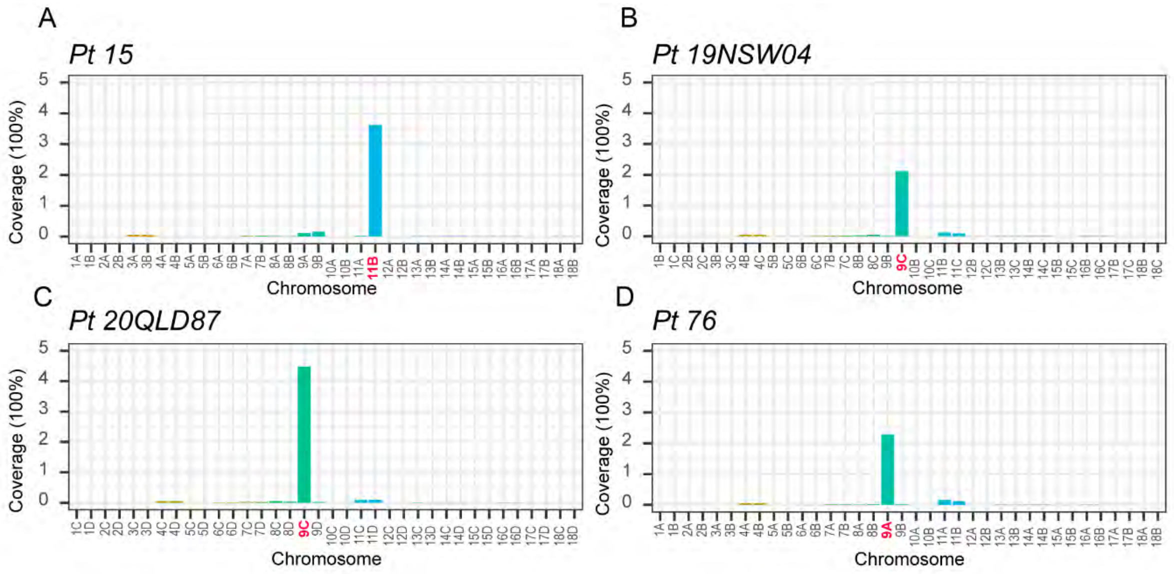


**S32. Nucleotide coverage distribution of Ty3_Pt_*STE3.2-3* on the chromosomes of four *P. triticina* isolates**

The plots show the percentage of nucleotides covered by the transposable element family Ty3_Pt_*STE3.2-3* on each chromosome of four *P. triticina* isolates *Pt 15* (A), *Pt 19NSW04* (B), *Pt 20QLD87* (C) and *Pt 76* (D). Chromosomes carrying *STE3.2-3* are highlighted in red.

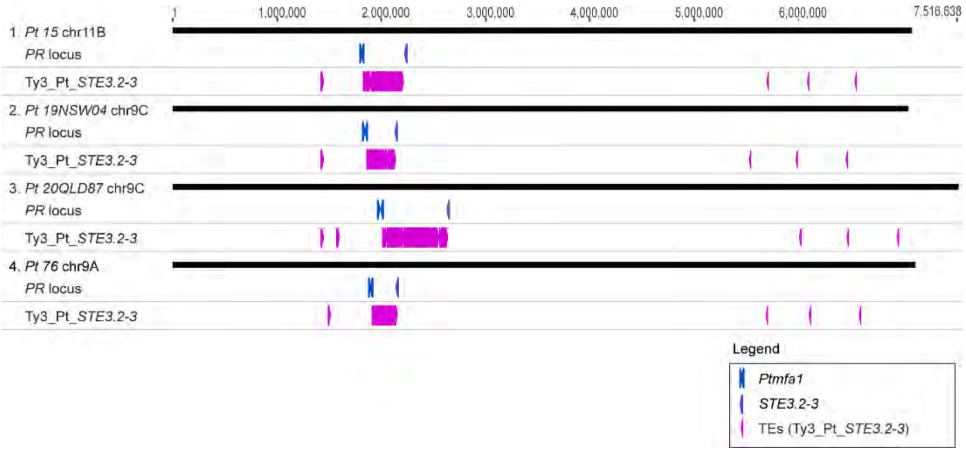

**S33. Schematic illustration of the location of Ty3_Pt_*STE3.2-3* copies at the *STE3.2-3* locus in four *P. triticina* isolates.**

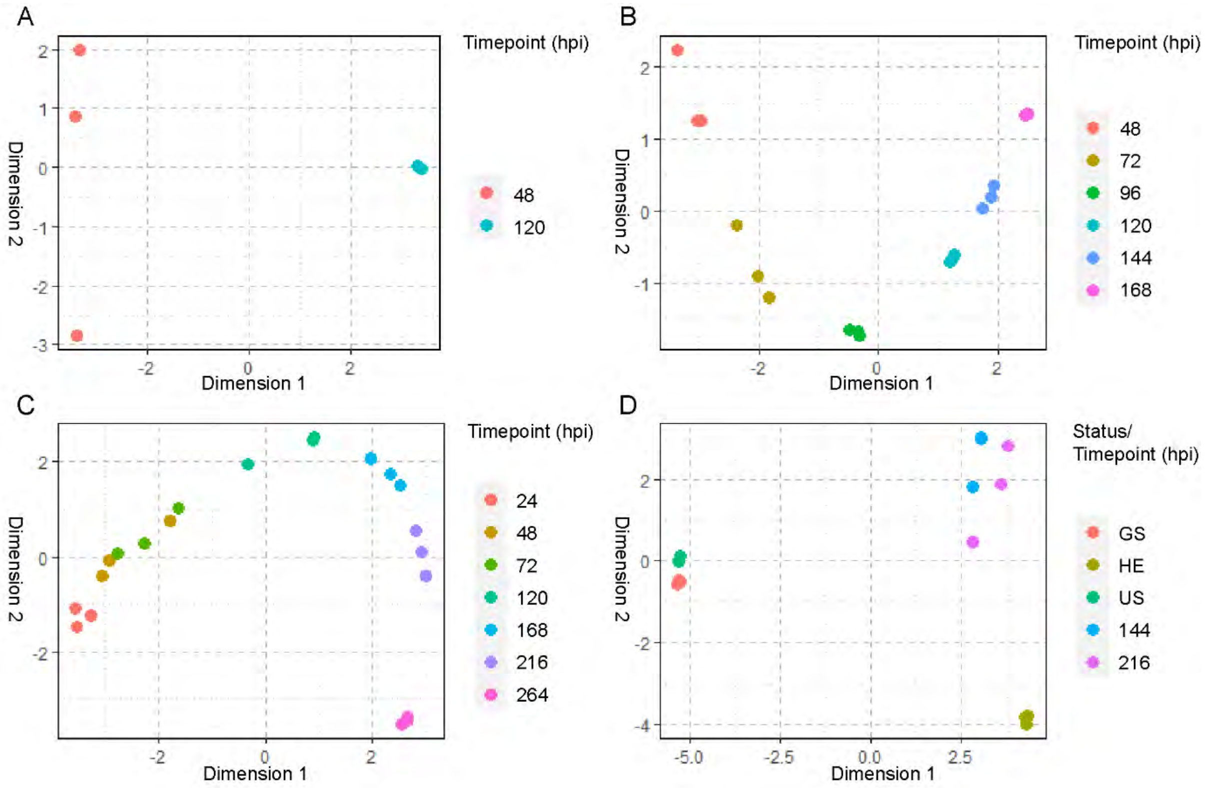


**S34 Fig. Multidimensional scaling (MDS) plot of RNA dataset used in this study.** MDS plots were made with TMM normalized counts for quality control, each dot represents a single sample, and replicates were color coded as indicated in each subfigure ledgend. (A) MDS plot of TMM-normalized value of *Pca* at 48 hour post infection (hpi) and 120 hpi. (B) MDS plot of TMM-normalized value of *Pgt* at 48, 72, 96, 120, 148 and 168 hpi. (C) MDS plot of TMM-normalized value of *Pst* at 24, 48, 72, 120, 168, 216 and 264 hpi. (D) MDS plot of TMM-normalized value of ungerminated spore (US), germinated spores (GS) stages, 144 hpi, 216 hpi and haustoria enriched samples (HE) of *Pst*.

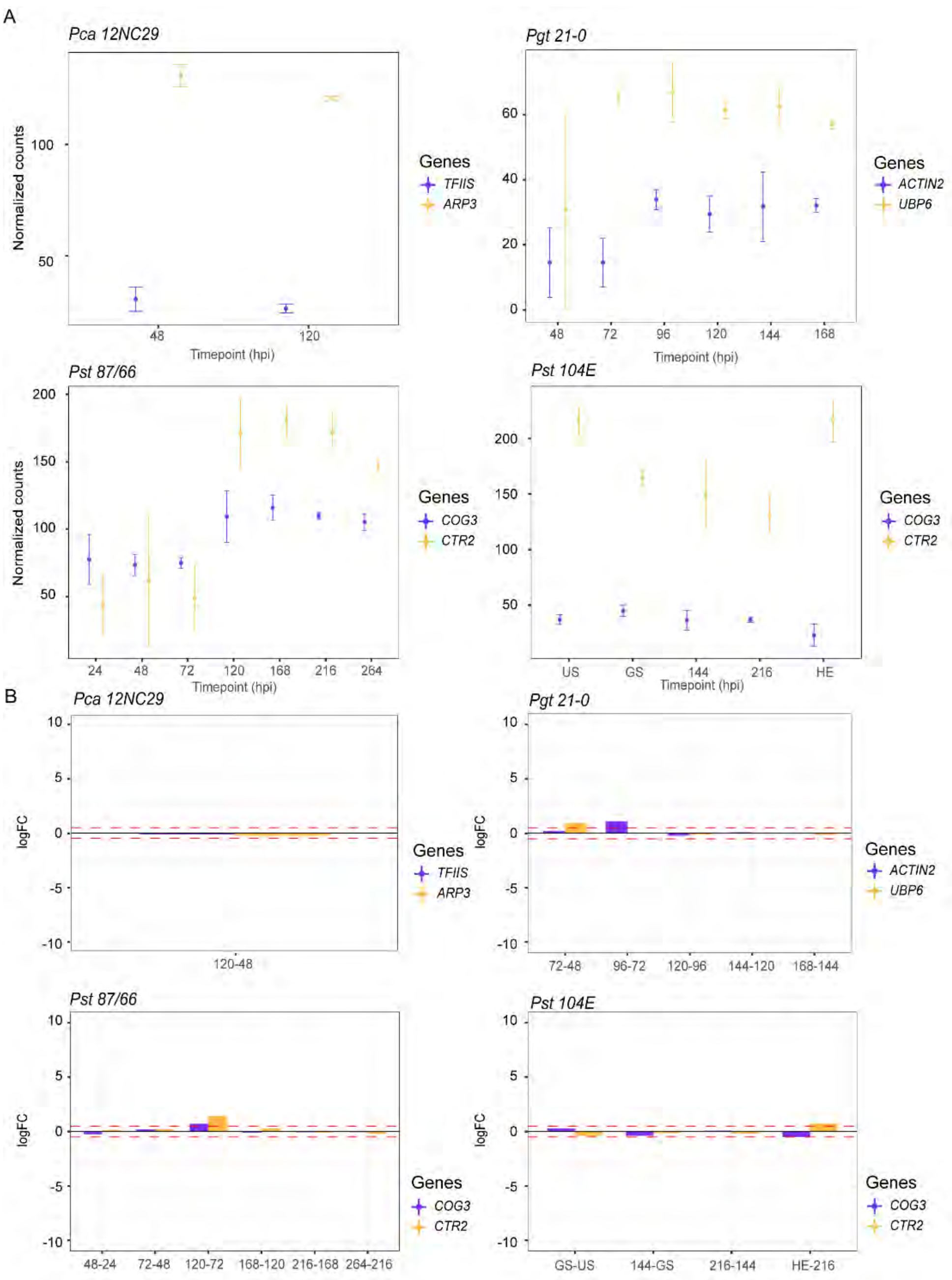


**S35 Fig. Expression of housekeeping genes in four RNAseq datasets.** (A) TMM-normalized value of housekeeping genes in *P. coronata* f. sp. *avenae* (“*Pca 12NC29*”), *P. graminis* f. sp. *tritici* (“*Pgt 21-0*”), and *P. striiformis* f. sp. *tritici* (“*Pst 87/66*” and “*Pst 104E*”). (B) Likelihood ratio test (LRT) method was applied to test significant upregulation of housekeeping genes between timepoints, none of the housekeeping genes show significant upregulation between timepoints. red dashed lines indicate logFC=0.5 and logFC=-0.5 respectively, stars above or below bars indicate statistically significant differences between the two adjacent time points: *$p$<0.05, **$p$<0.01, ***$p$<0.001. Genes are labelled with different colors. Transcription elongation factor TFIIS (*TFIIS*), Actin-related protein 3 (*ARP3*), Actin/actin-like protein 2 (*ACTIN2*), Ubiquitin carboxyl-terminal hydrolase 6 (*UBP6*), Conserved oligomeric Golgi complex subunit 3 (*COG3*), Ctr copper transporter 2 (*CTR2*).

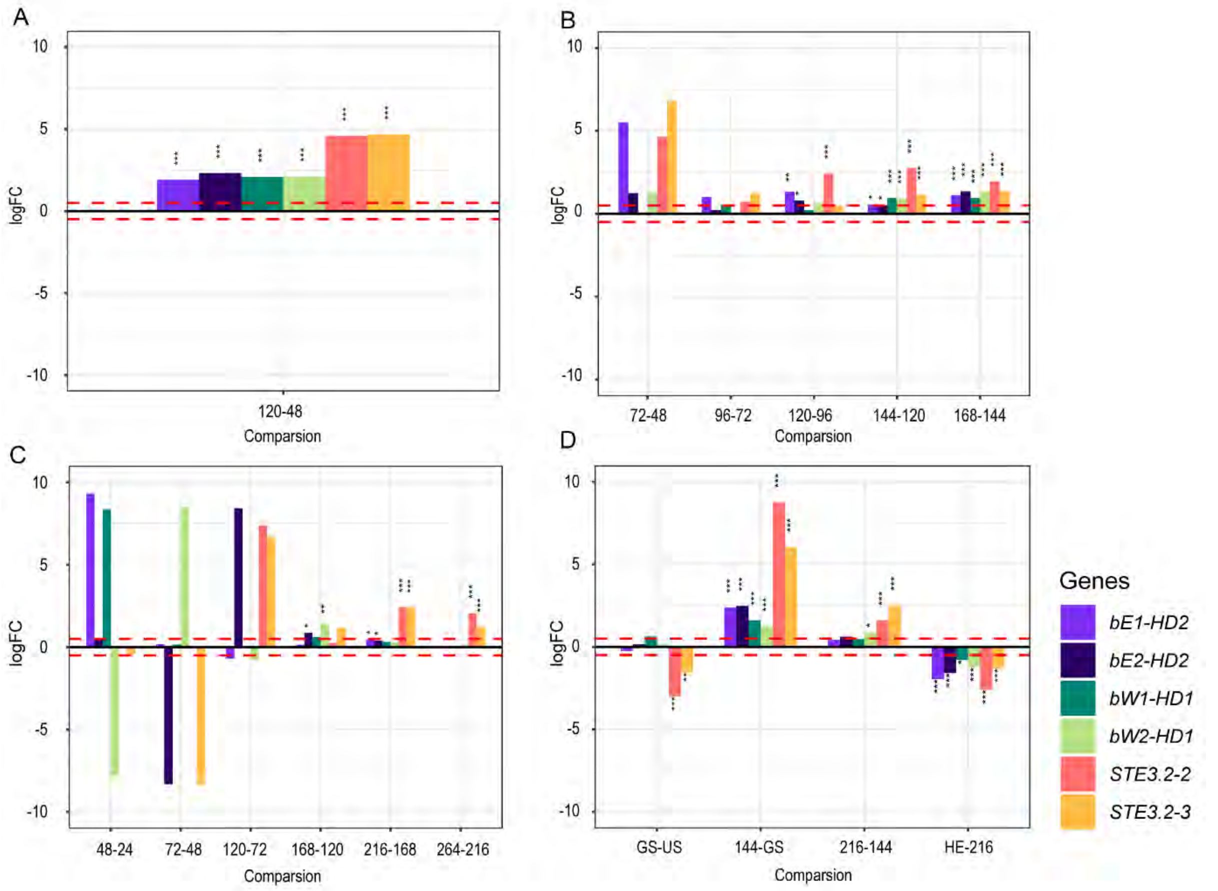


**S36 Fig. Upregulation of *MAT* genes in late stages of the asexual life cycle.** Likelihood ratio test (LRT) method was applied to test significant upregulation of *MAT* genes between timepoints, red dashed lines indicate logFC=0.5 and logFC=-0.5 respectively, stars above or below bars indicate statistically significant differences between the two adjacent time points: *$p$<0.05, **$p$<0.01, ***$p$<0.001. *MAT* genes were labelled with different colors. (A) The expression levels of *MAT* genes in *P. coronata* f. sp. *avenae* ("*Pca 12NC29*") were compared between 120 hours post infection (hpi) and 48 hpi. (B) The expression levels of *MAT* genes in *P. graminis* f. sp. *tritici* ("*Pgt 21-0*") were compared between 72 hpi and 48 hpi, 96 hpi and 72 hpi, 120 hpi and 96 hpi, 144 hpi and 120 hpi, 168 hpi and 144 hpi. (C) The expression levels of *MAT* genes in *P. striiformis* f. sp. *tritici* (*"Pst 87/66")* were compared between 48 hpi and 24 hpi, 72 hpi and 48 hpi, 120 hpi and 72 hpi, 168 hpi and 120 hpi, 216 hpi and 168 hpi, 264 hpi and 216 hpi. (D) The expression levels of *MAT* genes in *P. striiformis* f. sp. *tritici* ("*Pst*

*104E"*) were compared between germinated spores (GS) and ungerminated spores (US), 144 hpi and GS, 216 hpi and 144 hpi, haustoria enriched samples (HE) and 216 hpi.

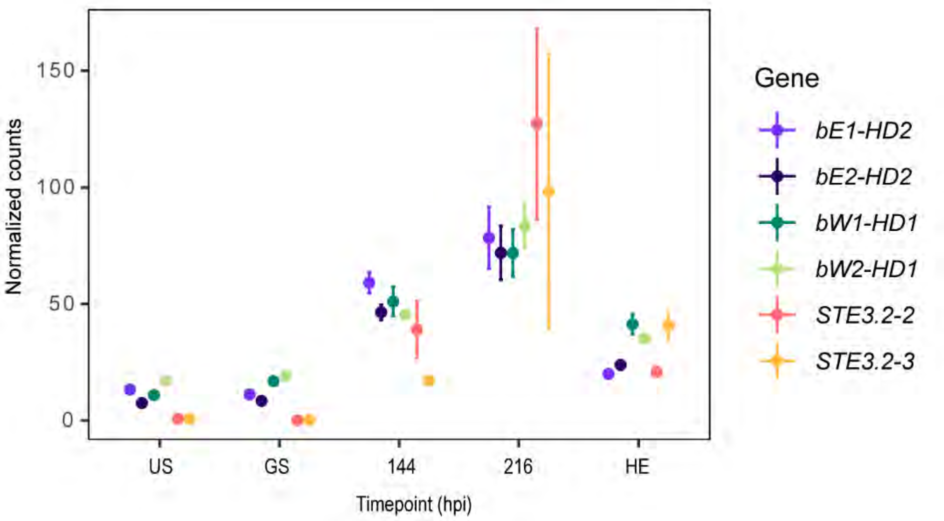


**S37 Fig. *MAT* genes are upregulated in the late asexual infection stage of *P. striiformis* f. sp. *tritici* ("*Pst 104E*").** TMM-normalized value of *MAT* genes in ungerminated spores (US), germinated spores (GS) stages, 144 hpi, 216 hpi and in haustoria enriched samples (HE) of *Pst*. Genes are labelled with different colours.
